# Precision Functional Mapping of Imagined and Experienced Pain

**DOI:** 10.1101/2025.08.06.668906

**Authors:** Michael Sun, Bogdan Petre, Ke Bo, Carmen I. Bango, Malka Hershkop, Sydney L. Shohan, Heejung Jung, Tor D. Wager

**Affiliations:** Dartmouth College; Oregon Health and Science University; Columbia University; Stanford University

## Abstract

Imagination is a principal human capacity, enabling simulation of sensations and situations for planning, learning, and empathy. We examined neural representations of experienced and imagined pain across eight body sites using deep-phenotyping, precision functional MRI with over seven hours of scanning per participant (N = 9). Here we show, using conventional mapping, pattern classification, and representational similarity analyses, that both imagined and experienced pain activated nociceptive regions (e.g., dorsoposterior insula) and non-nociceptive regions (e.g., premotor areas). However, imagined pain did not reliably reactivate body site-selective nociceptive patterns in any region. Instead, imagined and experienced pain shared multivariate representations across widespread cortical and subcortical regions, particularly transmodal cortical areas including dorsomedial and lateral prefrontal cortex, hippocampus, and thalamus. These findings suggest that imagination generates distributed pain-like activity while preserving a neural distinction from experienced pain. This distinction may explain the difficulty of vividly imagining pain and underlie its role in empathy, planning, and pain vulnerability.

---

Imagination allows people to mentally simulate possible futures ^1^, enabling them to savor potential pleasures and develop mastery by anticipating events and their consequences. It allows us to refine our performance through mental practice: Envisioning the thrill of hoisting a championship trophy or the pain of touching a burning ember can serve as reinforcers that shape decision-making even in the absence of physical action^2–5^. Yet, imagination can be a double-edged sword. Imagining pain helps individuals prepare for medical procedures^6–9^, increase negative expectations, and produce aversive learning, components of placebo and nocebo effects in medical contexts ^11–14^. Imagery is also central to how we understand and empathize with others^15^, and is a crucial factor in cognitive and social development^16^. In clinical settings, this capacity is harnessed through imaginal exposure therapies, which activate neural circuits involved in extinction learning ^17^ —a core mechanism underlying symptom reduction in posttraumatic stress disorder ^18^, anxiety disorders (e.g., specific phobia and obsessive-compulsive disorder)^19^, and, increasingly, in chronic pain^20,21^. This raises some central questions: How does the brain represent imagined pain, a stimulus that must be veridical enough to guide behavior, yet not confused with the real thing? Does imagination activate early nociceptive representations, or are its effects instead mediated by higher-order cortical areas? Can the brain walk the fine line of engaging somatic and affective representations without fully triggering the alarm systems reserved for actual injury?

Imagination has been a central topic in neuroscience. Early neuroimaging work in vision helped resolve a classic debate over whether or not imagination is depictive, resembling perception and involving sensory reactivation (e.g., visualizing an object reactivates early visual cortex)^22^, or propositional, whereby imagination encodes abstract, language-like descriptions rather than perceptual details^23^. More recently, studies have used multivariate pattern analyses to demonstrate re-activation of content-selective patterns in early perceptual cortices in vision^24–28^, olfaction^29,30^, taste^31^, and audition^32–39^, including activation of somatotopic regions in somatosensory cortices^40^. These findings suggest that imagery may activate early sensory representations. At the same time, early sensory activation is typically weaker^28,41–44^, and some recent studies have highlighted distinct, opposing activity patterns underlying visual perception and imagery^45^. The degree to which imagination can activate sensory representations, the conditions under which it does so, and alternative cortical substrates for imagination remain open questions.

The effects of imagination on pain are even less well understood. Pain is particularly crucial for both empathy ^46–50^ and personal decision-making and learning^51^, and it involves distinct pathways^52^, representations^53,54^ and multivariate fMRI patterns ^55,56^ from other forms of somatosensation. On one hand, a seminal study identified little activation in pain-related regions with imagination, with greater activation in the absence of stimulation after hypnotic suggestion ^57^. This finding suggests that pain is particularly difficult to volitionally imagine in a veridical fashion, which perhaps underlies the so-called “empathy gap” in pain ^58^. However, other studies have found overlapping activations in pain-related regions, including the anterior cingulate cortex (ACC), insula, and primary and secondary somatosensory cortices (S1, S2)^15,46,57,59^, as well as subcortical areas like the thalamus, amygdala, and parahippocampus ^60,61^. Prefrontal and parietal cortices have also been found to be activated during pain imagery, providing a potential substrate for imagery generation ^15,46,57,59^.

A central issue is that all of these areas are activated by many processes beyond pain ^55^, and may thus reflect changes in a variety of cognitive and affective processes, including language and emotion. The presence of overlapping activation does not imply shared representations of pain per se^62^. How deeply do top-down processes such as mental imagery access the sensory hierarchy—specifically, can they reinstate individualized, somatotopically organized nociceptive representations, or are they constrained to higher-order pain-related representations? To identify common engagement of pain representations, shared activity patterns should both reliably encode pain and be selective for pain^62^. Previous studies of pain imagery have not identified representations with these properties. For example, Derbyshire et al.^57^ do not quantify the relative magnitude of activation or representational pattern similarity for experienced and imagined pain, nor do they identify pain intensity-encoding nociceptive representations. Thus, how strongly imagination affects nociceptive representations, and what types of representations are shared across pain imagery and veridical pain experience, remains unknown.

We address this question using precision functional mapping (PFM). This approach leverages high-resolution, high-volume individual-level fMRI data to reveal reliable and anatomically specific brain representations that are often obscured in group-averaged analyses. Unlike conventional group-averaged analyses, which often blur fine-grained functional topographies due to inter-individual variability, precision functional mapping (PFM) enables the discovery of individualized neural signatures with high reliability and reproducibility ^63,64^, despite lower total group sample size^65^. PFM studies typically involve the study of “highly sampled individuals”, collecting several hours of data per participant, allowing for the identification of distinct functional boundaries, networks, and representational patterns that are consistent across sessions and tasks. We analyzed 7+ hours of fMRI data during imagined and perceived heat pain on 8 distinct body sites (Figure 1), allowing us to identify body site selective nociceptive representations in individual participants, and to test both imagined and perceived pain and their body site selectivity on new, independent scan days. This allowed us to test whether imagination activates individualized nociceptive representations in key areas targeted by ascending nociceptive pathways, including spinothalamic pathway ventral posterior lateral thalamus (VPL) and dorsal posterior insula (dpIns), medial pathway intralaminar thalamus and anterior midcingulate (aMCC).

**Figure 1:**
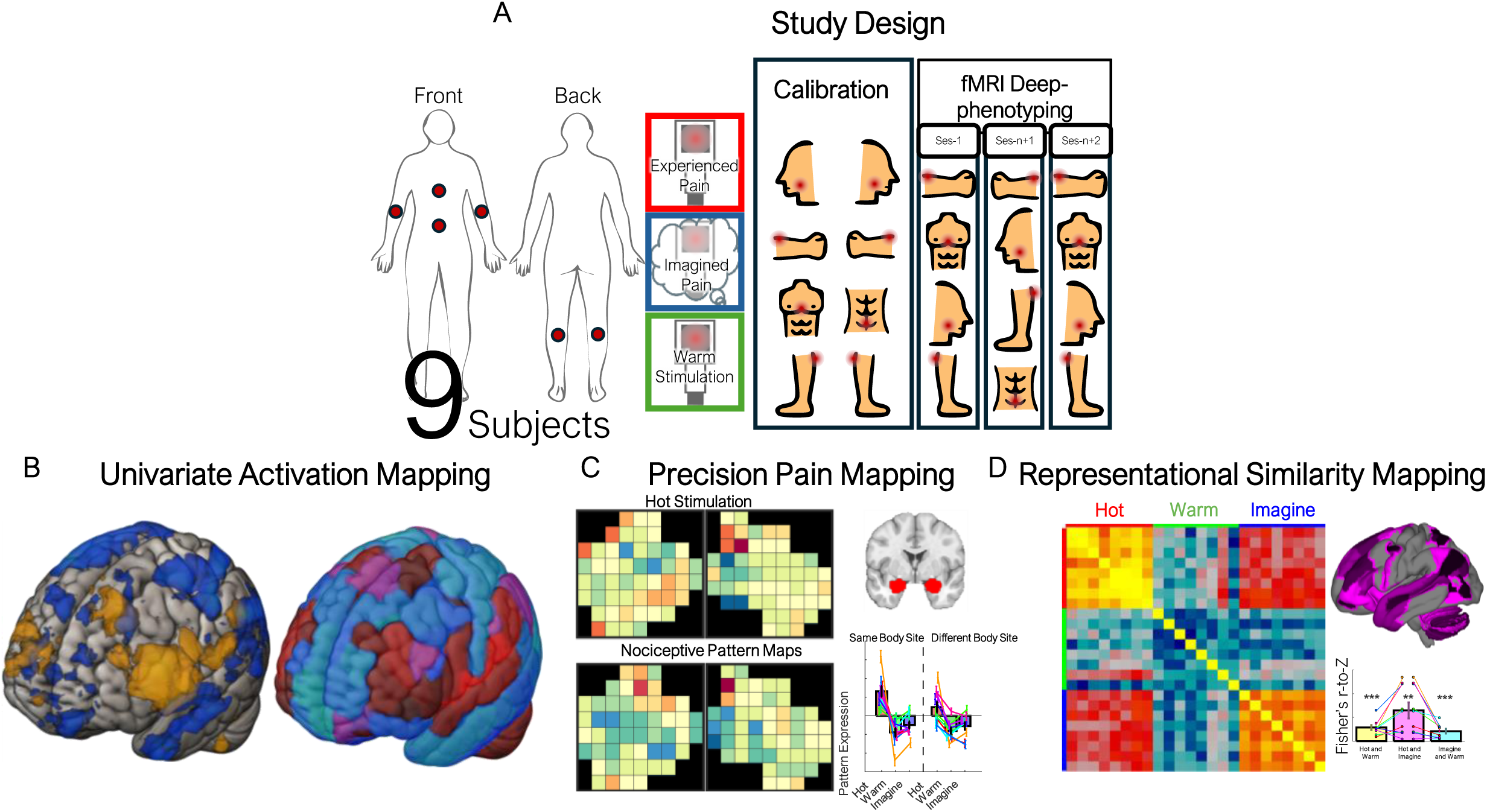
Study design and analytical approaches. (A) A within-participant experimental design had participants calibrate painful and non-painful stimulation temperatures at each of 8 body sites (chest, abdomen, and bilateral lower jaw, arms, and legs). In the fMRI scanner, participants underwent ten sessions of fMRI scanning, featuring pseudo-randomized trials of hot or warm stimulation, or instructions to imagine themselves receiving painful heat stimulation. FMRI data are analyzed through the lens of (B) Univariate Activation Maps, (C) Precision Classification Pattern Tests, and (D) Representational Similarity Analyses. (B-D) show group results from each analysis, elaborated further in the text. Study timeline and scan days can be found in Supplemental Figure 1. In (C) and (D), data are presented as mean ± SEM across n = 9 participants.

PFM has been most commonly applied to resting state functional connectivity, revealing individualized network organization. However, resting-state connectivity does not capture how specific cognitive or sensory states are instantiated within these networks. Task-based PFM, by contrast, enables the identification of individualized, state-dependent multivoxel representations. For example, Gordon et al., 2023 discovered three intereffector regions (superior, medial, inferior) across primary somatomotor cortices, that support an Action Mode Network/Somato-cognitive Action Network, whereby the body is prepared to synchronize to act in response to a demand. These sites are functionally connected with SMA, thalamus, posterior putamen, postural cerebellum, dorsal anterior cingulate, parietal regions, and the insula. Building on this logic, we apply task-based PFM to test whether imagined pain re-engages individualized nociceptive representations, or instead recruits distributed pain-like patterns in non-nociceptive cortex. This approach is uniquely suited to addressing questions about the representational limits of imagination.

To identify other brain systems in which pain and imagined pain involve similar representations, we used Representational Similarity Analysis (RSA) in a series of regions spanning the brain and compared the distribution of findings to established maps of cortical hierarchies^66^. This allowed us to identify ‘pain-like’ representations across higher-level cortical systems. These approaches allowed us to identify and localize substrates for imagined pain in the human brain.

## Results

Prior to neuroimaging, each participant underwent a calibration session using an Ascending Method of Limits procedure^67^ to determine the minimum temperature discriminable from baseline (32°C; thermal detection threshold), maximum tolerable stimulus intensity (pain tolerance), and mapping between temperature and pain self-report on each body site (Figure 2 and Methods). The pain tolerance values for each body site within-participant were used for “Hot” stimulation in fMRI, and ranged from 46-48.5°C; see Methods). Thermal detection threshold for each body site plus 1°C was used for “Warm” (34.5-39.5 degrees) in fMRI. Thermal detection thresholds varied significantly by body site, F(7, 14.93) = 5.29, p = .003, partial ² = .71. Post-FDR^68^ correction (q < .05), the left arm had significantly lower thresholds (greater sensitivity) than the face (vs. left face q < .007; vs. right face q < .001), legs (vs. left leg: q < .001; vs. right leg: q = .002), chest (q = .001), abdomen (q = .03), and right leg (q = .002). The right arm also showed lower thresholds than the face (vs. left face q < .013; marginal vs. right face q < .060), legs (vs. left leg: q < .002; marginal vs. right leg: q < .071), and chest (q = .016). Finally, the abdomen showed significantly lower thresholds than the left leg (q = .034). Pain tolerance also significantly varied by body site, F(7, 12.619) = 13.67, p < .001, as both left face (vs. left arm: q < .001; vs. right arm: q = .002; vs. left leg: q = .006; vs. right leg: q < .001; vs. chest: q = .018; vs. abdomen: q < .001) and right face (vs. left arm: q < .001; vs. right arm: q = .002; vs. left leg: q = .003; vs. right leg: q < .001; vs. chest: q = .004; vs. abdomen: q < .001) showed higher pain tolerances than all other body sites. Body sites did not vary significantly in pain ratings, Intensity: F(7, 13.23) = 1.45, p = .266, Valence: F(7, 15.09) = 1.45, p = .258.

**Figure 2:**
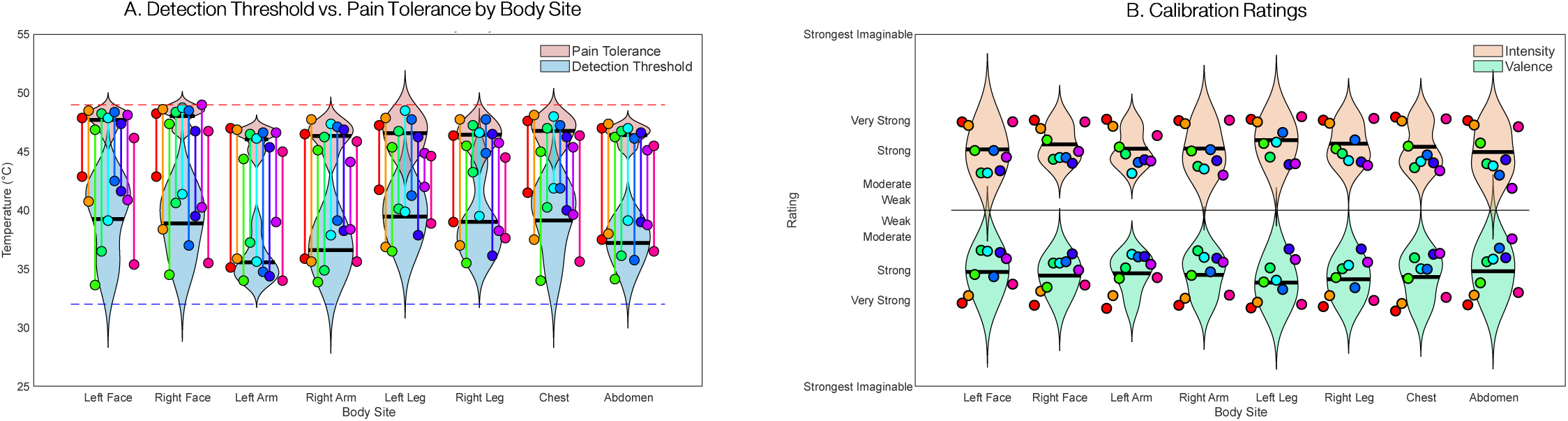
Behavioral and Self-Report Results. (A) Average detection and pain tolerance temperatures for each body site and participant (dots with connecting lines). Baseline temperature was 32°C (blue dashed line), and the maximum temperature delivered was 49°C (red line). (B) Pain valence and intensity in response to high heat (tolerance intensity) during calibration. No significant differences in pain metrics were found between body sites post-calibration. Participant-level data is available in Supplementary Figure 2-3 and body site-level data can be found in Supplementary Table S1.

### Group-level and individual participant activation maps

To evaluate whether experienced and imagined pain engage similar neural systems at the level of univariate activation—consistent with prior group-level pain-mapping studies—we first examined group-level and individual contrasts for Experienced pain (Hot–Warm) and Imagined pain (Imagine–Warm; all FDR q < 0.05 corrected for multiple comparisons; Figure 3, Supplementary Figures 4 and 5, Methods, and Supplementary Table S2).

**Figure 3:**
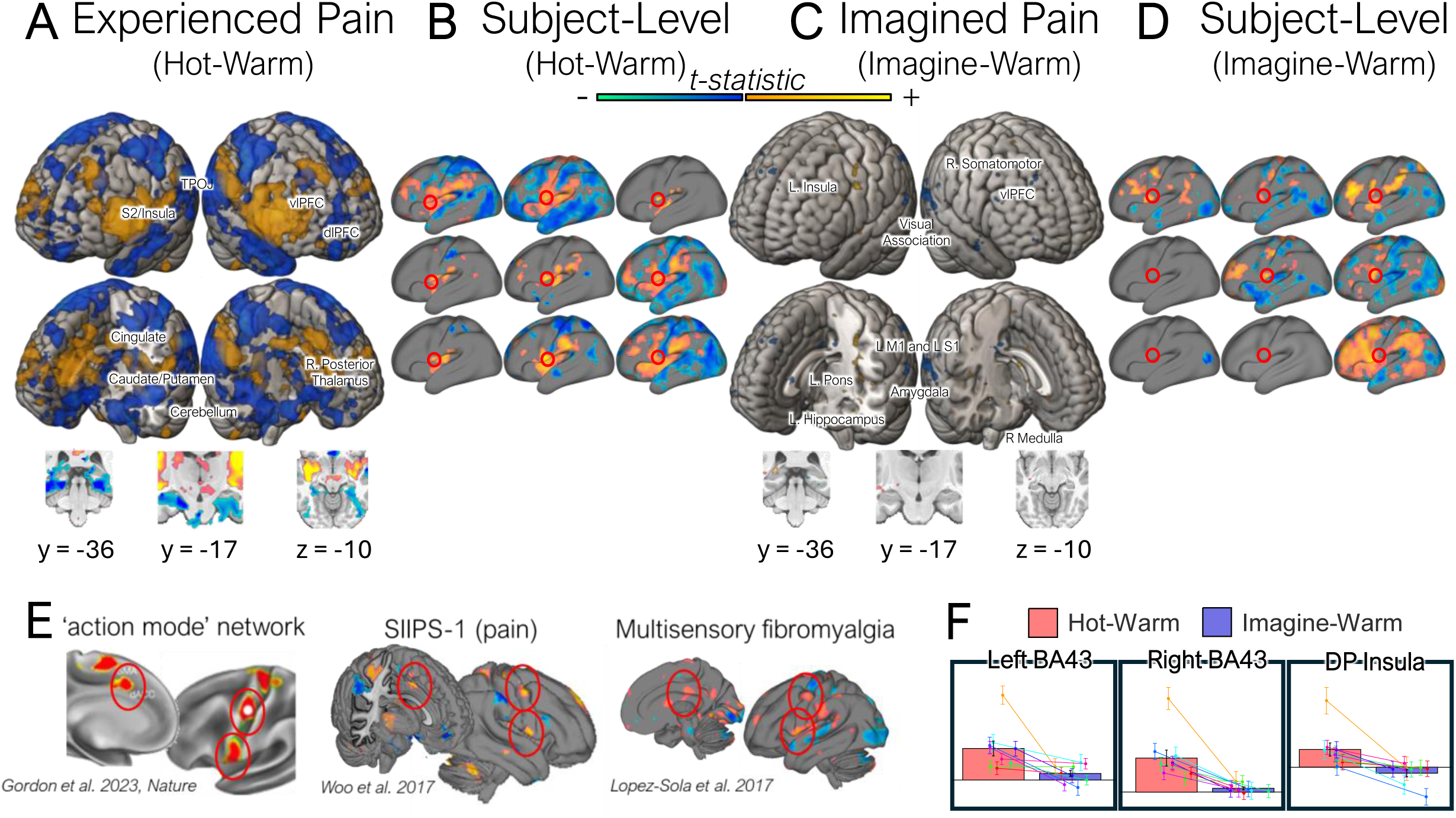
Group-level and participant-level univariate contrast results for experienced pain (Hot–Warm) and imagined pain (Imagine–Warm). Shown on top are group-level and participant-level t-maps (yellow = activation, blue = deactivation). (A) Experienced pain activated S2, insula, operculum, paracentral lobule, premotor, thalamus, vlPFC, dlPFC, cingulate, parietal inferior lobule, TPOJ, medial temporal, A1, AAC, V1, V8, midbrain, ventral striatum, medulla, putamen, globus pallidus, and caudate. (C) Imagined pain activated hippocampus, amygdala, insula, operculum, paracentral lobule, premotor, vlPFC, lateral temporal, midbrain, medulla, and pons. (B, D) Participant-level maps reveal consistent activation in an inferior intereffector region across all participants during experienced pain, and in 6 of 9 participants during imagined pain. This region overlaps with the ‘action mode’ network identified by Gordon et al. (2023)^69^ (E), and is also evident in previously published multivoxel pain signatures including SIIPS-1^72^ and the Multisensory Fibromyalgia signature^73^. (F) Bar plots show individual and group mean beta values in select regions. S1: primary somatosensory cortex; S2: secondary somatosensory cortex; vlPFC: ventrolateral prefrontal cortex; dlPFC: dorsolateral prefrontal cortex; TPOJ: temporoparietal occipital junction; A1: primary auditory cortex; AAC: auditory association cortex; V1: primary visual cortex; V8: eight visual area, visual ventral cortex. In (F), bars show group mean ± SEM across n = 9 participants; dots show individual participants.

Group analyses indicated significant activation in a broad set of typical pain-processing regions (Figure 3A). Many of these were activated in multiple individual participants as well (Figure 3B). Every individual participant showed pain-evoked responses (Hot - Warm) in core insular-opercular regions—spanning somatomotor and parietal operculum (right OP2, left OP4; FOP1-BA43, FOP3, FOP4), posterior and middle insular cortex (PoI1-2; MI), and surrounding association cortex (left TA2; left PFop). The somatomotor region BA43 lies at the ventral aspect of the motor cortex adjacent to M1 (BA4) and overlaps with the inferior ‘inter-effector’ region in Gordon et al.^69^. It is also the inter-effector region most closely connected with the mid-insula and posterior insula, which are selectively engaged in thermal pain^70,71^, and surrounding operculum. A wider set of regions was activated in at least 22.22% (2 of 9) of individual participants, including anterior mid-cingulate cortex, medial and lateral thalamus, dorsolateral and ventrolateral prefrontal cortices (dlPFC, vlPFC), and other subcortical forebrain cerebellar, and brainstem regions (Supplementary Figure 4 and Supplementary Table S2).

Imagined pain (Imagine - Warm) activated a much more limited set of regions at the group level, with stronger evidence for inter-individual variability. Group findings include ten regions activated in common with experienced pain (Figure 3, right), including left midbrain (dorsal lateral reticular formation), left posterior insula (PoI1, BA52), left anterior insula (piriform), left rostral insula, left premotor cortex (rostral BA6), the right inter-effector region BA43, right dACC, and left vlPFC (BA44, anterior inferior frontal junction). Interestingly, S1 (right BA2, bilateral BA3a, BA4), left ventral and dorsal BA6, right medial posterior BA6, lateral temporal cortex (dorsal and ventral temporal gyrus, posterior inferior temporal cortex), parahippocampal transition area, hippocampal formation (left subiculum, and bilateral CA23), left/right basolateral amygdala, left pons, and bilateral medulla were activated during imagined, but not experienced pain. In individual participants, the right inter-effector region BA43 (4/9) was the most consistently activated region that was shared between imagined and experienced pain. Outside of these regions, we found consistent activation in left rostral and ventral BA6 (5/9 participants), right medial posterior BA6 (4/9), and left BA4 in M1 (4/9). Left retroinsular cortex, left piriform cortex, left ventral temporal gyrus, left subiculum, right basolateral amygdala, and bilateral medulla, were not significantly active in any individual participant(Supplementary Figure 5 and Supplementary Table 2).

We also conducted conjunction analyses across experienced and imagined pain in individual participants (Supplemental Figure S6 and Supplementary Table S2). The most consistent common activation among participants was found in the ventral inter-effector regions (left BA43: 6/9; right BA43: 4/9). Other regions activated in both experienced and imagined pain within individuals included other somatomotor regions (bilateral rostral BA6, right FOP1, right SMA), insula (right middle, FOP3), left inferior parietal lobule (PFop), left dlPFC (BA46), and the right anterior putamen(Supplementary Figure 6). Thus, different participants tended to show FDR-corrected overlap in different regions. Notably, both experienced and imagined pain consistently engaged a focal region within the inferior intereffector zone of the primary somatomotor cortex, a region previously implicated in action-mode and somato-cognitive integration, despite the absence of nociceptive input during imagination.

### Reliability of imagined pain representations across time

Next, to assess consistency of imagined pain representations over extended task performance, we quantified within-subject reliability of multivariate activation patterns across runs and across days. Cross-session correlations of imagined-pain pattern expression revealed significant within-subject stability (mean r = 0.161; t(8) = 4.92, p = 5.8 × 10□⁴), exceeding a permutation-based null distribution generated by shuffling session labels within subject. A conservative split-half analysis (odd vs. even runs across sessions) similarly demonstrated significant reproducibility (mean r = 0.097; t(8) = 4.46, p = 0.001). Reliability was body-site selective, with higher similarity for the same imagined body site than for different sites across days, indicating consistent, content-specific engagement.

### Analyses of body site selective nociceptive representations

To test whether experienced pain gives rise to individualized, body site–specific nociceptive representations, and whether such representations are recapitulated during pain imagination, we trained participant-specific support vector machine (SVM) classifiers within key predefined pain-related regions (see Figure 5). For each region, we trained participant-specific support vector machine (SVM) classifiers for each region to distinguish hot from warm stimulation within each body site, which constituted potential nociceptive representations. We surmised that if there were nociceptive representations that were body site specific in each of these areas, such representations would be recapitulated during pain imagination. To cross-validate our results, we tested effects of interest on data from independent sessions collected on different days (leave-one-session-out cross-validation; Figure 4, Methods, and Supplementary Table S3). Nociceptive representations were defined as neural patterns with significant effects in test data for (1) [Hot - Rest] on the target body site, (2) [Hot - Warm] for the target body site, and (3) [target body site > other body sites] in the [Hot - Warm] contrast, to identify body-site selectivity. Nociceptive representations identified in this way were tested for [Imagine - Rest], [Imagine - Warm], and [Imagine - Warm] x [Target vs. Other body site] effects. For each putative nociceptive representation, we calculated the pattern expression on test data for each condition (Hot, Warm, Imagine; see Methods) and contrast across pattern expression scores (e.g., [Imagine - Warm]). We then tested these scores against zero at the group level to assess statistical significance treating participants as a random effect.

**Figure 4:**
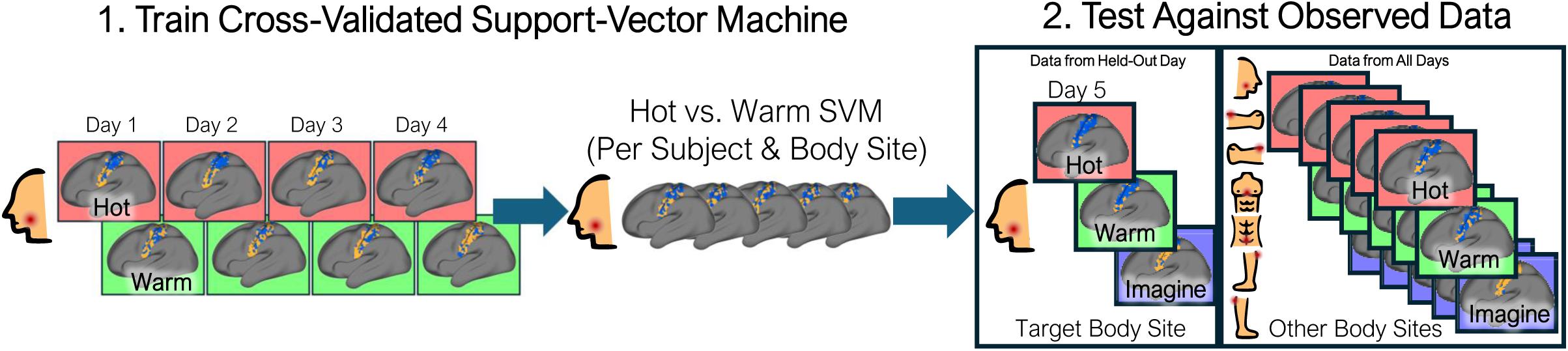
Precision functional mapping of nociceptive representations using cross- validated SVMs. Analysis pipeline for subject- and body site–specific precision functional mapping. Within each pain-processing region of interest, Hot vs. Warm conditions are discriminated using a leave-one-session (day)-out cross-validated support vector machine (SVM). Resulting participant- and body site–specific SVM weight maps are tested on held-out Hot vs. Warm and Hot vs. Rest data, as well as on Hot, Warm, and Imagine conditions from non-target body sites across all sessions. This approach assesses the specificity and generalization of nociceptive representations within individuals. Subject-level SVM weight maps and example pattern-expression results are shown in Figure 9.

**Figure 5:**
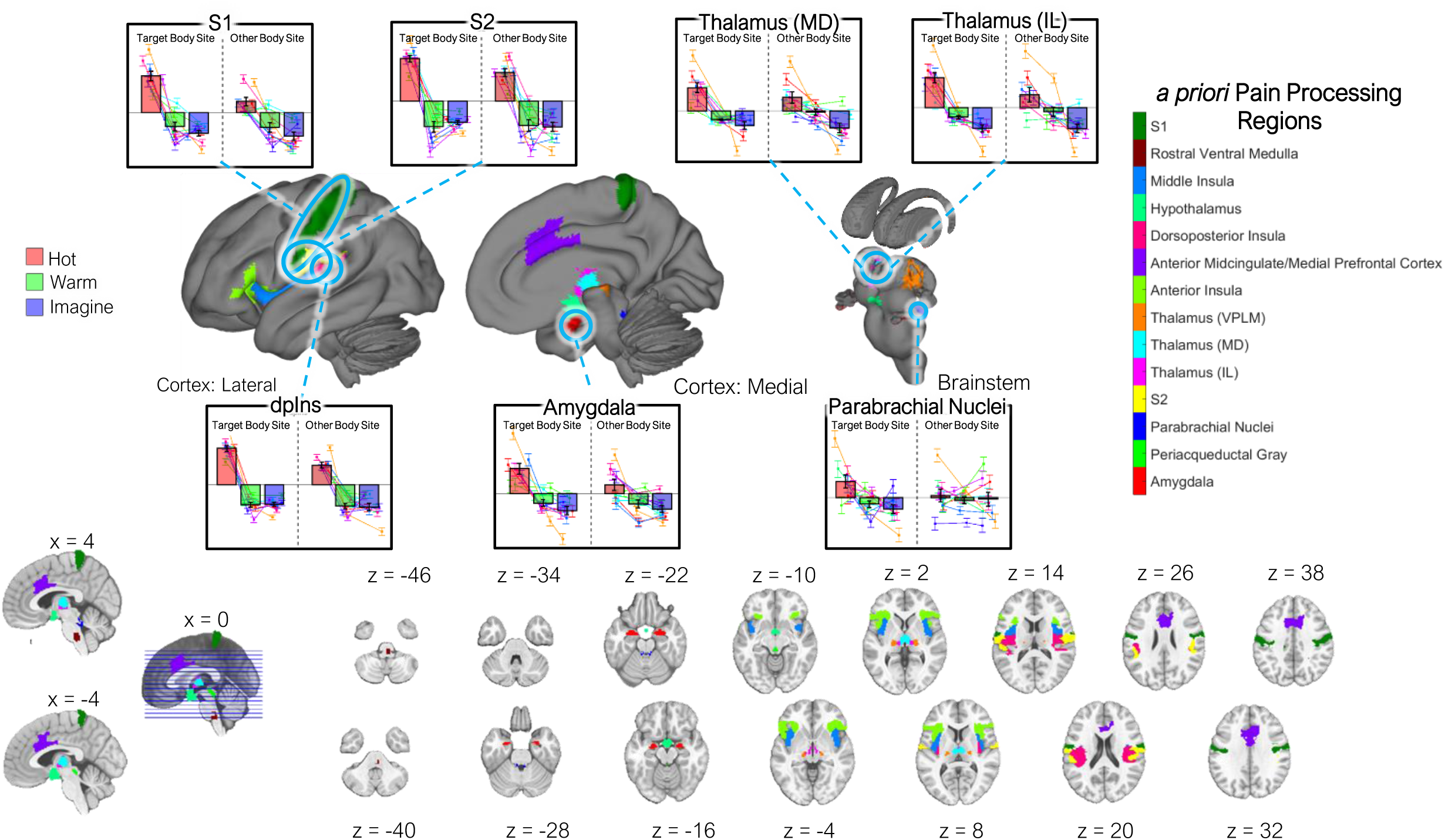
Nociception, body site-selectivity, and imagination effects in precision functional mapping of pain-processing regions of interest. Three regions – dpIns, S1, and S2– showed significant effects on individually defined nociceptive patterns (SVM weight maps trained on individual body sites) for [Hot - Rest], [Hot - Warm], and [Hot -Warm] x [Target vs. Other body sites]. Amygdala, thalamus, and parabrachial nuclei are also highlighted to show significant regions for four or more individual participants. The colors identify individual regions, which were defined bilaterally to allow for patterns to be differently lateralized for different body sites. Bar plots show individual and group pattern expression on independent test data for each of these regions. Colored lines show means and standard errors for individual participants. dpIns: dorsoposterior insula; S1: primary somatosensory cortex; S2: secondary somatosensory cortex; MD: Mediodorsal; IL: Intralaminar. Group data are presented as mean ± SEM across n = 9 participants.

Within the set of a priori pain-processing regions (Figure 5), 12 of 14 regions showed significant [Hot-Rest] and [Hot - Warm] pattern expression for individualized patterns in test data, and 3 of those regions – dpIns, S1, and S2 – also showed body site selectivity, fulfilling all three criteria for body site-specific nociceptive regions (Supplementary Figure 9). All three criteria were additionally significant at the single-participant level for a majority of participants in dpIns (8/9 participants), S1 (8/9 participants), and for at least 4/9 participants in S2, parabrachial nuclei, amygdala, and thalamus (IL and MD). Thus, multiple cortical and subcortical regions exhibited body site-selective nociceptive pattern representations, suggesting that fine-grained spatial encoding of painful stimuli is a robust and reliable feature of distributed pain-processing systems in the brain.

None of the 14 regions were significant for [Imagine-Rest] and [Imagine - Warm], nor did any region show body site selective responses for imagination (Figure 6). At the single participant level, 7/9 participants exhibited significant [Imagine-Rest] and [Imagine-Warm] pattern expression in at least one region, the most common of which was exhibited in S1 for 4/9 participants. Six of 9 subjects exhibited significant pattern expression in all three criteria including body site selectivity in at least one of 7 regions: parabrachial nuclei (3/9), hypothalamus (2/9), S1 (2/9), amygdala (1/9), periaqueductal gray (1/9), anterior insula (1/9), and middle insula (1/9). Thus, while some affective and subcortical regions were weakly engaged during imagined pain, the neural signatures lacked spatial specificity and robustness. No regions exhibited the consistent, body site-selective nociceptive representations seen during actual pain, suggesting that imagination does not evoke the same coding observed during experienced heat pain stimulation.

**Figure 6.**
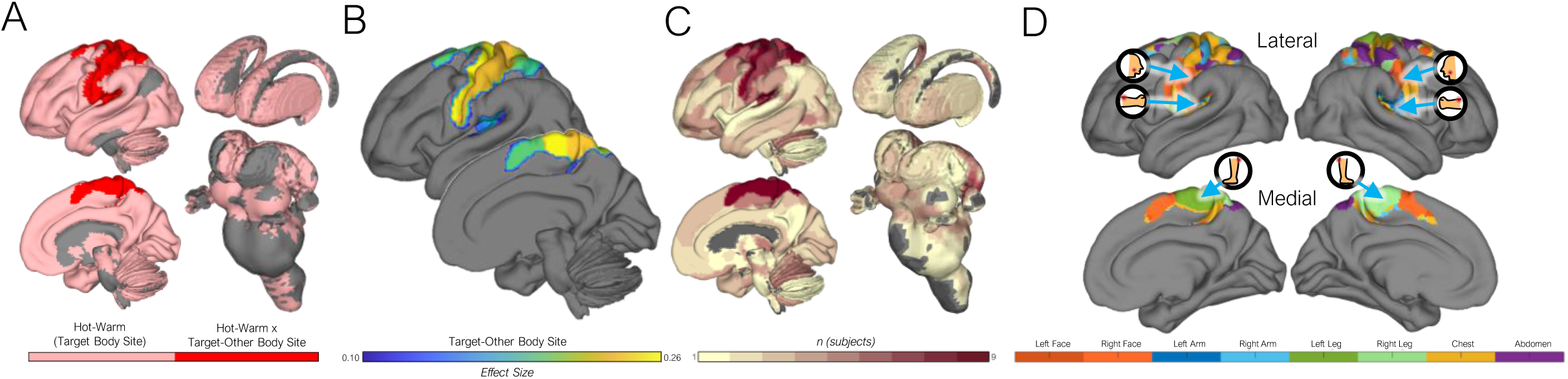
(A) Regions expressing significant Hot-Warm pattern expression shown in pink, and regions further expressing Target-Other Body Site specificity in expression shown in red. Regions shown include a priori pain pathways regions that are significant *p* < .05 and CANLab2024 regions after FDR correction. (B) shows an effect size map of the body site selective regions, and (C) shows the number of participants whereby [Hot-Warm] body site selectivity was significant after FDR correction at the single-participant level. (D) shows the body site selectivity at the group level for regions defined in (A) and (B) by computing t-value maps across body sites and taking a winner-take-all approach across body sites. The map shows clear contralateral lateralization in cortical representations for the left and right sides of the body, though midline body sites may preferentially activate ‘inter-effector’ regions. Note that this is a simplification and the body site-selective patterns analyzed do not require simple mapping from one region to one body site.

To assess whether imagined pain instead recruits pain-like representations in higher-order or associative regions beyond canonical nociceptive areas, we extended this analysis to a whole-brain search across 131 cortical and subcortical parcels (see Supplementary Figure 8; Supplementary Table S4; Methods). Ten parcels fulfilled all three criteria for nociceptive representations after FDR correction (red in Figure 6A, Supplementary Figure 9). Regions included parcels covering the pain-processing regions above (S1, S2, and posterior granular insula), A1, BA5, BA7 (anterior and postcentral), M1, SMA, and the dorsal intraparietal sulcus. 93 parcels showed significant [Hot – Rest] and [Hot – Warm] contrasts (pink in Figure 6A). The relative degree of body site selectivity is shown in Figure 6B in units of effect size (Cohen’s d). The dpIns (posterior granular insula), BA5, dorsal intraparietal cortex, M1, and SMA were significant in 9/9 individual participants, whereas a broader set of regions showed nociceptive representations in smaller numbers of participants (Figure 6C), pointing to potential individual differences in nociceptive representations. Note that the SVM patterns are agnostic to the spatial structure of body-site selective responses within each region, and reflect both large-scale classic somatotopy and more patchy or fine-grained patterns, including the possibility of multiple somatotopic maps contained within our ROIs.

Among significant regions with nociceptive representations, several also showed imagination-related activation. Lateral amygdala, anterior IPL, and anterior operculum showed significant [Imagine-Rest] and [Imagine-Warm] pattern expression at the group level. However, no regions showed nociceptive representations with body-site selectivity for imagined pain. At the single-participant level, no participants exhibited significant [Imagine-Rest] pattern expression in the three regions mentioned above. However, for [Imagine-Warm], 5/9 participants exhibited significant lateral amygdala pattern expression, and 4/9 participants exhibited significant anterior IPL and anterior opercular pattern expression. Only one participant showed any evidence for activation of body site-selective nociceptive representations during imagined pain (in BA8, BA9, and anterior putamen). Follow-up analyses of session- and run-level effects showed no evidence that imagined pain increasingly converged toward nociceptive representations with experience, indicating that the dissociation between imagined and experienced pain representations was stable across participants. These results show that imagination does not systematically activate nociceptive representations, and in the few cases it does, the activation was not selective to the body site imagined.

### Representational Similarity Analysis

Finally, to examine representational similarity between experienced and imagined pain in a modality- and classifier-independent manner, we conducted representational similarity analysis (RSA^74^) across the brain and within individual regions (Figure 7; see Methods and Supplementary Table S5). First, we constructed whole-brain voxelwise representational similarity matrices (RSMs) by computing within-participant Spearman’s spatial correlations from stimulus-evoked activity patterns in each of the 24 study conditions (i.e., Hot, Warm, and Imagine x 8 body sites per participant), shown in Fig. 7A and 7B. We calculated contrasts in pattern similarity related to target vs. other body sites (Fig. 7C left), similarity within each condition (Fig. 7C center), and similarity across (Hot, Warm) and (Hot, Imagine) conditions (Fig. 7C, right). The main comparison of interest for imagination was whether patterns were more similar for (Hot, Imagine) than for (Hot, Warm) and (Imagine, Warm), which would indicate ‘pain-like’ pattern activity. This analysis excluded images from the same run in calculating pattern correlations to avoid bias due to run contamination effects (see Methods).

**Figure 7:**
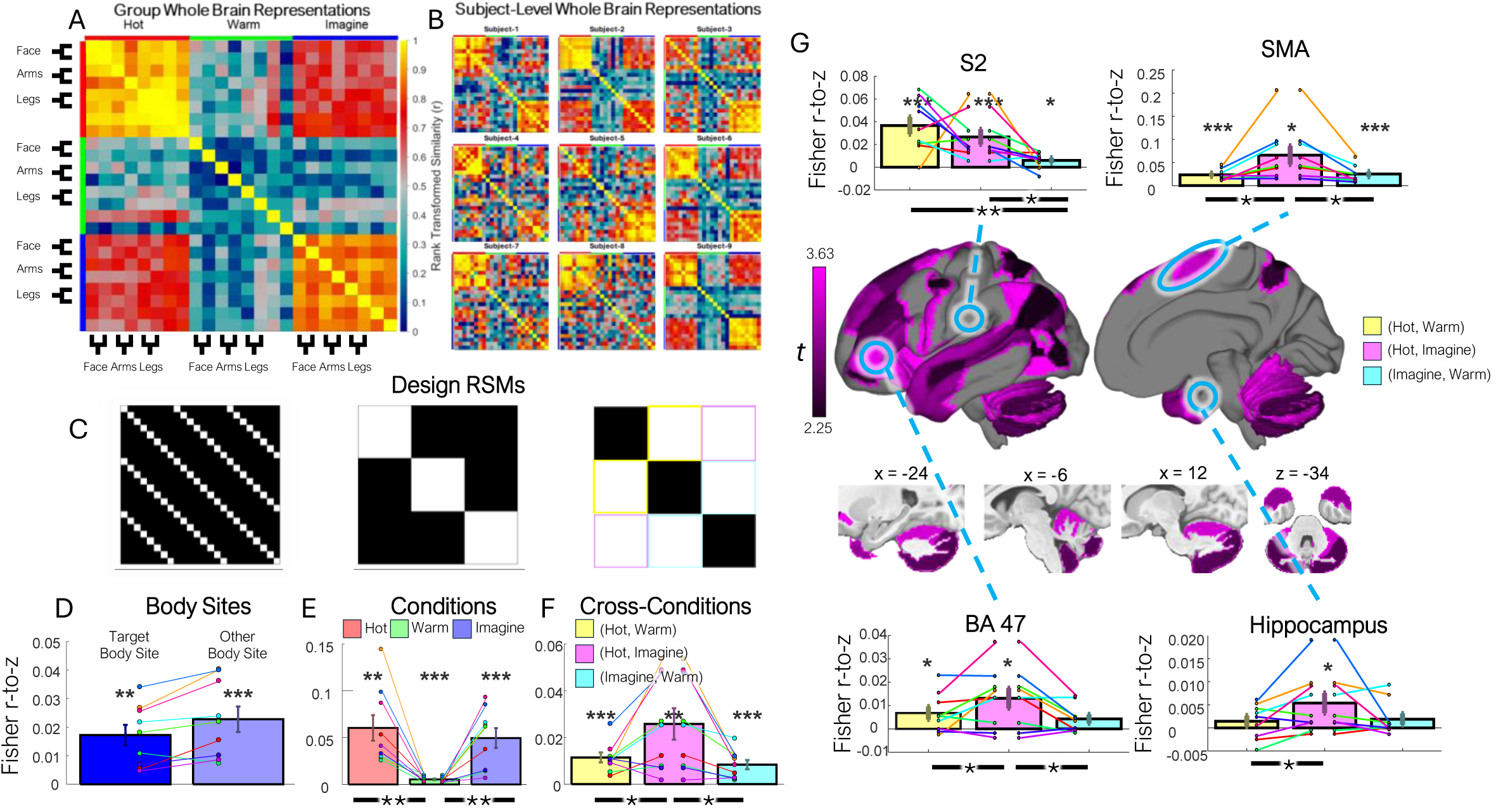
Representational Similarity Within and Between Conditions. (A) Group-level and (B) participant-level RSMs show considerable interparticipant similarity of whole-brain representations. (C) Model RSMs for Body Site Similarity (left), Intra-Conditional Similarity (center), and Cross-Conditional Similarity (right). White = 1 and black = 0 in indicator matrices. (D) the whole-brain target vs. other body site difference was significant for whole-brain representations, but target body site similarity exceeded other body site similarity in 10 somatosensory parcels (S1, S2/parietal operculum, insula; FDR q < .05; see Supplementary Figure 9) (E) The Hot and Imagine conditions exhibited significantly greater within-condition similarity compared to the Warm condition, with consistent effect direction across all participants. (F) In whole-brain RSA, the Hot and Imagine conditions (Hot, Imagine) showed significantly greater cross-condition similarity compared to Hot and Warm conditions (Hot, Warm) and Imagine and Warm conditions (Imagine, Warm). (G) Surface and subcortical maps of the 31 regions in which (Hot, Imagine) similarity simultaneously exceeds (Hot, Warm) and (Imagine, Warm) similarity (t-values shown are the average of the two) and at least one of these comparisons survive FDR correction, localizing the whole-brain phenomenon found in (F). The insets show Fisher z-transformed similarity for representative regions, with individual participants (lines) and group means ± s.e.m. (bars). Bars, group mean; lines, individual participants. * *p* < 0.05, ** *p* < 0.01 and *** *p* < .001. Statistical comparisons used one-sided (one-tailed) t-tests across participants (n = 9) on Fisher r-to-Z transformed correlations; parcelwise tests were FDR-corrected (q < .05); exact p-values are reported in the figure and Supplementary Tables 2–3.

**Figure 8:**
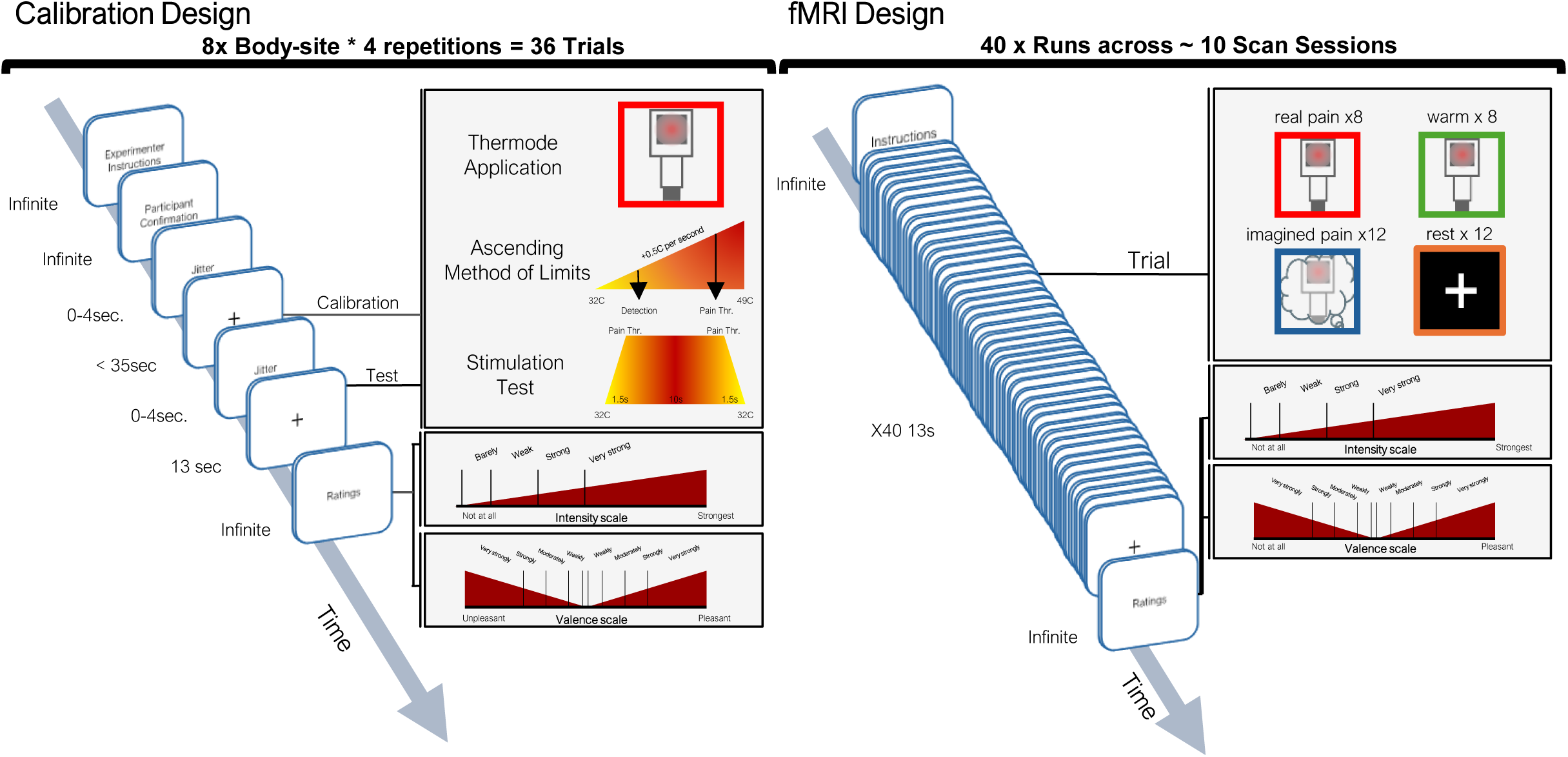
Study design for the deep phenotyping of real and imagined pain in eight body sites. The left depicts the experimental protocol for each trial of the pain calibration task. The right depicts the experimental protocol for each fMRI run targeting a single body site.

Whole-brain RSA did not evidence significant differences in similarity of activity patterns within a target body site compared to across other body sites (p=.993).. Patterns evoked during Hot (M=0.06(.01), t=4.32, p=.003, d=1.44) and Imagine (M=0.05(.01), t=4.86, p=.001, d=1.62) conditions were found to be significantly correlated across body sites, suggesting high consistency in the neural activation of these conditions. Warm patterns were also correlated across body sites (Warm: M=.004(.002), t=2.88, p=.02, d=0.96), but more weakly so (Hot-Warm: M=0.08(.04), t=3.96, p=.004, d=1.32; Imagine-Warm: M=0.04(.03), t=4.54, p=.002, d=1.51). The correlation between Hot and Imagine conditions (i.e., Hot/Imagine); M=0.03(.007), t=3.80, p=.005, d=1.27), was higher than the correlation between Imagine and Warm conditions (Imagine/Warm; M=0.008(.002), t=3.95, p=.004, d=1.32). Moreover, they exceeded the correlation between Hot and Warm conditions as well (Hot/Warm; M=0.01(.002), t=5.35, p=.001, d=1.78) despite the presence of actual somatosensory stimulation in both Hot and Warm conditions. Thus, at the whole-brain level, activity patterns reflect both body-site selectivity and similarity in evoked responses within test conditions. Crucially, imagination produces ‘pain-like’ activity patterns.

To test for neuroanatomical variability across brain systems in pattern similarity, we performed the same RSA analysis and contrasts for each of the 131 parcels spanning the brain (Supplemental Figure 8). Representations that were significantly greater for target body sites relative to other body sites were confined to 10 somatosensory parcels, 6 of which overlapped with SVM-derived body site selective nociception (S1, S2, A1, OP4, anterior BA7, and retroinsular/parabelt/belt complex) and 4 of which were unique (posterior operculum, anterior agranular insula, auditory transition area, and FOP1- BA43; Fig. S9 and Supplementary Table S5; FDR *q* < .05 one-tailed). The Imagine/Hot correlations were the dominant relationship in 74 out of 131 regions whereas Hot/Warm dominated in only 7 and Imagine/Warm in none (Figure S10 and Table S5). Of these 74 regions, 31 featured Imagine/Hot correlations significantly greater than both the Hot/Warm and Imagine/Warm correlations (each at uncorrected p < .05), with at least one comparison surviving FDR correction (q < .05; Figure 7G). These regions were broadly distributed across higher-order association cortex and most consistent in lateral prefrontal cortex (dorsolateral BA46/BA9/BA8 and ventrolateral BA45/BA47/IFS) and posterior parietal cortex (inferior parietal lobule, intraparietal sulcus, superior parietal lobule, and precuneus), together with higher-order visual and motion-sensitive areas (lateral occipital cortex, V7, and the MT+ complex — the single strongest region) and the cerebellum (Crus I/II and lobules I–X). Smaller contributions came from premotor cortex and SMA, auditory association cortex, orbitofrontal cortex (BA11), anterior temporal cortex (temporal pole), anterior insula, and the temporoparietal junction. Notably, the regions where Hot/Warm were greater than Imagine/Hot were concentrated in somatosensory and opercular cortex (S2, OP4, posterior operculum, and granular insula) and in subcortical nociceptive relays (thalamus, periaqueductal gray, and pons).

We calculated the spatial similarity between the 31 regions with similar imagined and experienced pain patterns with the principal gradient map of Margulies et al. 2016^66^, which defined a gradient from unimodal sensorimotor regions to transmodal regions that integrate information across modalities. The regions with imagined-experienced pain similarity overlapped with transmodal regions (mean correlation with the principal gradient r = 0.29 (s.e. = 0.004), t = 77.12, p < .001, d = 0.30). This widespread overlap suggests that imagining pain recruits many of the same cortical and subcortical systems involved in physically experiencing pain, and that cortical representations of imagined pain lie chiefly in transmodal association areas, rather than unimodal sensorimotor areas. Thus, imagining pain does not activate early nociceptive representations, but produces ‘pain-like’ activity patterns in high-level association areas and interconnected subcortical regions.

## Discussion

The ability to imagine pain underlies several crucial forms of cognition: it allows us to imagine our own future pain and pain under hypothetical decision scenarios, supporting decision-making and learning^13,14^, and allows us to imagine others’ pain, supporting empathy^47,49,73,75^. This study characterizes the neural representations of experienced and imagined pain across eight body sites using a deep-phenotyping, precision functional mapping framework, using three complementary analytic approaches to investigate the neural bases of imagined pain and the degree to which imagination activates early nociceptive representations and pain representations throughout the cortex and subcortex. These analyses converge to show that while experienced pain elicits widespread, robust, and body site-selective activations, imagined pain does not activate these same early body site-specific nociceptive representations. Instead, it activates selected regions, including recently-identified ‘inter-effector’ motor regions associated with an ‘action-mode’ network^69^, and pain-like patterns throughout a broad set of cortical and subcortical regions, particularly transmodal cortical areas at the top of the cortical processing hierarchy^66^. The engagement of the inferior intereffector region during imagined pain suggests that imagination recruits neural systems involved in preparing the body for action, even in the absence of nociceptive input, consistent with the idea that imagined pain accesses higher-order somato-cognitive representations rather than primary sensory nociceptive maps. These findings identify a new basis for pain imagination that both permits fictive activation of pain-related representations and preserves the distinction between imagination and experience. They suggest that mental imagery accesses higher-order pain-related representations without reinstating individualized nociceptive coding, placing principled limits on the representational scope of imagination.

### Idiosyncratic Pain Sensitivity and Neural Activation

Individualized calibration revealed notable variation in thermal sensitivity and pain tolerance across body sites, consistent with prior reports of regional differences in cutaneous innervation and nociceptive fiber density ^76^. Interestingly, the face—despite being highly innervated—exhibited the highest thermal pain tolerance (least sensitivity), whereas the arms and legs were most sensitive.

### Divergent and Overlapping Activation in Experienced and Imagined Pain

Consistent with prior studies and meta-analyses, pain (compared with non-painful warmth) activated a broad set of canonical nociceptive regions, including the thalamus, insula, S2, and cingulate cortex, alongside higher-order areas implicated in transmodal cognitive function ^77,78^. Many of these were activated in every or nearly every participant in individualized analyses across the multiple scanning days. A notable but expected absence of S1 activation arises because S1 responses to pain are spatially heterogeneous, somatotopically organized, and highly individualized. Regional mean activation can underestimate or obscure robust nociceptive representations that are evident at the pattern level (see Supplementary Figure 7). In contrast, imagined pain elicited activity in a more restricted set of regions, with more diversity in activated areas across individuals. Group-level activation consistent across individuals was found in only a few regions, and regions overlapping with thermal pain included the reticular formation, insula, premotor cortex, dACC, and vlPFC. Unique to imagined pain was activation in BA6, lateral temporal cortex, hippocampus and parahippocampus, basolateral amygdala, pons, and medulla, primarily at the group level and with limited consistency across individual participants. Imagined pain additionally engaged portions of S1 and premotor cortex at the group level, though these effects were spatially heterogeneous and less consistent across individuals than activity in action-related motor and premotor regions. Furthermore, the majority of participants activated a focal region consistent with the ventral inter-effector zone of the primary somatomotor cortex^69^, a motor–premotor area with body-wide representation. Many individual participants also showed activation in adjacent premotor regions, including the middle inter-effector motor region, as well as posterior somatosensory opercular cortex. The inter-effector regions, premotor cortex, anterior putamen, medial thalamus, and somatosensory operculum together comprise key components of the recently characterized “action-mode network,” a cortical–subcortical system implicated in motivated action, autonomic regulation^82,83^, and pain control^84^. The engagement of this network during imagined pain, in the absence of nociceptive input, suggests that pain imagery may preferentially recruit neural systems involved in action-oriented simulation and bodily state representation rather than canonical afferent nociceptive pathways. Precision mapping reveals body site-selective representations of experienced but not imagined pain in nociceptive regions

While previous studies have investigated pain imagery^15,46,57,59,60,61^ and related phenomena including placebo^85^, mindfulness^86^, and hypnosis^87^, most previous studies using pattern recognition to perceptual representations underlying mental imagery have focused on other domains, particularly vision^41,88^, audition^89^, and olfaction^90^. Thus, these findings represent the first attempt, to our knowledge, to assess whether mental imagery can activate individualized nociceptive representations. This is an important direction for future studies, as activation of ‘pain-related’ gross anatomical regions has been shown to reflect non-specific processes related to emotion, attention, motor function, and other processes^49,91–93^, whereas multivariate patterns are considerably more specific^55,94^.

We first assessed reliability of the imagination task by quantifying within-subject stability of imagined-pain multivariate patterns across runs and days, using cross-session correlations and ICC of pattern expression values. These analyses demonstrate robust temporal reproducibility, supporting consistent task engagement. We then identified individualized nociceptive representations that fulfilled three criteria: (1) activation during painful heat, (2) greater activation for heat than non-painful warmth, and (3) body site-selectivity, all established in independent test data from different scan days. These representations were located in early nociceptive regions that receive mono- or di-synaptic input from spinal nociceptive afferents^52,95,96^, including dpIns, S2, S1, ventrolateral and medial thalamus, and aMCC, but also in some transmodal association cortices and subcortical regions (e.g., thalamus, amygdala, basal ganglia) not directly associated with nociception. These findings add to a growing field identifying body site-selectivity in motor activity, touch, and pain beyond the somatomotor and insular cortices^97–101^, and suggest that pain somatotopy is supported by a broad, hierarchical network spanning sensory, affective, and integrative systems.

While imagined pain activated some of the same broad areas as experienced pain, particularly in the ‘action-mode network’ (see above), imagination did not activate nociceptive representations, and did not show the pattern of body site-selectivity apparent during experienced pain in any one region. In most cases this was also true at the individual-participant level. Some exceptions were that (1) six subjects activated body site-selective nociceptive patterns in at least one region among parabrachial nuclei, hypothalamus, S1, amygdala, periaqueductal gray, anterior insula, and middle insula, and (2) seven participants showed pain-pattern expression in functionally diverse but largely non-overlapping regions, with the highest overlap featured in S1, overlapping among four participants. Importantly, practice-related analyses indicated that imagined pain representations did not systematically increase in similarity to nociceptive patterns across sessions or runs, suggesting stable engagement rather than progressive learning or exaggeration with experience. Overall, the findings suggest that imagined pain does not rely on topographically organized representations in nociceptive regions, but may rely on distributed, higher-order representations integrating affective memory, visual imagery, and conceptual knowledge in an individualized fashion. Importantly, the selective and inconsistent recruitment of nociceptive regions during imagination—and during other non-somatosensory experiences such as viewing emotionally aversive images—raises the possibility that such engagement of pain-related representations in affective or conceptual contexts may be one mechanism linking mental simulation to the future development of pain disorders in otherwise healthy individuals. This interpretation aligns with a growing body of work from multiple groups demonstrating the involvement of pain-related brain patterns in non-nociceptive tasks and their predictive relevance for clinical outcomes.^102,103,13,86,104–107^

### Representational similarity reveals pain-like patterns across transmodal cortical areas

Representational Similarity Analysis (RSA) complements the other analyses presented here by providing a different lens on how pain-related and imagined states are encoded in the brain. Rather than isolating nociceptive representations per se, RSA assesses the similarity of multivoxel activity patterns evoked by painful, nonpainful, and imagined stimuli^74,108^. These patterns may partially reflect nociceptive input but are also shaped by a broader set of processes, including attention, affect, cognitive appraisal, and motivational state^109–111^. Importantly, RSA, like the individualized machine learning–based pattern recognition methods discussed above, enables the identification of idiographic neural representations that vary across individuals and conditions, without requiring shared anatomical topography across participants^112,113^. This makes RSA particularly well-suited for precision neuroimaging approaches that seek to model functional brain representations at the level of individual experience.

The RSA analyses showed that patterns evoked during experienced pain were more similar to those evoked by pain imagination than nonpainful warmth across widespread areas of the cortex and cerebellum. Critically, the pain-specific component of this similarity — greater resemblance to experienced pain than to warmth — was absent in primary somatosensory, opercular, and early auditory cortex (e.g., S2, OP4, A1), where nonpainful warmth and heat already evoke similar, body-site-specific patterns, and was instead concentrated in higher-order association cortex, including lateral prefrontal, posterior parietal, and temporoparietal prefrontal regions. Thus, imagined pain evokes a globally distributed pattern of ‘pain-like’ activity in transmodal areas^66^ thought to underlie higher-order abstractions and conceptual maps^114–116^. Several areas, including posterior temporal-parietal junction, and anterior temporal cortices, have received substantial attention as substrates of encoding abstract conceptual relations (i.e., cognitive maps) and representations that integrate narrative meaning across long time scales^117–119^, as well as in predictive processing-related regulation of posterior sensory areas e.g., ^120,121^. Thus, these results fit with the hypothesis that imagined pain does not re-activate early sensory representations, but produces pain-like activity patterns in higher-order maps of conceptual space, potentially reflecting embodied simulation^122,123^ or predictive processes that facilitate disambiguating sensory input in favor of expected percepts ^124,125^. This latter concept provides a link between imagination and placebo effects, which also activate the vmPFC and other transmodal areas like posterior temporoparietal junction^126–129^. Thus, imagination, like expectation, may shift internal models towards a pain-like state^130,131^, facilitating early detection and priming decision-making systems to respond quickly to actual nociceptive input.

### Limitations and future directions

This study has several limitations that could be addressed in future studies. First, nociceptive representations could be defined based on criteria of specificity and generalizability in a more comprehensive fashion. This has been possible with more macroscopic, population-level patterns by testing across studies^55,56^, and there is a natural tension between testing multiple body sites and including many non-somatic conditions within the same individuals (i.e., imagining pain). Secondly, the lines between nociceptive sensory representations, pain perception, and other sequelae of pain such as action planning are unclear. There is perhaps no single study that can definitively disentangle them, and converging evidence across neuroscience methods (e.g., invasive electrophysiological recording^132^ will be required to continue to understand the neuroscientific basis of pain imagery. In addition to these issues, future studies could address the ability to activate perceptual representations with training, in special populations with chronic pain – particularly pain engaged by non-nociceptive stimuli. For example, reports of nociceptive pain-signature expression in somatoform pain disorders during non-noxious image viewing highlight the need for approaches capable of distinguishing individualized nociceptive coding from higher-order pain-related representations (e.g., Boëgarts et al., 2023^133^, in functional neurological disorder). In addition, the propensity to activate pain-like representations could be investigated as a neuromarker for vulnerability to chronic pain after injury or insult. This propensity may also confer the ability to empathize with others in pain, increasing either empathic care or empathic distress^62,134^. For example, some areas with the greatest pain - imagined pain similarity in this study have also been targeted with neurostimulation^135,136^ to promote pain empathy, and lesions (e.g., of vmPFC) reduce empathy^137,138^. Thirdly, this study was not designed to test demographic moderators of pain representations. The intensive deep-phenotyping approach prioritized within-subject precision, necessarily limiting sample size. Nonetheless, the sample included heterogeneity in age, sex, ethnicity, handedness, and prior pain research exposure. Systematically examining how individualized pain representations vary across demographic and cultural factors will require substantially larger samples that combine deep phenotyping with population-level diversity, representing an important direction for future work. Lastly, we did not collect trial-by-trial ratings of imagery vividness or intensity. However, multiple lines of evidence support consistent engagement with the imagined pain task despite the absence of such data. Imagined pain elicited stable, body-site–specific multivariate neural patterns that were reproducible across runs and across days within individuals. Moreover, practice-related analyses revealed no systematic convergence of imagined pain toward nociceptive representations with experience, arguing against progressive exaggeration or demand effects. Together, these findings indicate that imagined pain engaged reliable, individual-specific neural representations over extended task performance, even in the absence of explicit phenomenological ratings. Relatedly, alternative imagination instructions featuring a more cognitive focus (e.g., imagining one’s pain to be so intense one cannot focus on anything else) may result in findings that differ from the results shown here, where the focus was on intense sensory/emotional imagery. The present findings provide representational constraints on pain imagination but do not, on their own, establish mechanistic accounts of planning, empathy, or future-oriented behavior. Rather, they delineate which pain-related representations are accessible to imagination, thereby enabling future studies that combine precision functional mapping with explicit behavioral and causal manipulations. Ultimately, addressing how imagined pain interacts with nociceptive processing, or how such representations are altered in chronic pain or psychiatric conditions, will require experimental designs beyond the scope of the present precision-mapping framework.

By leveraging individualized calibration and precision functional mapping of densely sampled data over repeated fMRI sessions per participant, this study demonstrates that pain imagination engages pain-like patterns across a broad set of cortical and subcortical regions, including activation of multiple regions associated with the recently defined ‘action mode network’ and pain-like multivariate representations in transmodal cortical regions. It does not activate targeted, body site-specific nociceptive representations in early somatosensory regions. This provides a way for imagination to engage decision-making, planning, and potentially perceptual inference processes to simulate pain, while maintaining a distinction between real and imagined experience by virtue of selective activation of sensory representations by real experience. By delineating which pain-related representations are and are not accessible to imagination, the present findings constrain models of how non-nociceptive processes might influence pain without directly triggering sensory alarm systems. Furthermore, by dissociating higher-order, transmodal pain-like representations from early nociceptive coding, the present framework provides a foundation for future studies to test how imagination influences pain-related behavior—such as prediction, regulation, empathy, and decision making—even in the absence of sensory reinstatement. Prior work showing certain clinical populations engaging nociceptive pain signatures in the absence of direct noxious stimulation cannot yet determine whether early, individualized nociceptive representations are being reinstated or whether such responses reflect higher-order pain-related processes. The present framework provides a more precise approach, enabling the identification of individualized nociceptive representations and testing whether—and under what conditions—they can be reexpressed by non-nociceptive inputs.

More broadly, these findings can establish a foundation for future work examining how imagined and experienced pain are represented across the brain, how pain-related imagery may interact with cognition, learning, and social processes, and how these representations may differ across individuals/contexts, or be altered in clinical or developmental contexts.

## Methods

This research complied with all relevant ethical regulations; the study protocol was approved by the Dartmouth College Institutional Review Board (Committee for the Protection of Human Subjects). This study is part of a registered national clinical trial examining placebo effects on pain (ClinicalTrials.gov: NCT04653064). A preregistration on OSF (https://osf.io/zv4ec/) was created prior to analysis for the broader Warmth, Anticipation, Somatotopy, Aversiveness, and Body Imagery (WASABI) N-of-Few protocol, which encompasses body mapping, pain imagination, and regulation components. Although data collection had begun at the time of preregistration, no analyses of the present dataset had been conducted, and all analyses reported here postdate the preregistration.

### Participants

Nine participants (4 females, 5 males; 7 right-handed, 2 left-handed; 6 White, 2 Asian, and 1 identifying as multi-racial) were recruited from the Dartmouth Department of Psychological and Brain Sciences between the ages of 24-56 (M=35.63 years, SD=11.62 years). Written informed consent was obtained in accordance with the Declaration of Helsinki. Preliminary eligibility of participants was determined through an MRI safety screening form. No significant abnormalities were detected in any participant’s structural scans. Participants were compensated $25/hour of scanning, $10/hour of online/behavioral testing, and $7.25/hour of behavioral setup, and were offered a completion bonus matching their earnings up to the conclusion of data collection; participants who discontinued were paid a prorated rate for their time of participation. Sex was determined by self-report. The study was not designed or powered to analyze sex differences; sex-disaggregated data are provided in Table 1.

**Table 1:** Participant Demographics.

| Participant No | Sex | Age<sup>1</sup> | Detection Threshold |  | Pain Tolerance |  | Pain Intensity |  | Pain Valence |  |
| --- | --- | --- | --- | --- | --- | --- | --- | --- | --- | --- |
|  |  |  | M | SE | M | SE | M | SE | M | SE |
| 1 | M | 50s | 39.56 | 1.46 | 47.23 | 0.34 | 51.41 | 0.49 | -54.72 | 1.03 |
| 2 | F | 20s | 37.50 | 1.07 | 47.86 | 0.23 | 48.84 | 1.20 | -49.38 | 2.24 |
| 3 | F | 30s | 34.52 | 0.69 | 45.73 | 0.43 | 36.44 | 2.15 | -39.38 | 2.16 |
| 4 | M | 30s | 38.63 | 1.74 | 47.14 | 0.37 | 27.56 | 5.29 | -28.84 | 5.54 |
| 5 | M | 30s | 39.30 | 1.39 | 47.53 | 0.41 | 26.97 | 3.33 | -29.63 | 3.72 |
| 6 | M | 40s | 39.64 | 1.11 | 47.44 | 0.23 | 32.69 | 3.22 | -34.06 | 3.75 |
| 7 | F | 20s | 38.34 | 1.33 | 46.50 | 0.48 | 26.94 | 2.93 | -24.78 | 3.44 |
| 8 | M | 20s | 39.64 | 1.13 | 46.13 | 0.67 | 24.84 | 3.50 | -27.50 | 3.83 |
| 9 | F | 30s | 36.14 | 0.96 | 45.60 | 0.50 | 49.92 | 2.36 | -46.50 | 9.01 |

<sup>1</sup>Age is obfuscated to prevent identification of participants. Race and handedness are reported in aggregate in the Methods for the same reason.

### Experimental Protocol

#### Thermal Stimulation

For thermal stimulation, a TSA2 Advanced Thermosensory Stimulator (TSA2^139^; Medoc Advanced Medical Systems, Ramat Yishai, Israel) with a 16 mm^2^ Peltier thermode end plate was used. This end plate delivered stimulation to, from top-to-bottom, the bilateral mandibular base (face), xiphoid process (chest), bilateral volar flexor below the cubital fossa (arms), umbilical surface of the abdomen below umbilicus (abdomen), and bilateral medial heads of the gastrocnemius below popliteal fossa (legs), and was affixed to each of the eight body sites with compression sleeves. Thermode stimulation induces a fast, sustained, and well-calibrated sensation at levels safely calibrated to each individual participant. Stimulus application was computer controlled and recorded with PsychoPy3 v2020.2.5 ^140^.

#### Heat Calibration with the Ascending Method of Limits

Participants varied in prior exposure to pain research at enrollment; however, all participants underwent extensive thermal calibration prior to neuroimaging, ensuring familiarity with the thermode and painful sensations, and to determine the temperature sensitivity and pain tolerance of each bodysite to be used in the fMRI setting. Calibration was done using an Ascending Method of Limits procedure delivered using PsychoPy3. Each trial of the procedure targeted a randomly-selected body site, with the experimenter affixing the thermode at the beginning of each trial. After one trial each of eight body sites, the process was repeated three additional times for a total of 32 total trials per participant. Each trial begins with the participant indicating that they are ready to receive a thermal stimulus with the space-bar. Stimulation began after a 0-4s jitter period at 32 degrees Celsius, escalating a half-degree per second and continuing to escalate until the thermode reached a maximum of 49 degrees Celsius, after which the trial would terminate. Participants indicated with the spacebar at the moment they could detect a change in temperature, which indicated their detection threshold. A subsequent spacebar press terminated the stimulation and indicated their pain tolerance. Following a subsequent randomized jitter period of 0-4s, a 13s stimulation pulse, peaking at the terminated temperature for 10s with a 1.5s ramp-up and ramp-down time, was delivered to a body site. The participants were then instructed to make unipolar responses to questions regarding the perceived stimulus intensity (‘How intense was that overall?’) and bipolar responses to valence (‘How pleasant was that overall?’) of that pulse using a mouse-pointer. Responses on each pole were made on a generalized Labeled Magnitude Scale (gLMS; ^141,142^ with anchors for No Sensation (0% of scale length), Barely Detectable (1.4% of scale length), Weak (6.1% of scale length), Moderate (17.2% of scale length), Strong (35.4% of scale length), Very Strong (53.3% of scale length), and Strongest Imaginable (100% of scale length). Intensity ratings were anchored between “Not [intense] at all” (left) to “Strongest Imaginable Sensation” (right), while valence ratings ranged from “Strongest Imaginable Unpleasantness of any kind” (left) to “Strongest Imaginable Pleasantness of any kind” (right), with neutral valence at the center of the scale.

#### Neuroimaging Sessions

During each neuroimaging session, participants were tasked to undergo runs (i.e., scan repetitions) consisting of 8.89 minutes of pseudo-randomized presentations of eight high-heat trials (bodysite-specific pain tolerance temperature), eight low-heat ‘warm’ trials (bodysite-specific detection threshold + 1 degree celsius), or 24 no-stimulation trials, 12 of which were prepended with a cue to imagine high-heat stimulation. The Peltier thermode was manually positioned to stimulate each body site, randomized and then signaled to the experimenter by an instruction message prior to each run. All thermal stimulation was applied by the experimenter and secured in place; participants did not manipulate or hold the stimulator during stimulation. Participants were presented with an imagination instruction (featured in ^109^ that they were asked to memorize prior to the first run. Each trial consisted of either a white fixation cross during which the individual would receive a 13 second thermal stimulation, or a 2-second picture cue of the designated body site with a text reminder instruction to imagine followed by 11 seconds of an orange fixation cross indicating the need to sustain the imagination. Upon completion of a run, participants were then instructed to use a slider to rate (1) the valence and (2) the stimulus intensity of the most painful thermal event, and (3) their overall bodily comfort after the run using a gLMS scale.

The imagination instruction prior to the start of each fMRI session emphasized vivid sensory imagery as follows: “During this scan, you will occasionally see picture cues of a body part where the thermode is attached. When you see this, try to imagine as hard as you can that the thermal stimulations are more painful than they are. Try to focus on how unpleasant the pain is, for instance, how strongly you would like to remove yourself from it. Pay attention to the burning, stinging and shooting sensations. You can use your mind to turn up the dial of the pain, much like turning up the volume dial on a stereo. As you feel the pain rise in intensity, imagine it rising faster and faster and going higher and higher. Picture your skin being held up against a glowing hot metal or fire. Think of how disturbing it is to be burned, and visualize your skin sizzling, melting and bubbling as a result of the intense heat.”

During fMRI data acquisition, imagination trials were prepended by the following reminder text: “Picture your [bodysite] being held up against a glowing hot metal or fire. Visualize the skin on your [bodysite] sizzling, melting and bubbling.”

### FMRI Data Acquisition

#### MRI System and Hardware

MRI data acquisition is reported here using the OHBM Committee on Best Practice in Data Analysis and Sharing (COBIDAS) guidelines <u>(Nichols et al. 2017)</u> and a full checklist available in the Supplementary Materials. All MRI data were acquired on a 3.0 Tesla Siemens MAGNETOM Prisma scanner (Siemens Medical Systems, Erlangen, Germany) at the Dartmouth Brain Imaging Center, using a 32-channel head coil. Participants’ heads were stabilized with inflatable cushions and foam padding.

#### Structural MRI (T1-weighted) Acquisition

High-resolution T1-weighted structural images were acquired using a 3D MPRAGE sequence with:

- TR = 2000 ms, TE = 2.11 ms, TI = 1000 ms, flip angle = 8°
- Voxel size = 0.8 × 0.8 × 0.8 mm, slice thickness = 0.8 mm
- Base resolution = 320, matrix = 320 × 320 (FoV = 256 × 256 mm)
- Orientation = sagittal
- Slices = 224, slice oversampling = 14.3%
- Fat suppression = water excitation (fast)
- Parallel imaging = GRAPPA (acceleration factor 3, 32 reference lines)
- Bandwidth = 240 Hz/pixel, echo spacing = 7.2 ms
- Magnetization preparation = non-selective inversion recovery AutoAlign (Head > Basis) was enabled.

#### Functional MRI (BOLD) Acquisition

Functional images were collected using a T2-weighted multiband EPI* sequence with the following parameters:

- TR = 460 ms, TE = 27.2 ms, flip angle = 44°
- Voxel size = 2.7 × 2.7 × 2.7 mm, slice thickness = 2.7 mm, no slice gap
- Base resolution = 82, matrix = 82 × 82 (FoV = 220 × 220 mm)
- Slices = 56, orientation = axial/transversal, multislice mode = interleaved
- Phase-encoding direction = A→P
- Multiband acceleration factor = 8
- Bandwidth = 3048 Hz/pixel, EPI factor = 82, echo spacing = 0.49 ms
- Fat suppression = fat-sat, RF excitation = standard
- iPAT / GRAPPA = none
- Prescan normalization = on, distortion correction = off

Each functional run consisted of 1160 volumes, reconstructed as magnitude images, with no dummy scans removed (ignore = 0). Single-band reference images were acquired.

### Analysis

#### Self-Report

All self-report data were analyzed using Matlab (MATLAB, The MathWorks, Inc., Natick, Massachusetts, United States). For the Ascending Method of Limits Calibration, participants rated the intensity and valence of each heat-stimulation trial. For each fMRI run, participants rated the intensity and valence of the highest-pain sensation they felt after every fMRI run, and their overall body comfort to assess the safety of continued scanning.

#### Preprocessing

To mitigate the effects of magnetic disequilibrium at the start of each run, six TRs (2.76 seconds) were spent to allow the scanner to reach a steady state, and were discarded during subsequent analysis. Results included in this manuscript come from preprocessing performed using fMRIPrep 23.1.4 ^143,144^; RRID:SCR_016216), which is based on Nipype 1.8.6 ^145,146^; RRID:SCR_002502).

Preprocessing of B0 inhomogeneity mappings. For each participant, a B0- nonuniformity map (or fieldmap) was estimated based on two (or more) echo-planar imaging (EPI) references with FSL topup ^147^.

##### Anatomical data preprocessing

All T1-weighted (T1w) images were corrected for intensity non-uniformity (INU) with N4BiasFieldCorrection ^148^, distributed with ANTs ^149^, RRID:SCR_004757). The T1w-reference was then skull-stripped with a Nipype implementation of the antsBrainExtraction.sh workflow (from ANTs), using OASIS30ANTs as target template. Brain tissue segmentation of cerebrospinal fluid (CSF), white-matter (WM) and gray-matter (GM) was performed on the brain-extracted T1w using FSL fast, RRID:SCR_002823^150^. An anatomical T1w-reference map was computed after registration of T1w images (after INU-correction) using mri_robust_template (FreeSurfer 7.3.2^151^). Brain surfaces were reconstructed using recon-all (FreeSurfer 7.3.2, RRID:SCR_001847^152^, and the brain mask estimated previously was refined with a custom variation of the method to reconcile ANTs-derived and FreeSurfer-derived segmentations of the cortical gray-matter of Mindboggle (RRID:SCR_002438^153^).

##### Functional data preprocessing

For each of the BOLD runs found per participant (across all tasks and sessions), the following preprocessing was performed. First, a reference volume and its skull-stripped version were generated by aligning and averaging single-band references (SBRefs). Head-motion parameters with respect to the BOLD reference (transformation matrices, and six corresponding rotation and translation parameters) are estimated before any spatiotemporal filtering using mcflirt (FSL, ^154^. The estimated fieldmap was then aligned with rigid-registration to the target EPI (echo-planar imaging) reference run. The field coefficients were mapped on to the reference EPI using the transform. The BOLD reference was then co-registered to the T1w reference using bbregister (FreeSurfer) which implements boundary-based registration ^155^. Co-registration was configured with nine degrees of freedom to account for distortions remaining in the BOLD reference. First, a reference volume and its skull-stripped version were generated using a custom methodology of fMRIPrep. Several confounding time-series were calculated based on the preprocessed BOLD: framewise displacement (FD), DVARS and three region-wise global signals. FD was computed using two formulations following Power (absolute sum of relative motions, ^156^ and Jenkinson (relative root mean square displacement between affines^154^) FD and DVARS are calculated for each functional run, both using their implementations in Nipype (following the definitions by ^156^). The three global signals are extracted within the CSF, the WM, and the whole-brain masks. Additionally, a set of physiological regressors were extracted to allow for component-based noise correction (CompCor^157^). Principal components are estimated after high-pass filtering the preprocessed BOLD time-series (using a discrete cosine filter with 128s cut-off) for the two CompCor variants: temporal (tCompCor) and anatomical (aCompCor). tCompCor components are then calculated from the top 2% variable voxels within the brain mask. For aCompCor, three probabilistic masks (CSF, WM and combined CSF+WM) are generated in anatomical space. The implementation differs from that of Behzadi et al. in that instead of eroding the masks by 2 pixels on BOLD space, a mask of pixels that likely contain a volume fraction of GM is subtracted from the aCompCor masks. This mask is obtained by dilating a GM mask extracted from the FreeSurfer’s aseg segmentation, and it ensures components are not extracted from voxels containing a minimal fraction of GM. Finally, these masks are resampled into BOLD space and binarized by thresholding at 0.99 (as in the original implementation). Components are also calculated separately within the WM and CSF masks. For each CompCor decomposition, the k components with the largest singular values are retained, such that the retained components’ time series are sufficient to explain 50 percent of variance across the nuisance mask (CSF, WM, combined, or temporal). The remaining components are dropped from consideration. The head-motion estimates calculated in the correction step were also placed within the corresponding confounds file.

The confound time series derived from head motion estimates and global signals were expanded with the inclusion of temporal derivatives and quadratic terms for each ^158^. Frames that exceeded a threshold of 0.5 mm FD or 1.5 standardized DVARS were annotated as motion outliers. Additional nuisance timeseries are calculated by means of principal components analysis of the signal found within a thin band (crown) of voxels around the edge of the brain, as proposed by ^159^. The BOLD time-series were resampled into standard space, generating a preprocessed BOLD run in MNI152NLin2009cAsym space. First, a reference volume and its skull-stripped version were generated using a custom methodology of fMRIPrep. The BOLD time-series were resampled onto the following surfaces (FreeSurfer reconstruction nomenclature): fsnative, fsaverage6. The BOLD time-series were resampled onto the left/right-symmetric template “fsLR” ^160^.

### General Linear Modelling (GLM)

#### Participant-Session Level GLM

First-level GLM analyses were conducted in SPM12 at the level of each participant’s session, using SPM FAST to account for temporal autocorrelation ^161,162^. The experimental runs for each session were concatenated and boxcar regressors, convolved with the canonical hemodynamic response function, were constructed to model periods for the 13s thermal stimulations (hot and warm), 2s imagination cue presentation, 11s imagination period, and post-trials rating periods. Resting fixation cross epochs were used as an implicit baseline. Each session was high-pass filtered at 180s. Nuisance covariates included 24 motion regressors, 5 temporal components (tCompCor), 5 anatomical components (aCompCor), and cerebrospinal fluid from fmriprep output. Spike regressors were also generated using the CANlab *Tool outliers()* that implements a number of spike detection methods and flags outlying images using Mahalanoubis distance. This procedure resulted in a beta image for each participant’s run.

#### Participant-Level GLM

Participant-level GLM analyses were conducted by first smoothing each participant’s run beta images with a 6mm Full-Width Half-Max smoothing kernel and subjecting these images to robust regression using the *regress(‘robust’)* command from CANlab Tools. The design matrix used Warm images as the intercept, and included dummy- coded condition regressors of interest (Hot vs. Warm and Imagine vs. Warm). Eight regressors were used to average body site effects (using the *CANlab Tool create_orthogonal_contrast_set())*. Cerebral spinal fluid and white matter regressors were included as regressors of interest. Furthermore, session and run were centered and included as well.

The resulting contrasts of interest (Hot vs. Warm and Imagine vs. Warm) for each participant were thresholded with the false-discovery rate (FDR^68^, *q* < .05 to make inferences about whether a voxel was activated or deactivated for a participant’s contrast.

#### Group-Level Univariate Analyses

Group-level GLM analyses were conducted by taking the resulting participant-level contrast images of interest and performing robust parcel wise regression using 518 regions of interest (ROIs i.e., parcels) defined by the CANLab 2024 atlas and the *robust_parcelwise()* command from CANlab Tools. The results were thresholded with the false-discovery rate, q < .05 across ROIs to make inferences about whether an ROI was activated or deactivated on average.

#### Multivoxel Pattern Reliability Analyses

To assess the consistency of imagined pain representations over extended task performance, we quantified within-subject reliability of multivoxel activation patterns across runs and across scanning sessions. Reliability was evaluated using cross- session Pearson correlations of pattern expression values within each subject. Statistical significance was assessed using permutation-based null distributions generated by randomly shuffling session labels within-subject.

To provide a conservative estimate of reliability, we additionally conducted split-half analyses comparing pattern expression derived from odd versus even runs across sessions. All reliability analyses were performed separately for each body site, allowing us to assess whether imagined pain representations were both temporally stable and body site specific within individuals.

#### Precision Functional Mapping with Support Vector Machine

Precision functional mapping was performed by training a support vector machine (SVM) experienced pain classifier, classifying hot from warm images for each body site in each participant. Using the *xval_SVM()* function from CANlab Tools, we trained SVM models on each participant’s unsmoothed, run-level replicate images for each body site. Patterns were extracted from a priori pain ROIs (see Figure 9 for an example with dorsoposterior insula, and CANlab 2024 atlas ROIs that were downsampled to their second level (*labels_2*) then bilaterally merged to allow for adequate full-body pattern representation within 131 regions (Supplemental Figure 8). Classification was cross- validated using a leave-one-session-out approach, yielding a Hot-Warm SVM body site weight map for each fold of the data.

**Figure 9:**
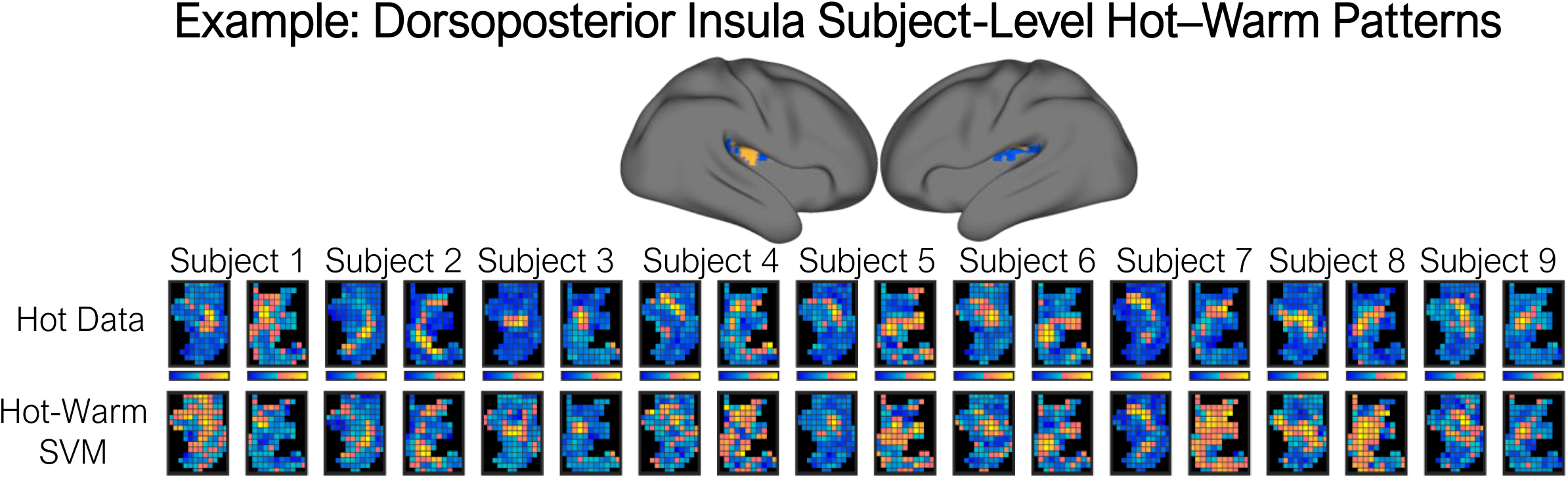
Example group-average SVM patterns in dorsal posterior insula (dpIns). Top: dpIns test data for the Hot (painful) condition for each participant. Bottom: SVM weight maps in dpIns for each participant. Pattern expression response values are subsequently compared across conditions (e.g., [Imagine - Warm]) and tested against zero at the group level (i.e., participant as a random effect) to establish significance. Follow-up tests assess significance within individual participants across sessions.

To assess the consistency of body site-specific pain representations, pattern expression was computed with the dot product for each body site’s Hot-Warm weight maps against that participant’s unsmoothed, run-level images across all conditions (Hot, Warm, and Imagine) and all body sites. For each ROI, a mixed-effects model was fit to predict pattern expression, with the Warm condition at the same body site as the intercept. Regressors included dummy-coded terms for Condition (Hot vs. Warm, Imagine vs. Warm), Body Site Comparison (Target vs. Other Body Site), their interaction, and participant-level nesting with random slopes and intercepts. A priori contrasts were specified as follows:

##### Conditional Averages

- Warm: Target Body Site (Intercept Model)
- Warm: Other Body Site – Warm: Target Body Site
- Hot: Target Body Site – Intercept
- Imagine: Target Body Site – Intercept
- Hot: Other Body Site – Hot: Target Body Site
- Imagine: Other Body Site – Imagine: TargetBody Site

##### Tests of Body Site-Selectivity

- Hot: Target Body Site – Hot: Other Body Site
- Imagine: Target Body Site – Imagine: Other Body Site
- Hot: Target Body Site – Warm: Other Body Site
- Imagine: Target Body Site – Warm: Other Body Site

##### Tests of Pain Body Site-Selectivity

- (Hot: Target Body Site – Hot: Other Body Site) – (Warm: Target Body Site – Warm: Other Body Site)
- (Imagine: Target Body Site – Imagine: Other Body Site) – (Warm: Target Body Site – Warm: Other Body Site)

A brain region was considered body site-selective in experienced or imagined pain if (1) it exhibited significant, positive pattern expression in the Hot or Imagine condition and (2) expression was significantly greater for the target body site than other body sites. The involvement was considered pain-specific if it also significantly differentiated from the Warm condition. Statistical significance was determined using false discovery rate (FDR) correction across regions (q < .05).

#### Session- and Run-Level Effects

To examine if repeated exposure might allow imagined pain to become more ‘pain-like’ over time, or conversely, might more clearly separate real versus imagined pain, we examined whether imagined pain increasingly changed the degree each brain region expressed nociceptive (Hot–Warm) representations with experience. Specifically, we tested for systematic changes in individualized, body-site–specific precision representations across sessions (practice/learning) and runs (within-session fatigue or habituation), using linear mixed-effects models predicting pattern expression from Session and Run with subject-specific random slopes. The degrees of freedom of these effects were Satterthwaite corrected, and no Session × Run interactions were observed.

#### Body Site Mapping

Trial-level gifti images for the cortex were first smoothed with a 6mm FWHM kernel, then subjected to robust regression to generate associated t-maps for each body site from each participant. The first hot-stimulation trial of each run was removed because of the outsized whole-brain activation intensity induced by these trials. The t-maps were averaged by body site within participant (∼5 session maps per body site per participant), before being averaged by body site across participants (9 participant maps per body site). The resulting eight body site t-maps were subjected to winner-take-all analysis, such that the highest positive t-value at each gifti vertex on the cortical surface across the eight maps took on the color of the respective body site.

#### Representational Similarity Analysis

Representational similarity matrices (RSMs) were computed using Spearman’s rho spatial correlations on unsmoothed run-level data for each participant. Images were averaged within each of the three conditions (Hot, Warm, and Imagine) across the eight body sites, resulting in 24 total conditions and a corresponding 24 × 24 RSM. This allowed us to assess representational similarity within each condition and across body sites, both within and between conditions, for each participant. To control for potential run-session-related confounds, cross-condition body site diagonals were recomputed while excluding runs within sessions that contained the target body site, as Hot, Warm, and Imagine conditions for a given body site were always collected within the same run. Participant-level RSMs were then averaged to create a 24 × 24 group-level RSM. Separate RSMs were generated for the whole brain, as well as for each CANlab 2024 ROI (Supplementary Figure 8). To evaluate overall similarity within and across conditions, we conducted a series of one-tailed t-tests across participants after applying Fisher’s *r*-to-Z transformation to address the non-normality of the rho statistic. Tests were conducted for each of the main conditions (hot, warm, and imagine vs. 0) and cross-conditions (hot and warm, hot and imagine, imagine and warm) reflecting our a priori hypothesis that there are voxelwise patterns that reflects each condition across body sites, and that these patterns recapitulate themselves positively across conditions. We also tested condition contrasts (e.g., Hot vs. Warm) as well as cross-conditional contrasts (e.g., Hot and Imagine vs. Hot and Warm). Body-site specificity was assessed by contrasting within-body-site with between-body-site similarity, with significance determined by false discovery rate (FDR) correction across parcels (q < .05). An ROI was determined to be related to pain imagination if cross-condition correlations for (Imagine, Hot) correlation exceeded both the (Hot, Warm) and (Imagine, Warm) correlations at an uncorrected threshold (p < .05 in each, one-tailed) and at least one surviving FDR correction (q < .05)..

## Data Availability

The fMRI data generated in this study are part of a registered clinical trial (ClinicalTrials.gov identifier NCT04653064; https://clinicaltrials.gov/study/NCT04653064) and are available under restricted access, as the full dataset is pending public release alongside a formal data descriptor. Upon release, the dataset will be deposited in OpenNeuro. Until then, access can be obtained by contacting the corresponding authors; requests will be responded to within four weeks, and access will be granted for research purposes consistent with participant consent. Source data underlying the figures are provided at https://github.com/canlab/2026_Sun_et_al_Pain_Imagination.

## Code Availability

We used the open-source software fMRIprep for image preprocessing and the CANLab Tools in Matlab to write custom code for data processing, which is available on GitHub at http://github.com/canlab. Analysis code used in this study is publicly available at https://github.com/canlab/2026_Sun_et_al_Pain_Imagination.

## Acknowledgements

We are grateful for assistance from Terry Sackett, Maryam Amini, Melanie Kos, Stephanie Sun, Eilis Murphy, Samuel Bergerson, Kristoffer Mansson, Sreekar Kasturi, Jason Davis, and Charles Mazof for assistance in data collection for this study. We thank Byeol Kim, Ben Graul, Li-Bo Zhang, and Zhaoxing Wei for comments on the manuscript, and Luke Chang and Marianne Reddan for insightful discussion.

## Author Contributions Statement

M.S. and T.D.W. conceived and designed the study. M.S., C.B., M.H., and S.S. collected the data. M.S. analyzed the data. M.S. wrote the original draft. T.D.W. supervised the project and acquired funding. M.S., K.B., H.J., B.P., and T.D.W. reviewed and edited the manuscript. All authors approved the final manuscript.

---

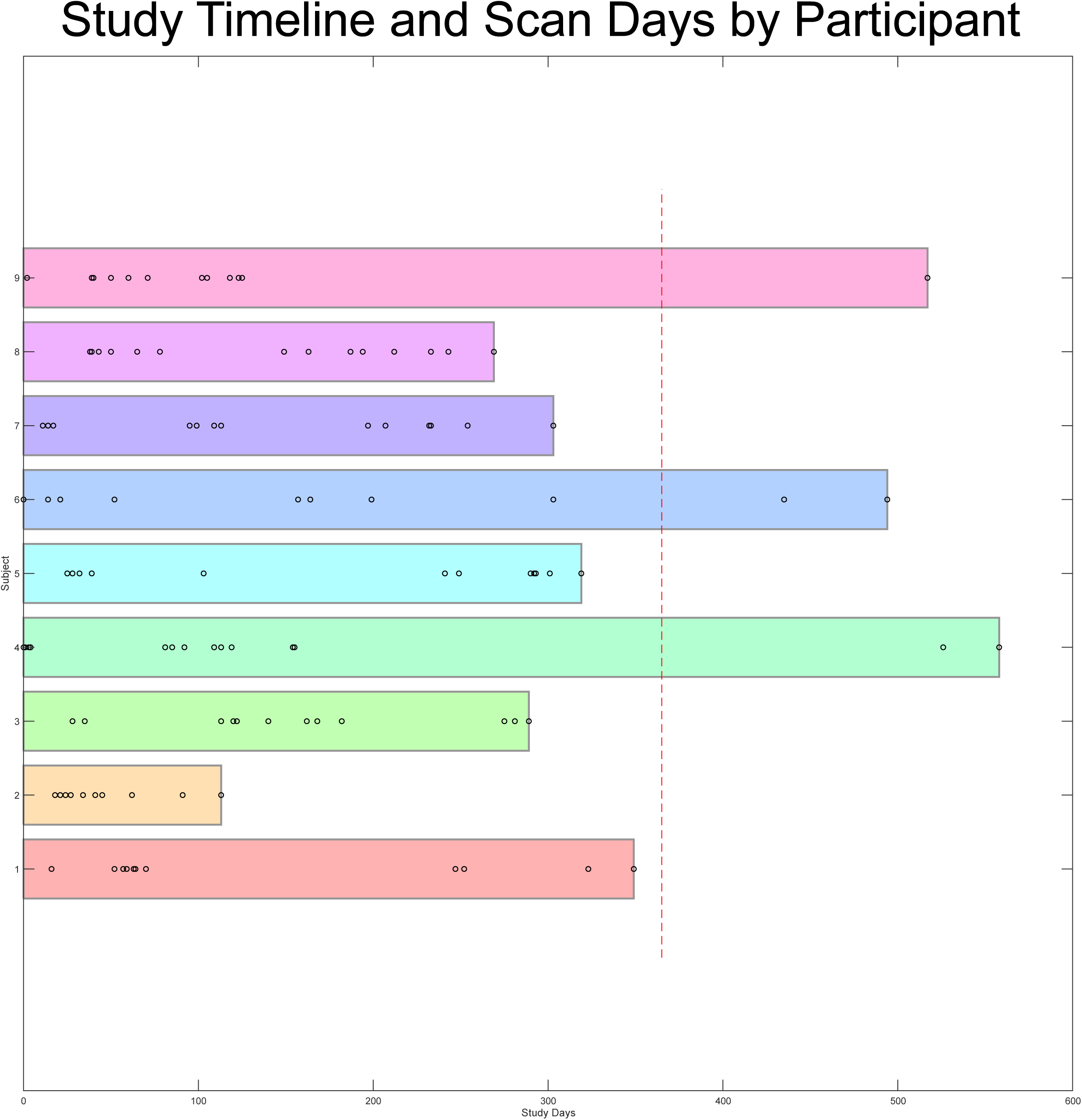

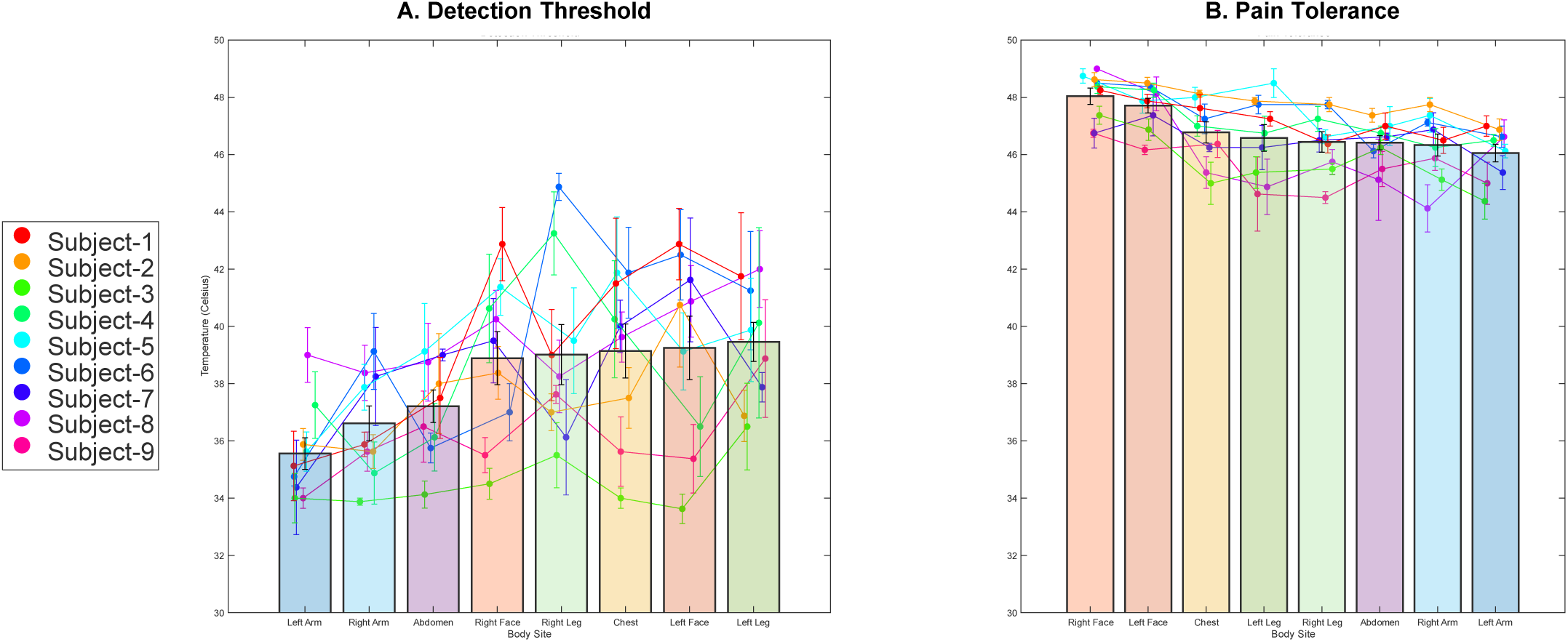

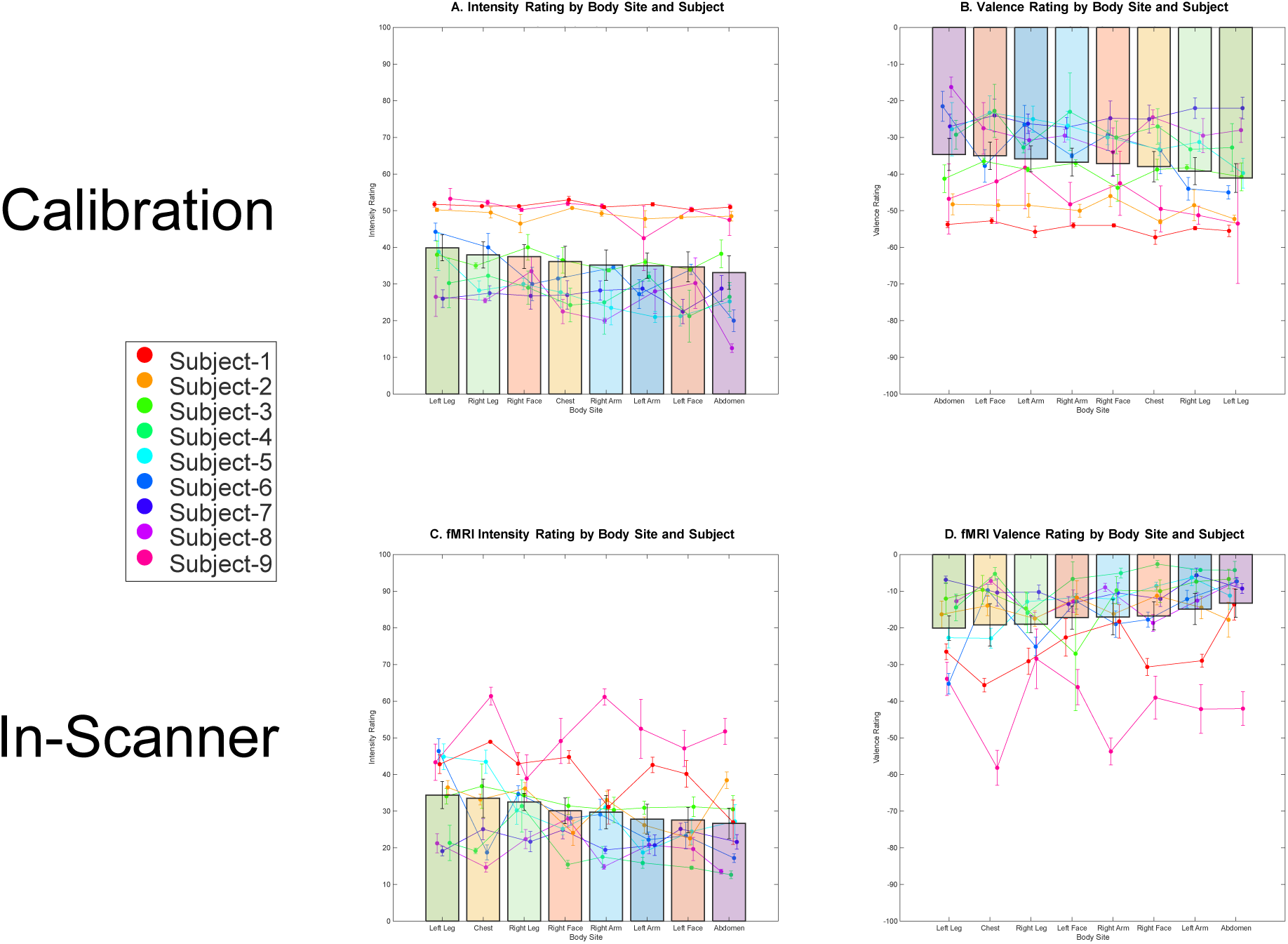

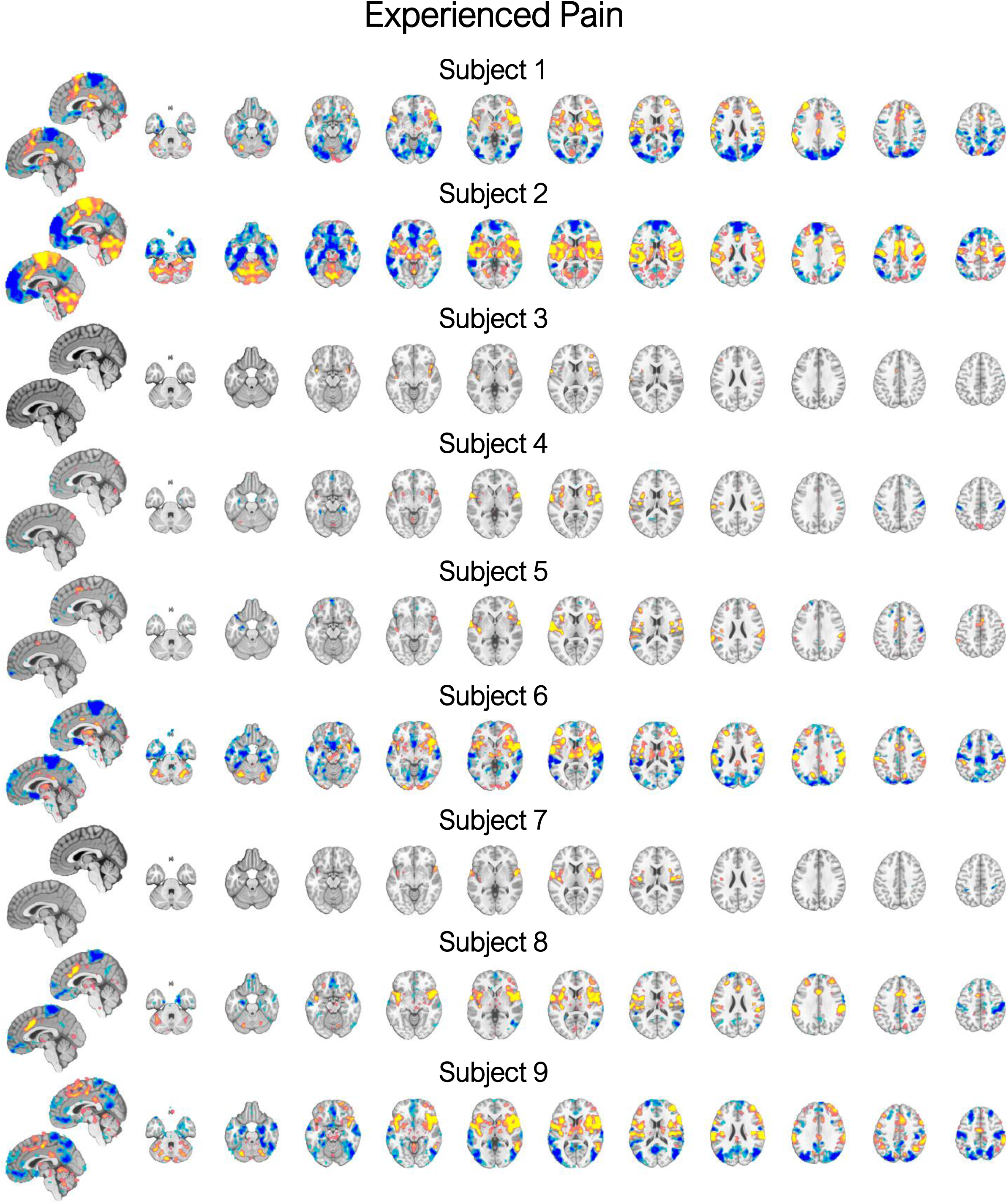

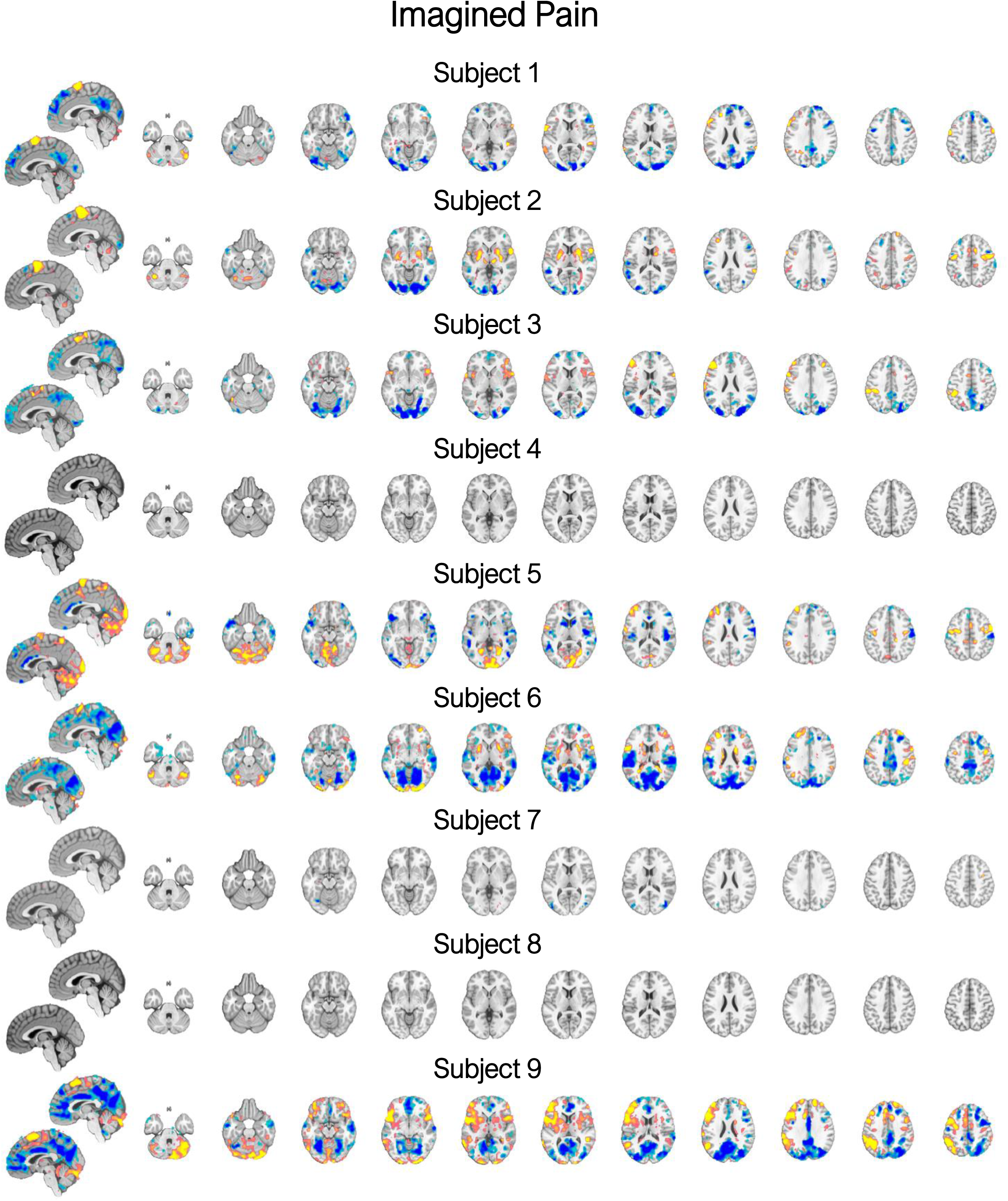

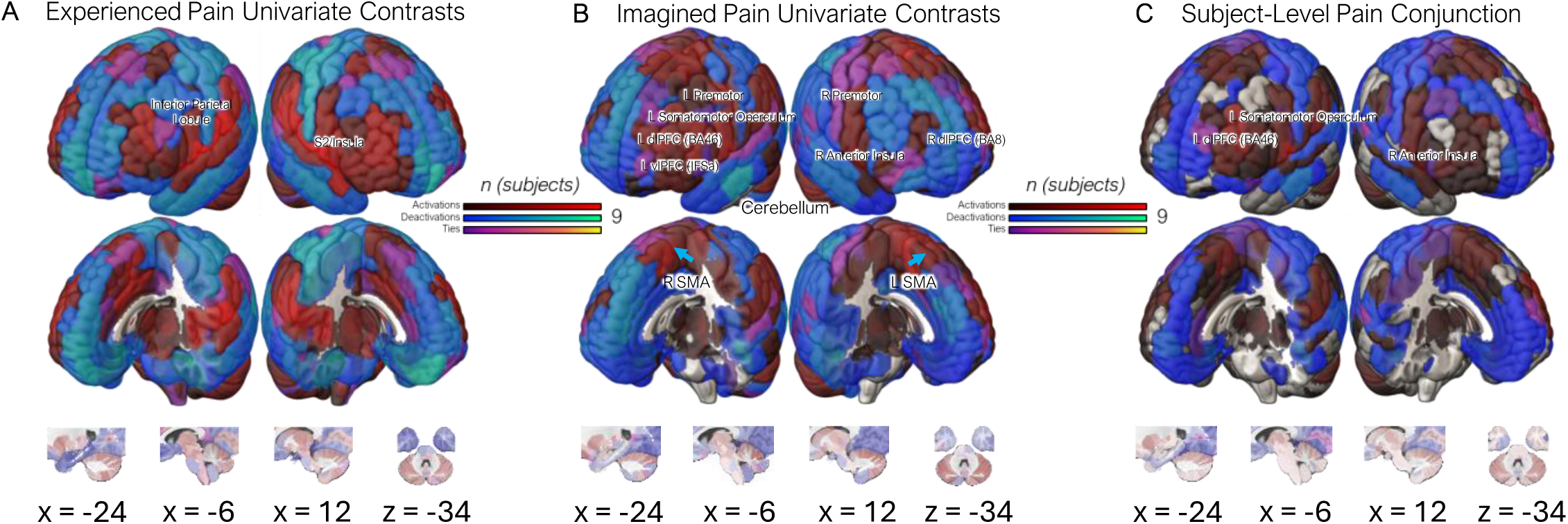

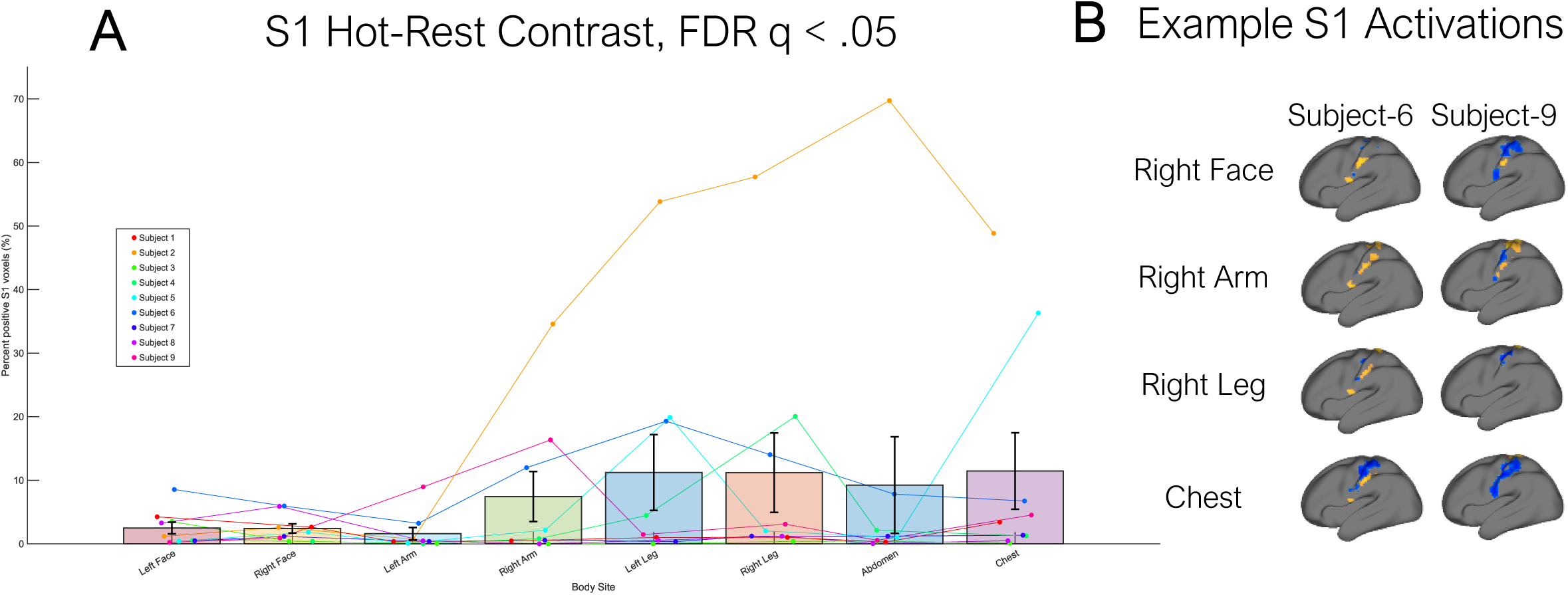

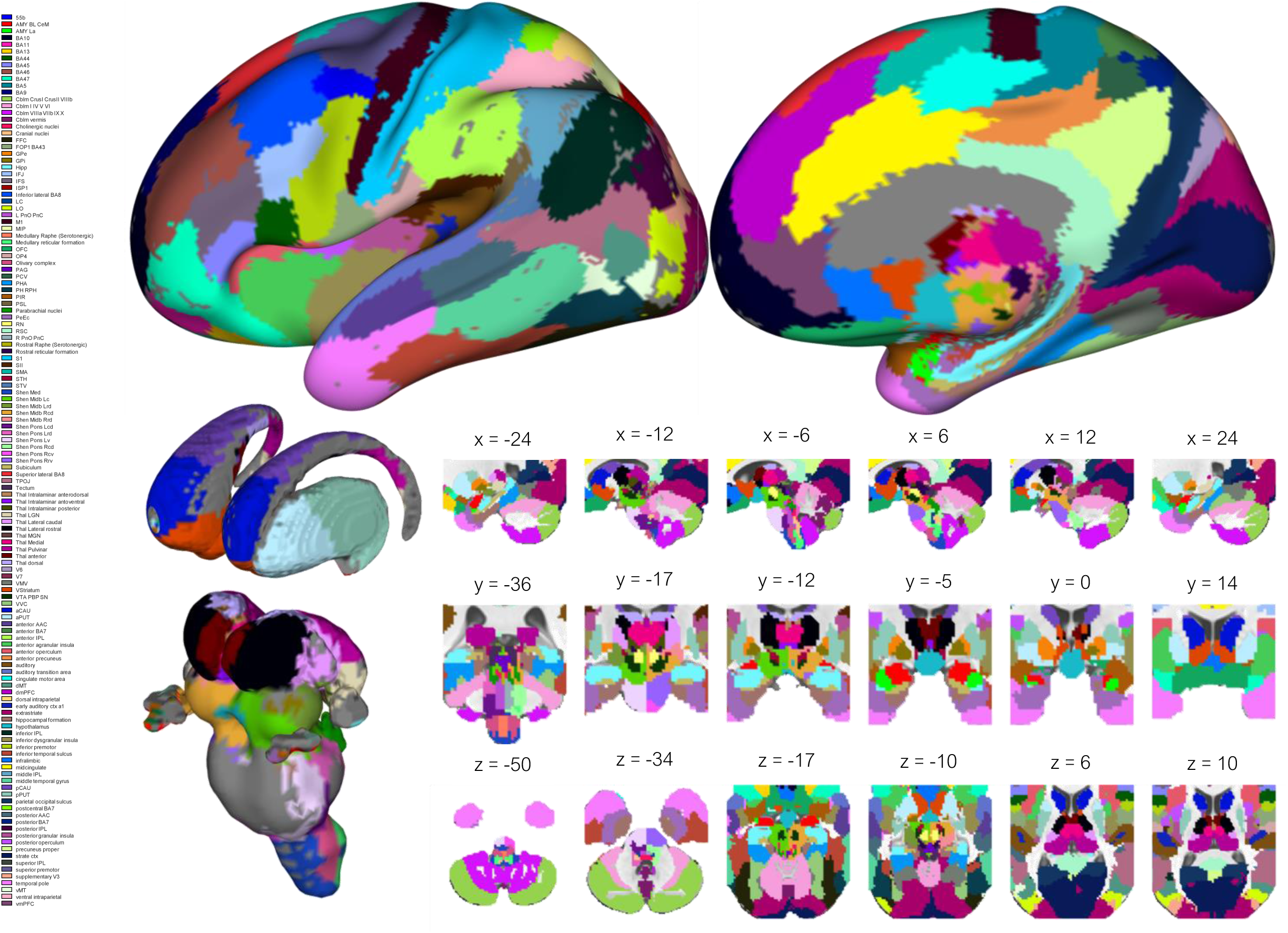

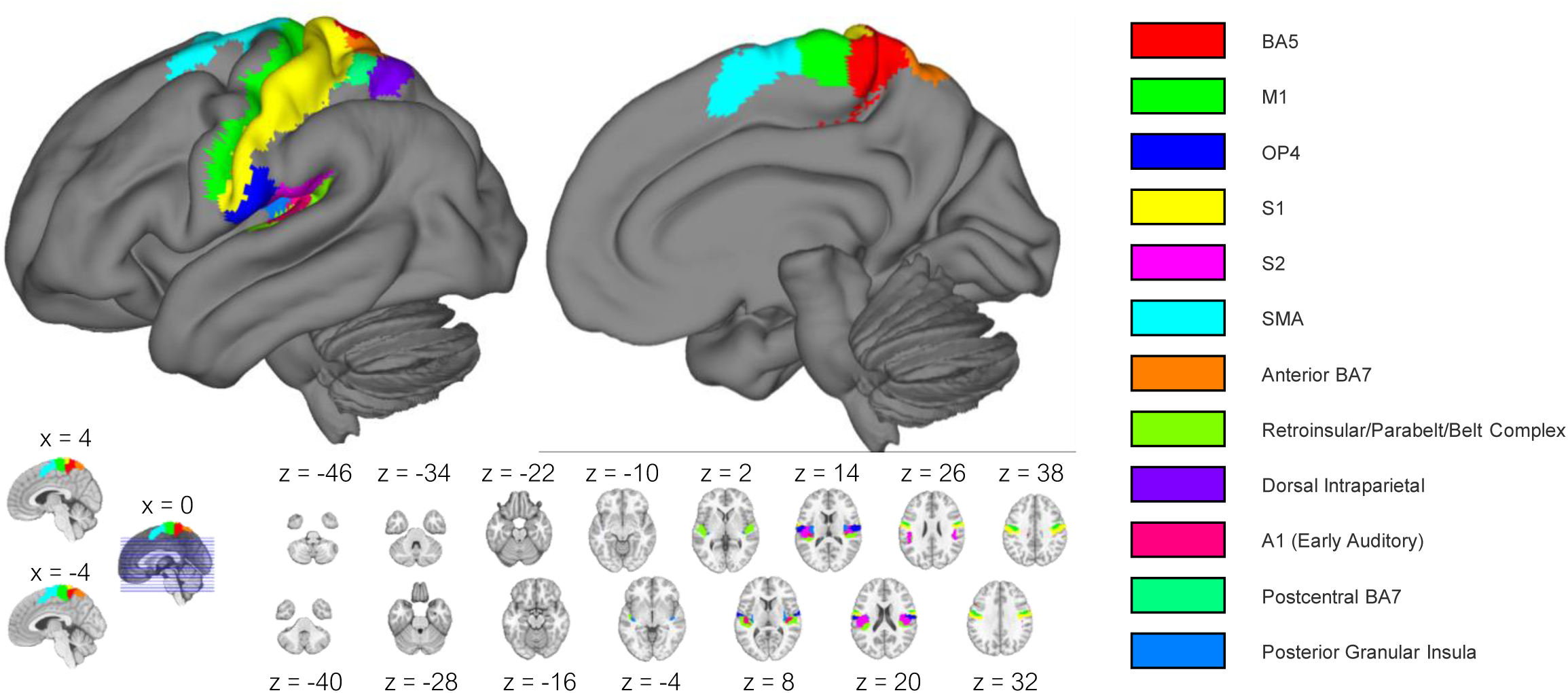

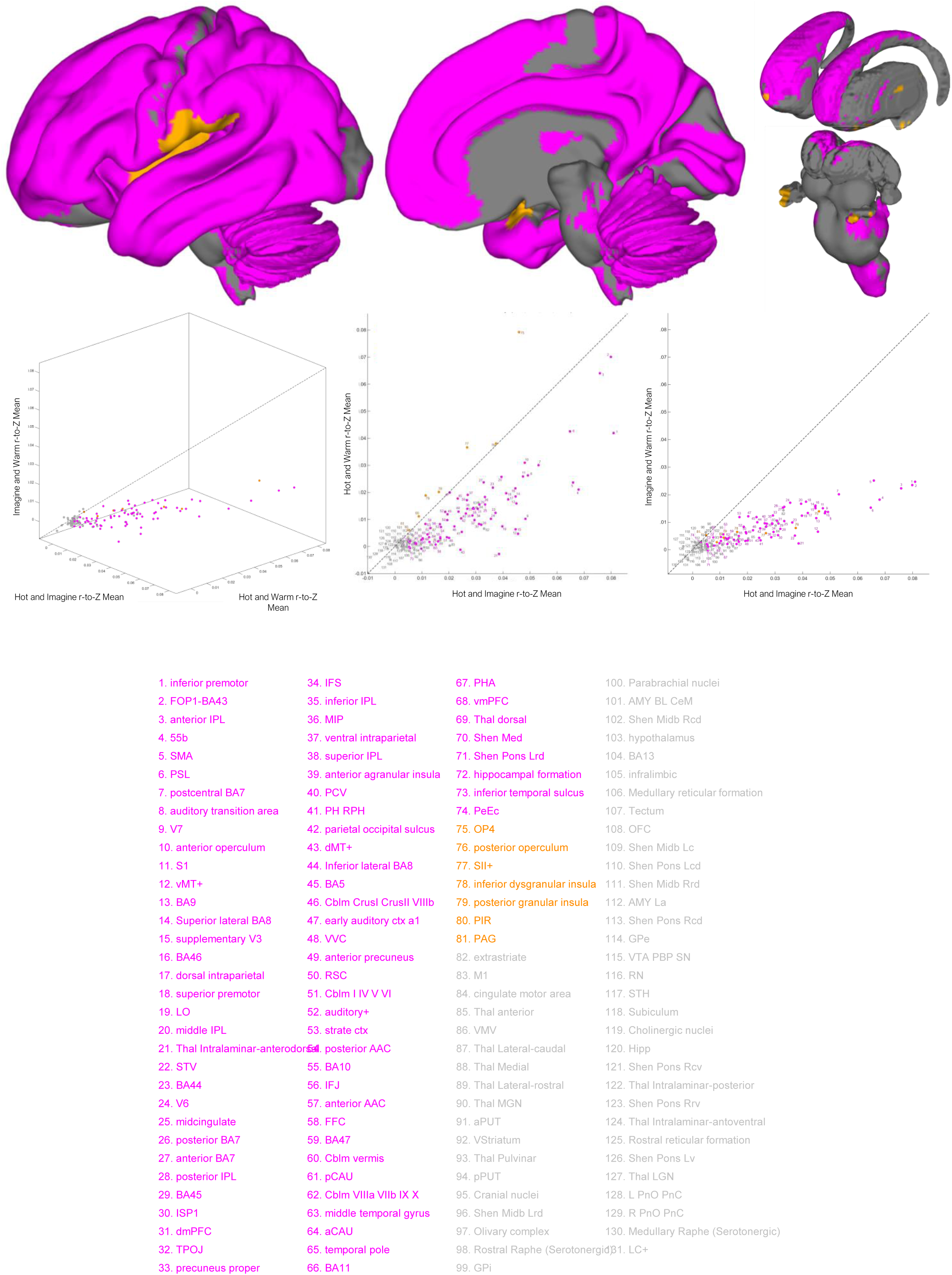

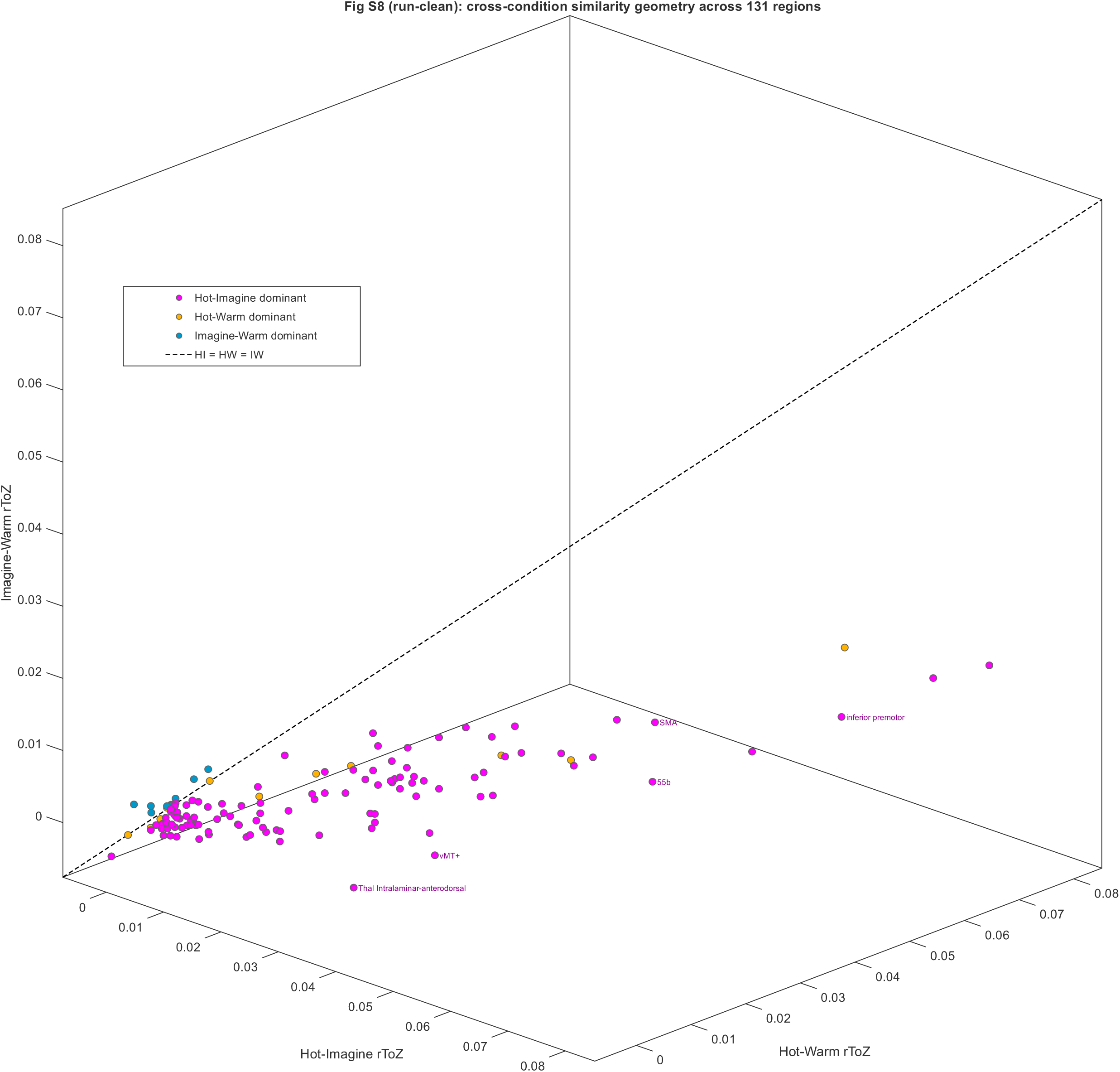

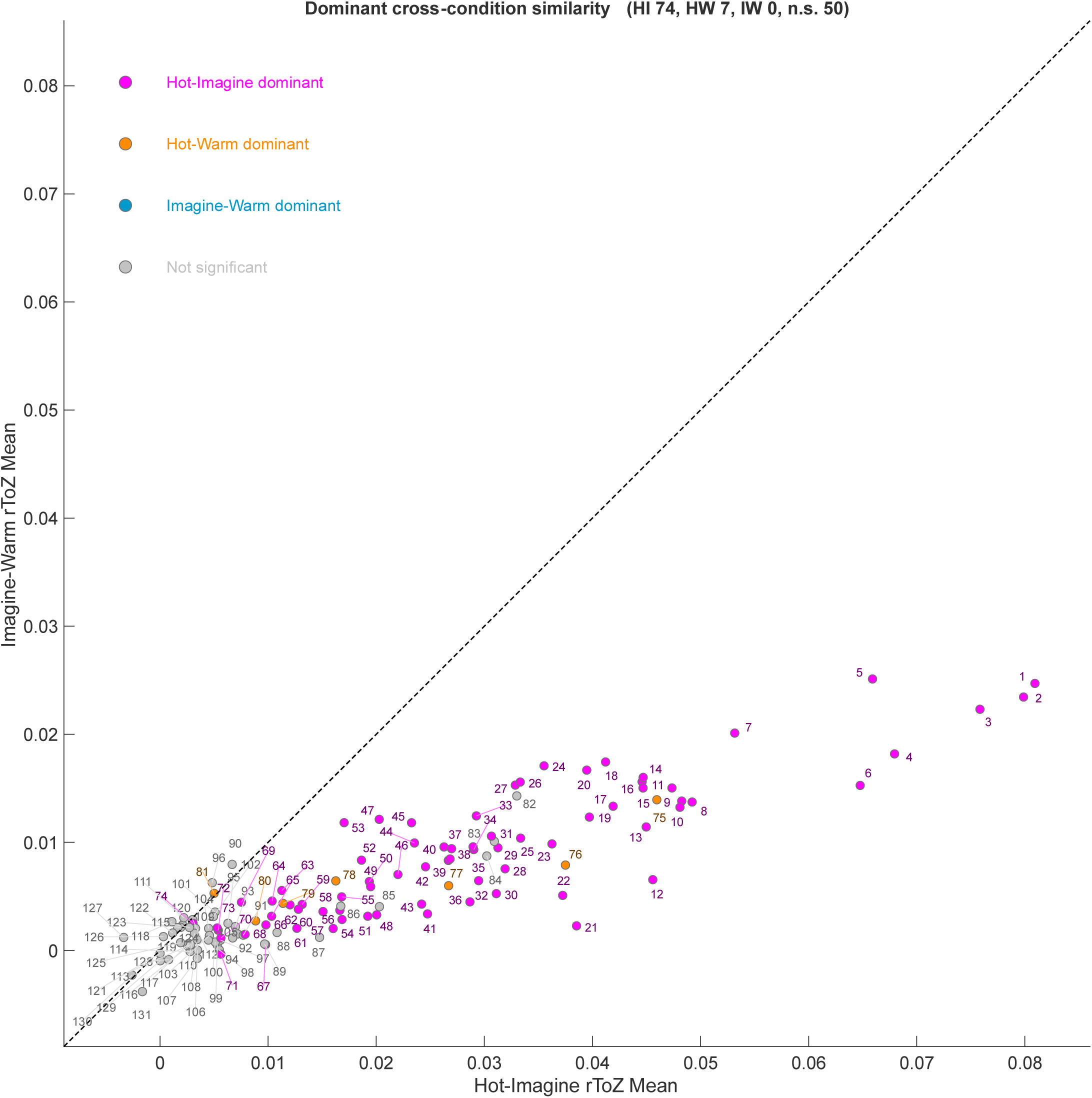

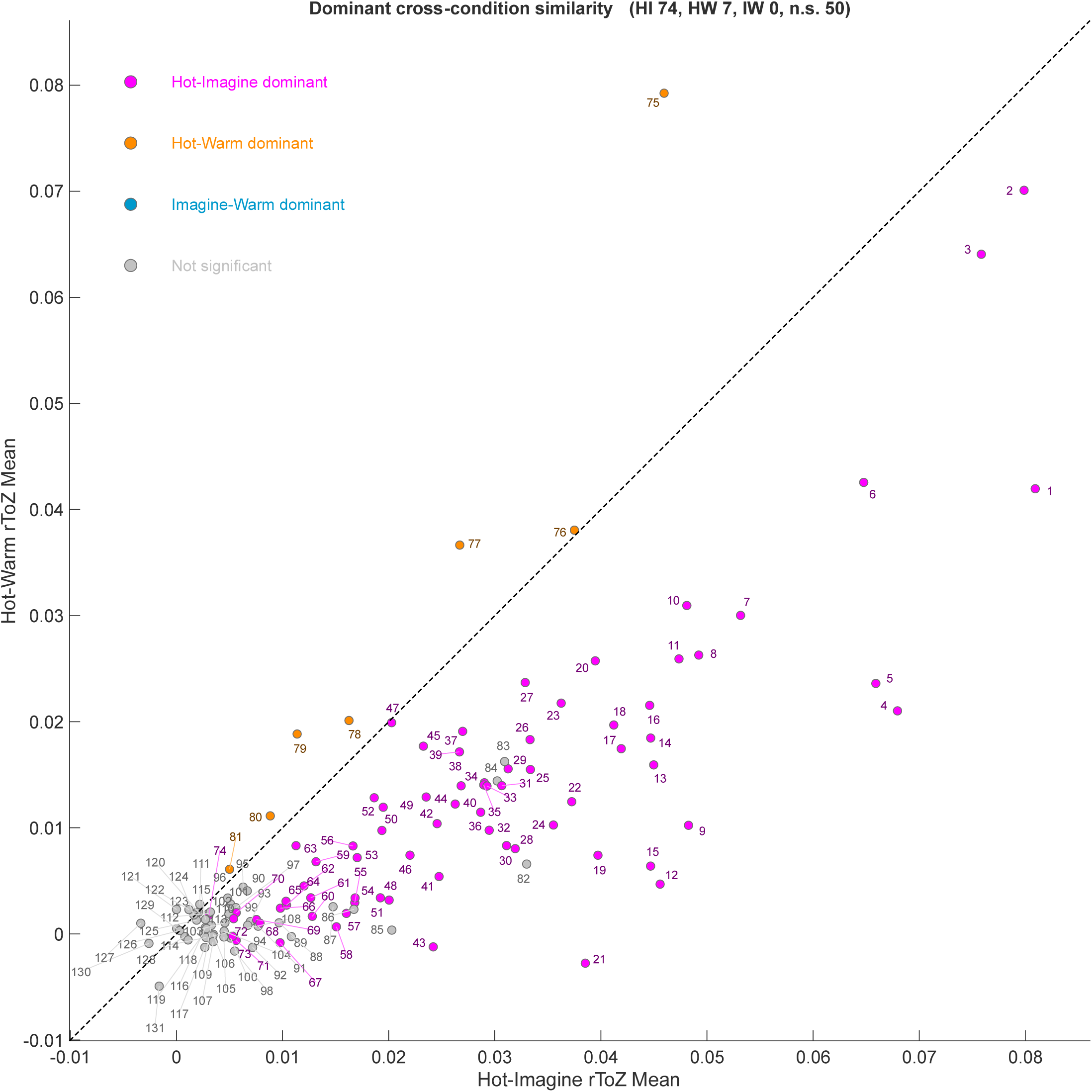

