## Supplemental Materials for "Precision Functional Mapping of Imagined and Experienced Pain"

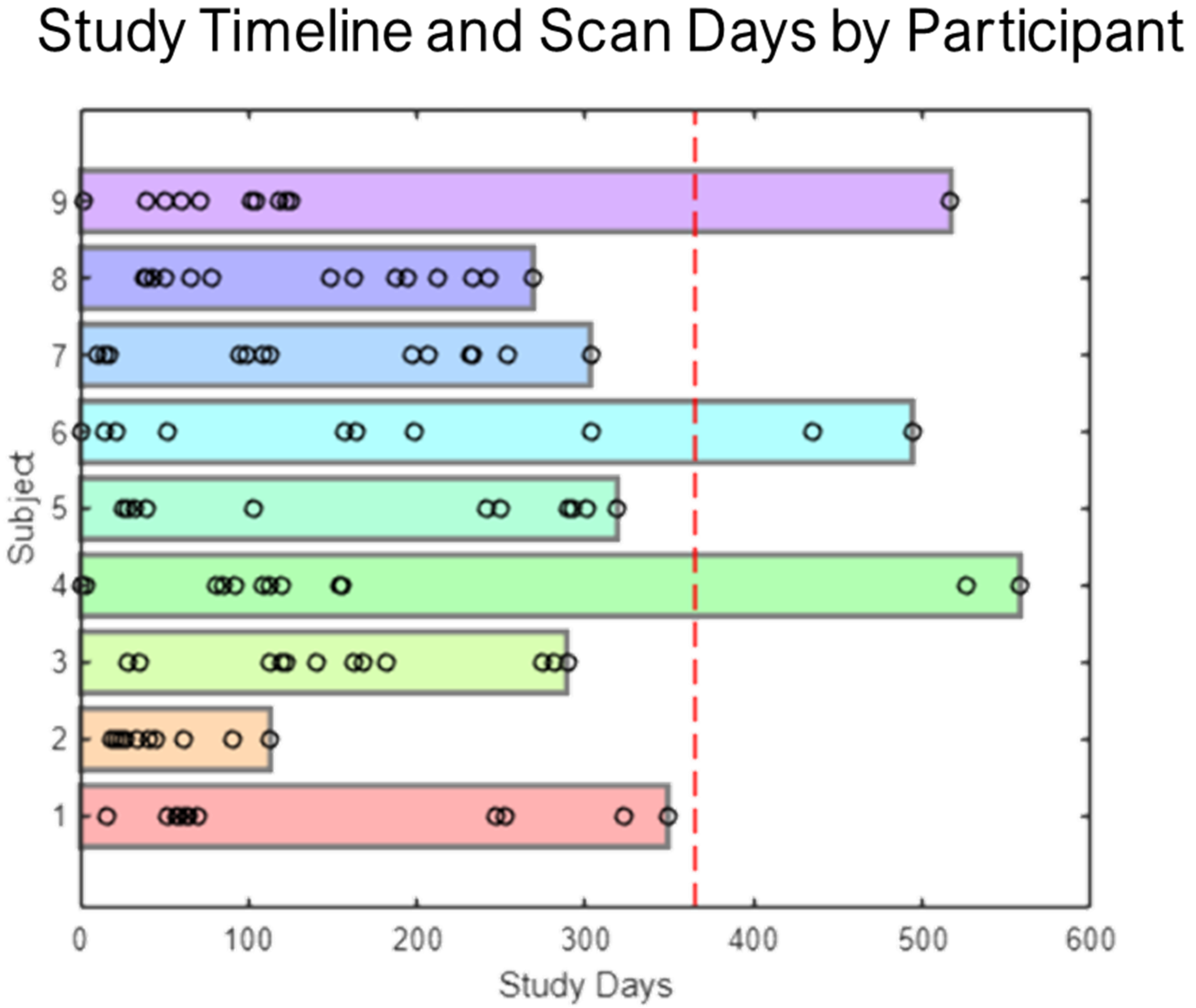


**Supplemental Figure 1: Study timeline and scan days by participant.** Horizontal bars indicate the total calendar span (in days) over which each participant was involved in the study. Circles denote individual fMRI scanning days. Participants were scheduled for 10 scanning sessions; however, some participants completed additional sessions due to re-scanning, whereas others completed multiple sessions on the same day, resulting in different numbers of distinct scan days. The dashed red line indicates one year (365 days) as a point of reference.

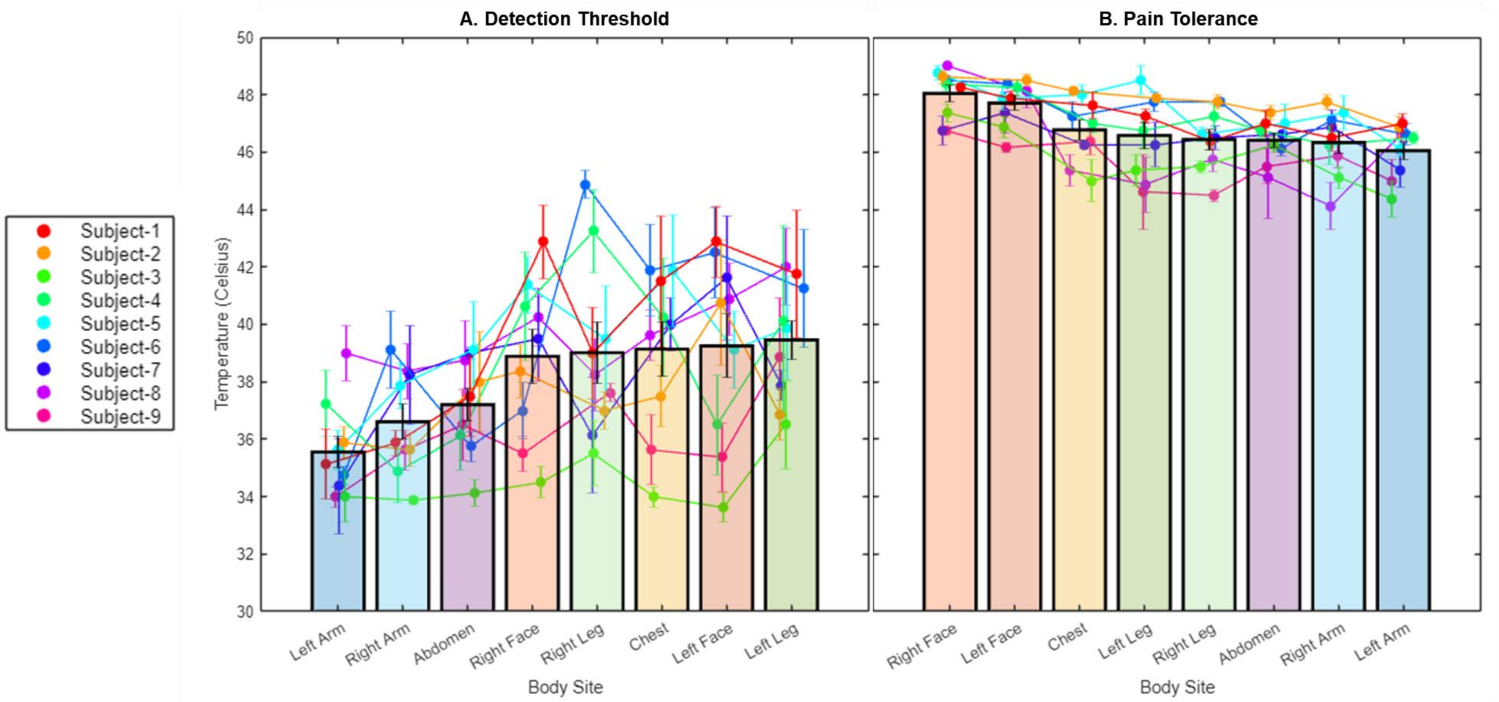


**Supplemental Figure 2: Participant-level thermal detection thresholds and pain tolerance across body sites.** Participant-level (A) thermal detection thresholds and (B) pain tolerance at each body site. Bars show group mean ± SEM across n = 9 participants; dots show individual participants.

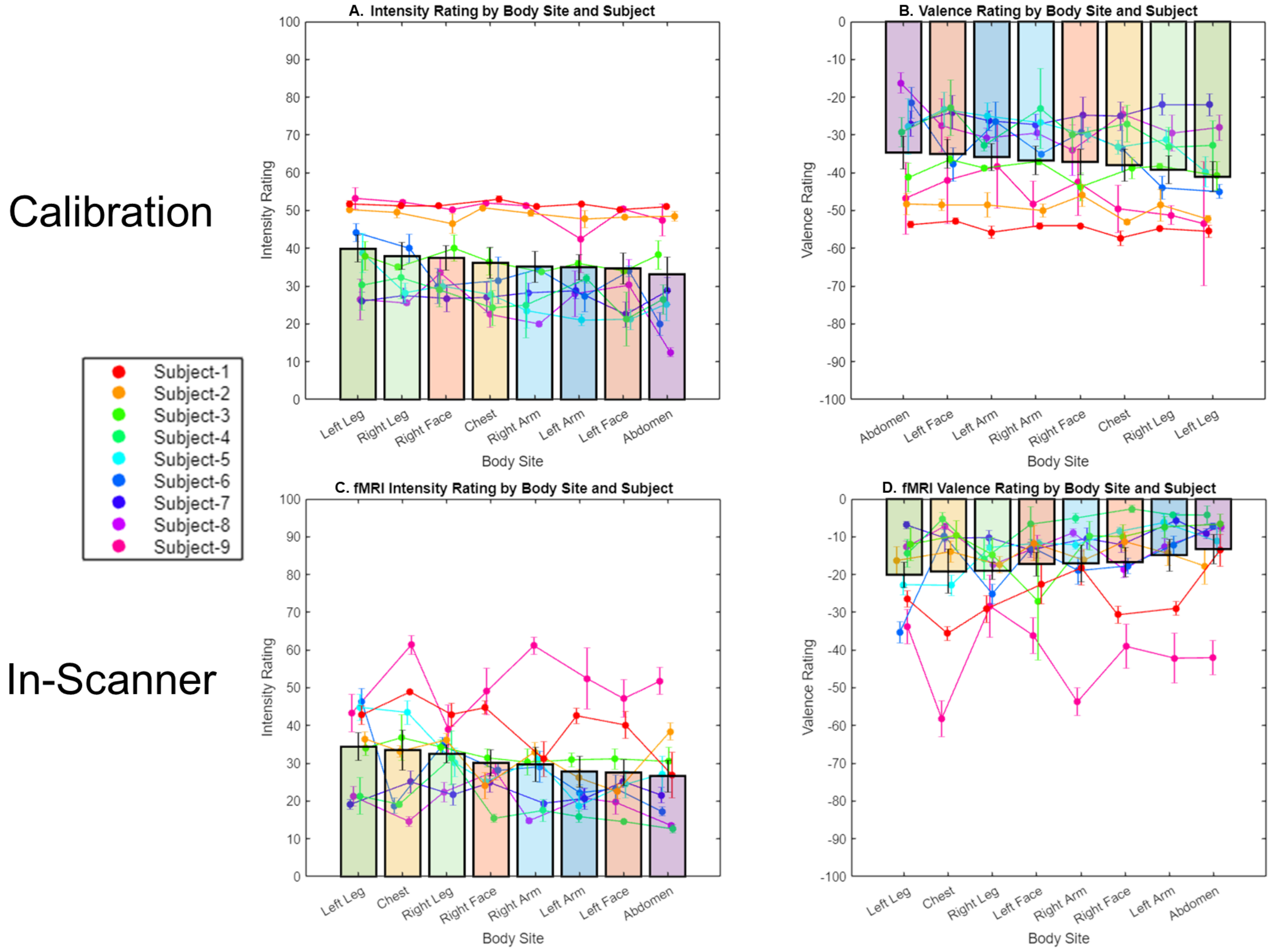


**Supplemental Figure 3: Participant-level intensity and valence ratings during calibration and scanning.** Participant-level ratings during calibration for (A) intensity and (B) valence, and during fMRI scan-sessions for (C) intensity and (D) valence ratings. Bars show group mean ± SEM across n = 9 participants; dots show individual participants.

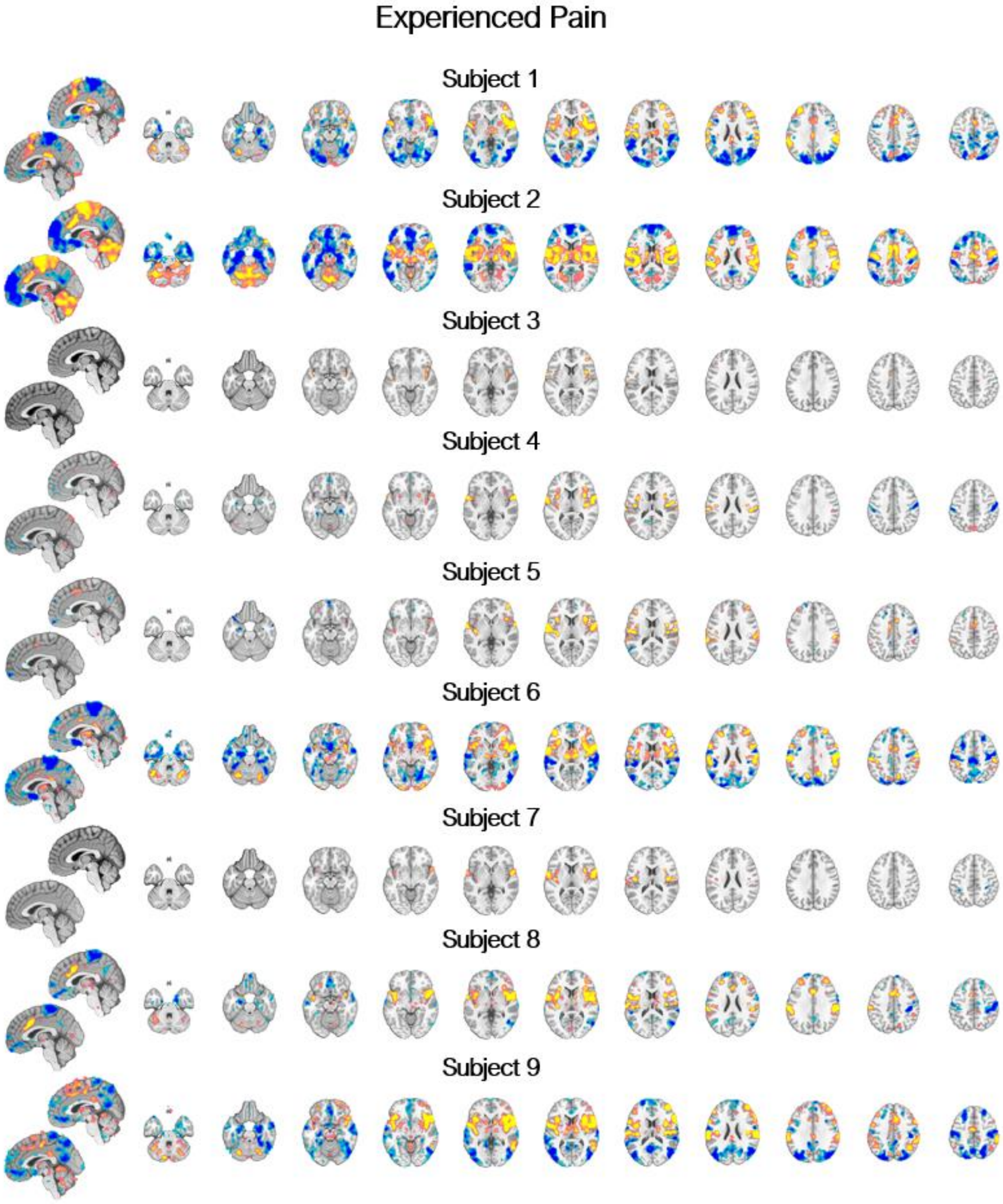


**Supplemental Figure 4: Participant-level experienced pain contrast (hot–warm).** Participant-level experienced pain contrast (hot-warm). FDR q < .05.

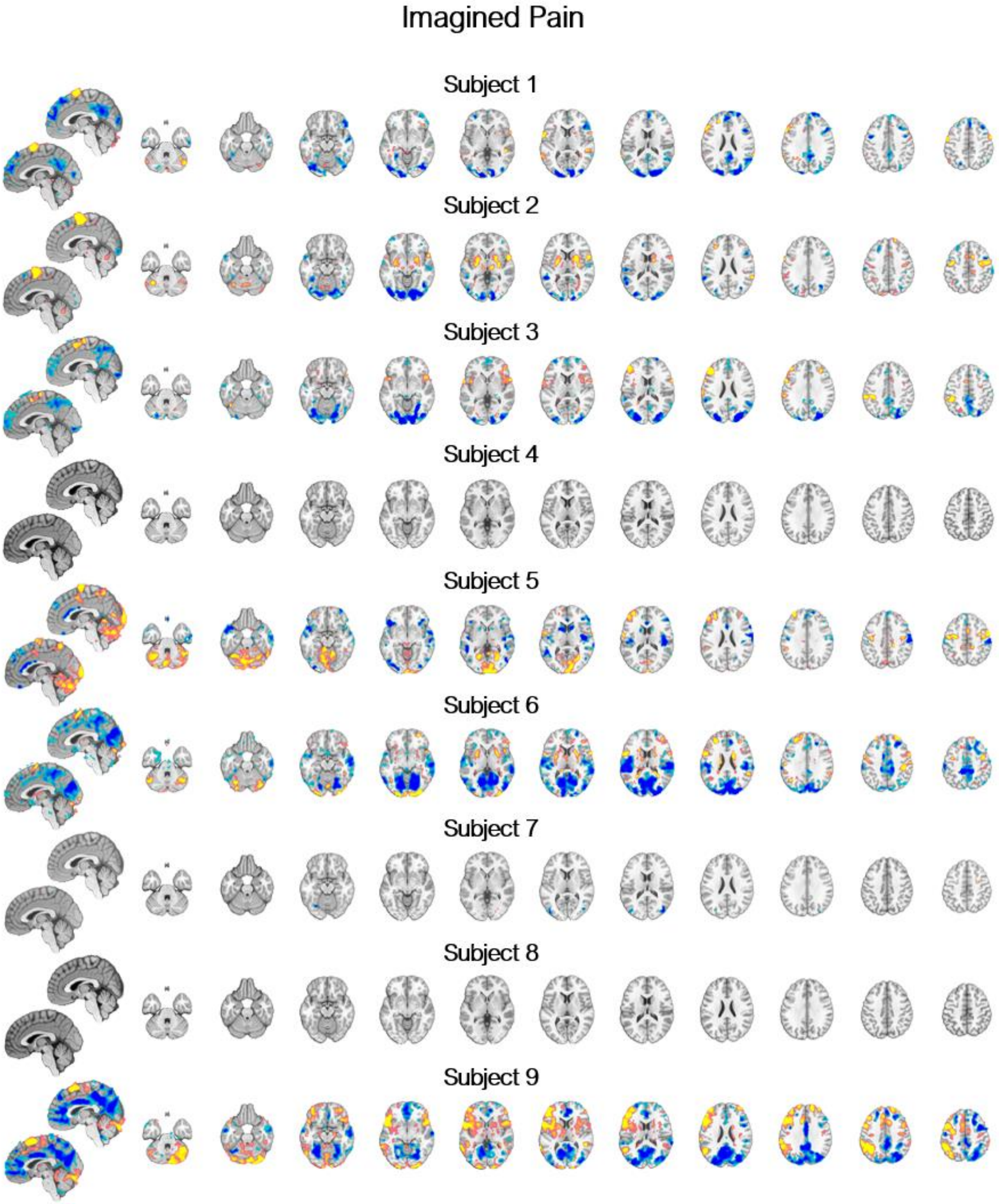


Supplemental Figure 5: Participant-level imagined pain contrast (imagine–warm). Participant-level imagined pain contrast (imagine-warm). FDR q < .05.

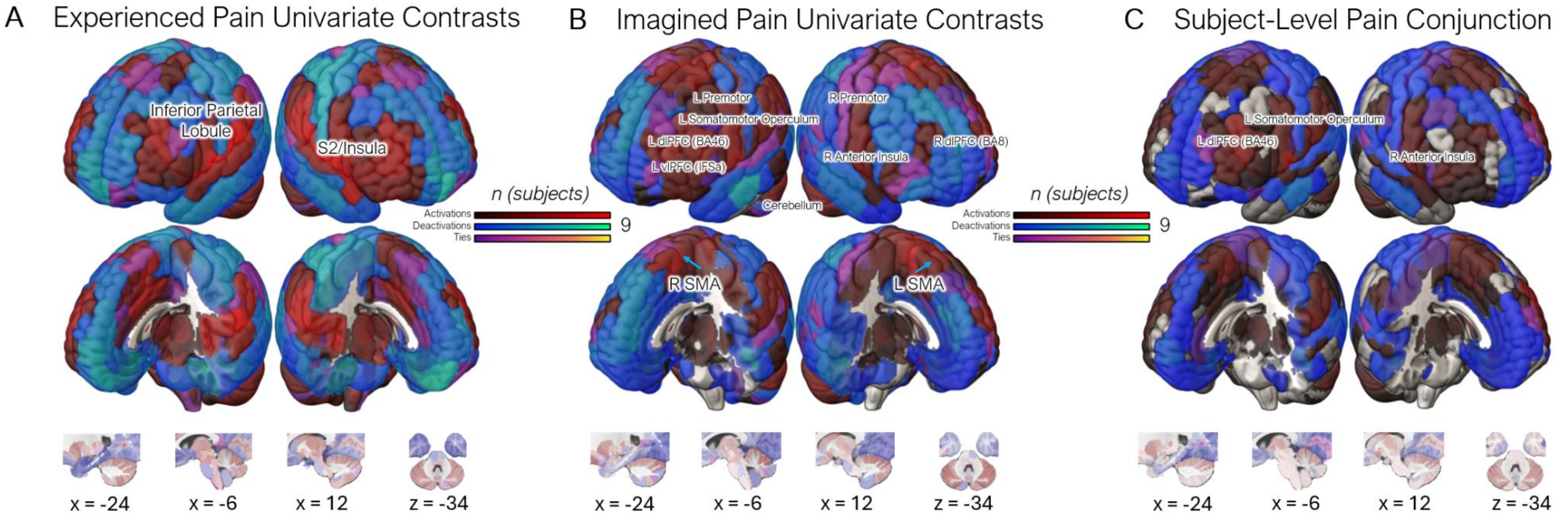


Supplemental Figure 6: Count maps of FDR-thresholded participant-level univariate activations of regions. Each region required a coverage threshold of 5% of total region voxels activated or deactivated by the participant to be counted toward the respective counts of "activated" or "deactivated" regions FDR $q < .05$. Regions are colored in accordance to whether they are activated or deactivated by the majority of participants, or if there are an equal number of participants that are activating and deactivating.

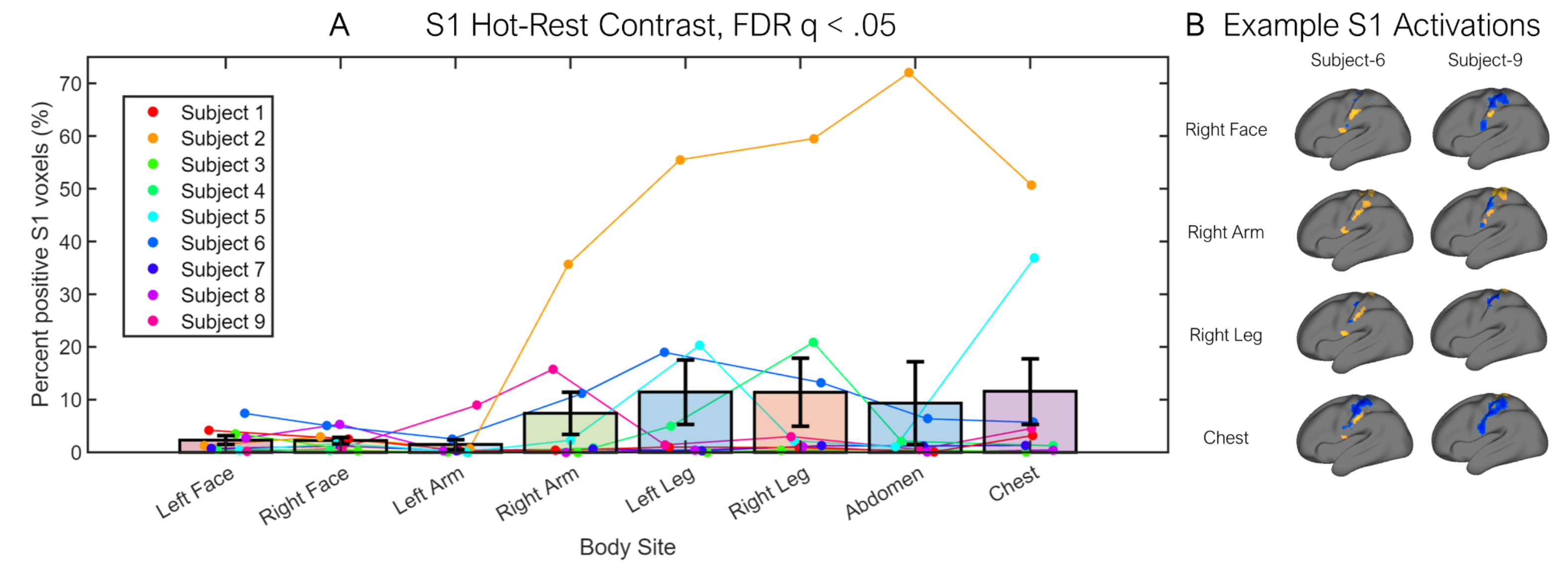


**Supplemental Figure 7: Body-site–specific S1 activation and inter-individual variability.** (A) Percentage of S1 voxels showing significant activation for the Hot–Rest contrast (FDR q < 0.05) is plotted for each body site. Bars indicate the group mean ± SEM across subjects, with individual subjects shown as colored points connected across body sites (colors denote subjects). S1 activation was observed in all subjects for the left face, right face, right leg, abdomen, and chest, whereas activation was observed in 8/9 subjects for the left arm and left leg, and in 7/9 subjects for the right arm. (B) Subject-level surface maps for example subjects 6 and 9 illustrating focal, body-site–specific S1 activation for the Hot–Rest contrast (FDR q < 0.05), with substantial inter-individual variability in the spatial extent of activation. Bars show group mean ± SEM across n = 9 participants.

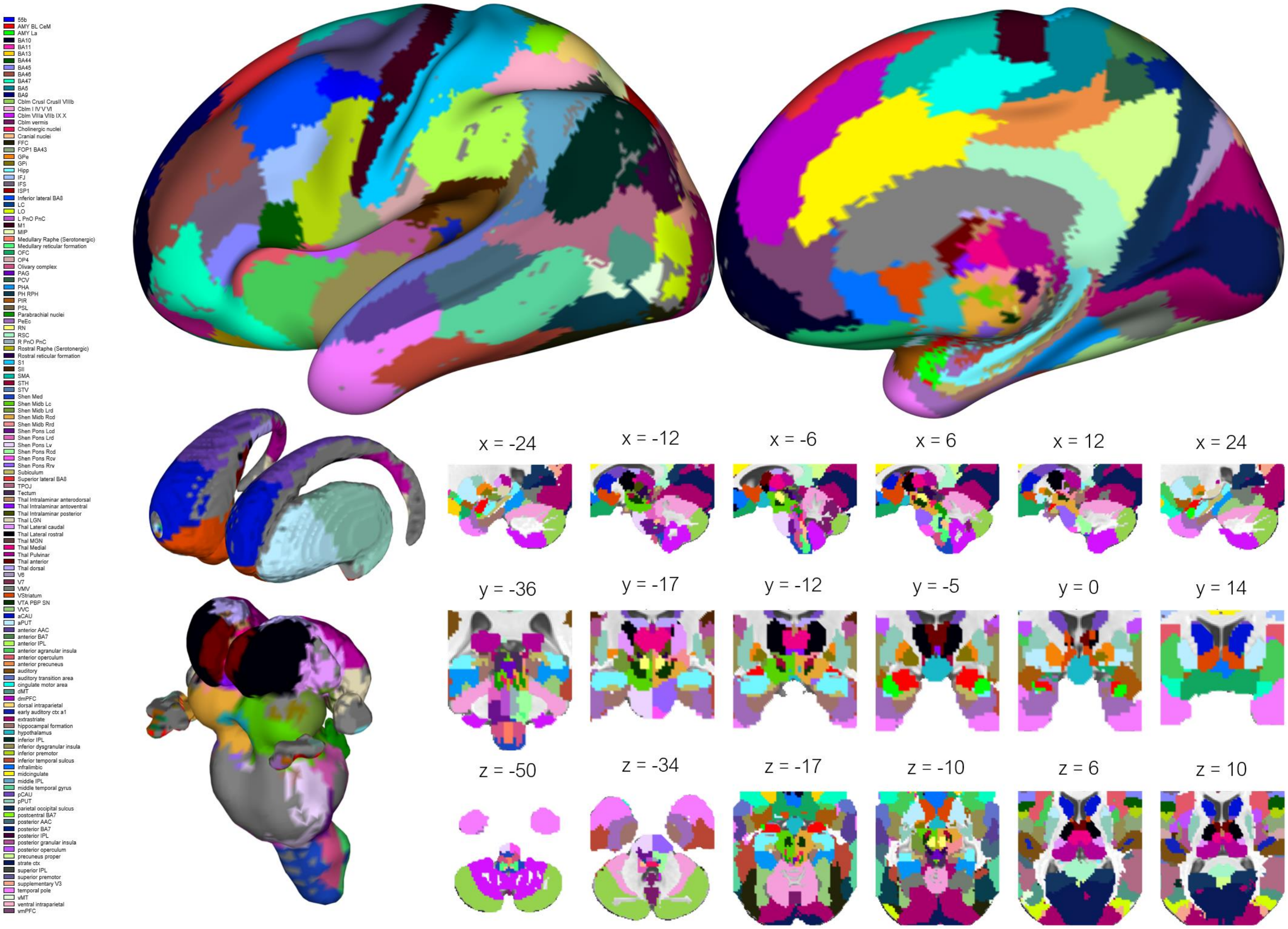


Supplemental Figure 8: The 131 Region CANLab 2024 Atlas.

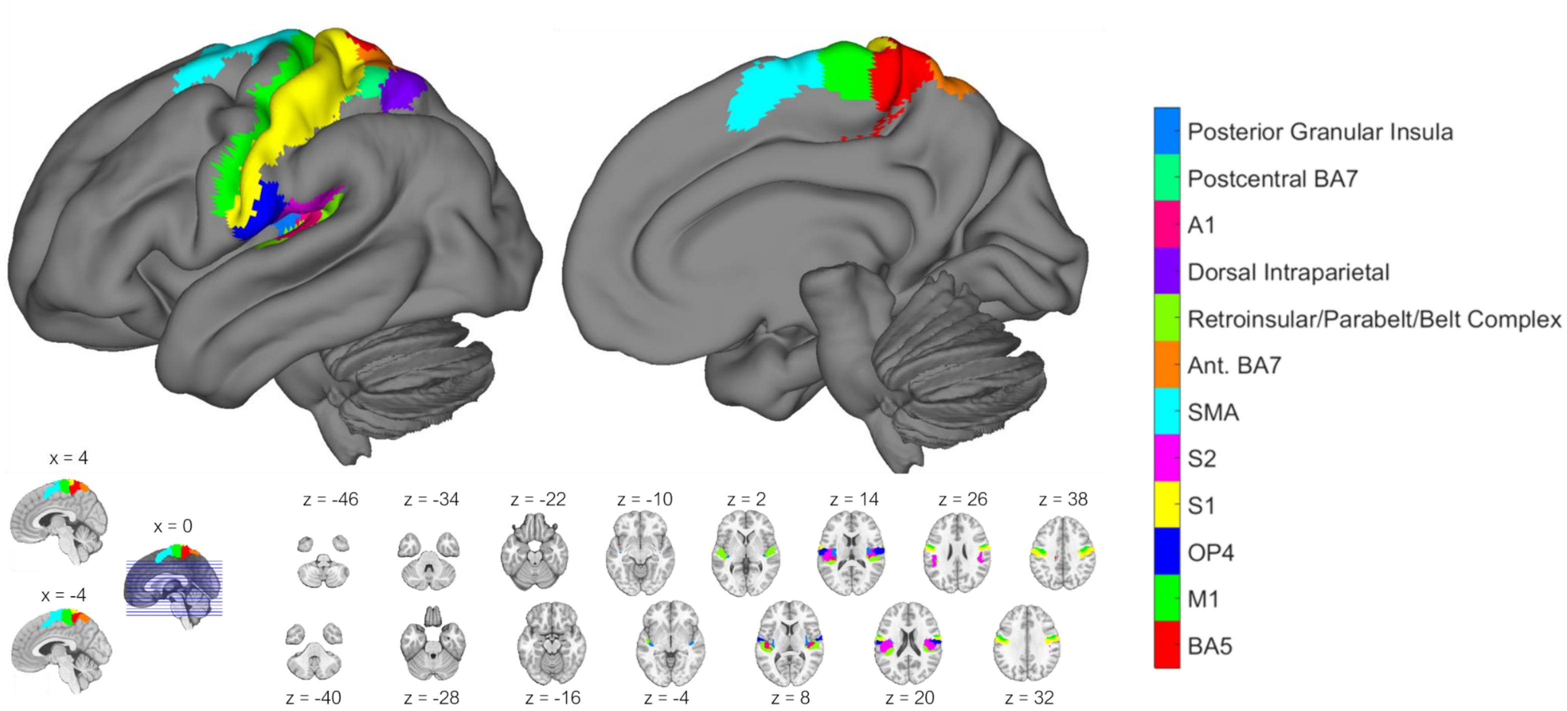


**Supplemental Figure 9: FDR-corrected nociceptive body site selective parcels from the CANLab 2024 Atlas.** BA: Brodmann Area. VTA: Ventral Tegmental Area. LGN: Lateral Geniculate Nucleus. IL: Intralaminar. SMA: Supplementary Motor Area. S2: Secondary Somatosensory Cortex. S1: Primary Somatosensory Cortex. OP4: Opercular Area 4. M1: Primary Motor Cortex.

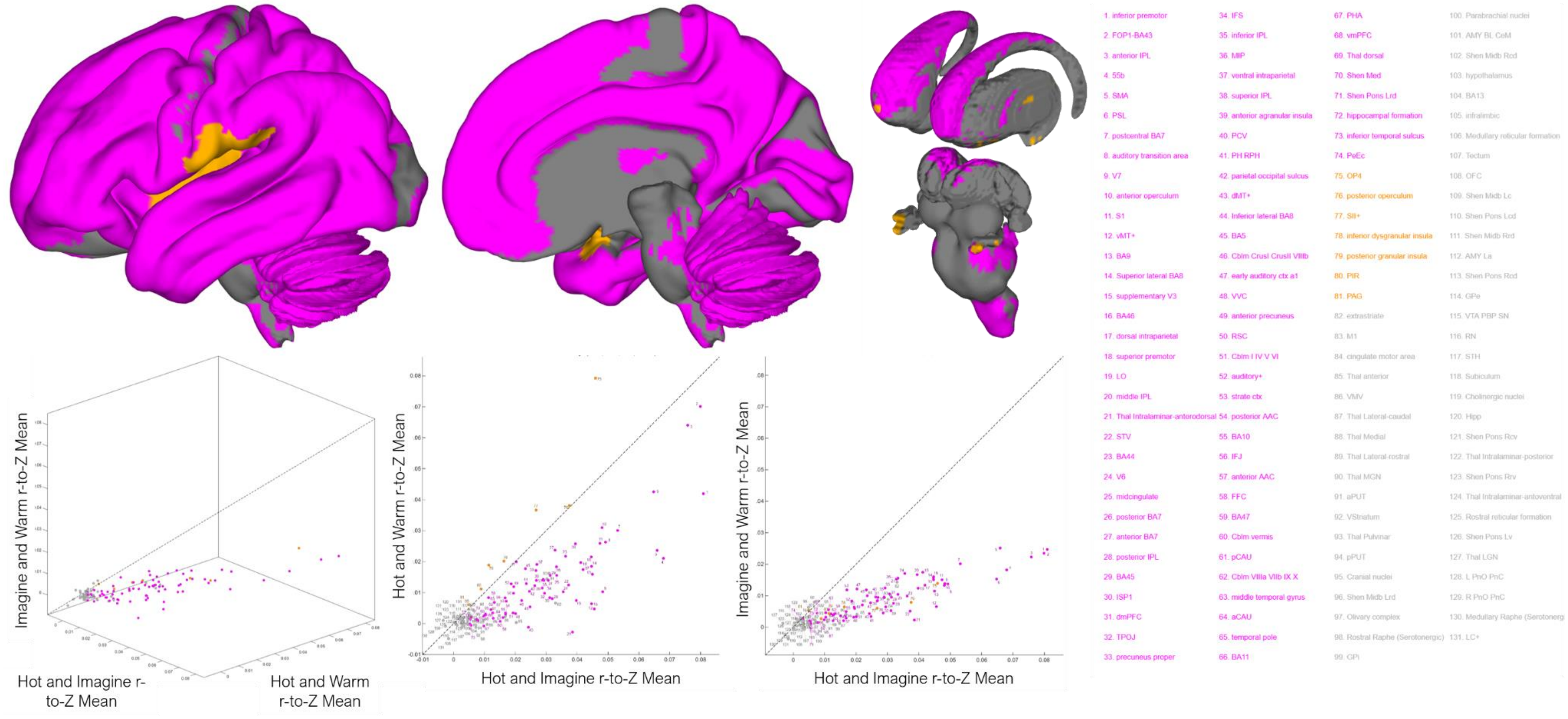


**Supplemental Figure 10: Regional dominance of cross-condition representational similarity.** Regions were colored by their highest significant condition-pair similarity. A region is colored by that dominant pair only if its dominant similarity was itself significantly greater than zero across participants (one-sample t-test, two-tailed $P < .05$); otherwise it is shown in gray. In most parcels (74/131), the dominant, significant similarity was between the Hot and Imagine conditions (magenta), indicating that imagined-pain activity patterns most closely resembled those evoked by experienced pain. Seven parcels (amber) were instead most similar between Hot and Warm. These included the periaqueductal gray (PAG), piriform cortex (PIR), and a cluster of opercular–insular somatosensory regions (SII+, OP4, posterior operculum, inferior dysgranular insula, and posterior granular insula). This is consistent with experienced thermal input dominating these early somatosensory areas. No region showed a significant dominant similarity between Imagine and Warm. In the scatter plots, each point is one region colored by its dominant significant pair, and the dashed diagonal marks the identity line, along which the two cross-condition similarities on the axes (or, in the three-dimensional plot, all three) are equal. Gray regions are those whose highest cross-condition similarity did not reach statistical significance.

Supplemental Table S1. Calibration temperatures and rating metrics

| Body Site | Detection Threshold | | Pain Tolerance | | Calibration Valence | | Calibration Intensity | | Scanner Valence | | Scanner Intensity | |
|---|---|---|---|---|---|---|---|---|---|---|---|---|
| | M | SE | M | SE | M | SE | M | SE | M | SE | M | SE |
| Left Face | 39.25 | 1.11 | 47.71 | 0.26 | -35.00 | 4.71 | 34.68 | 2.62 | -17.19 | 3.14 | 27.57 | 3.43 |
| Right Face | 38.89 | 0.93 | 48.04 | 0.29 | -37.14 | 4.20 | 37.47 | 2.77 | -16.72 | 3.84 | 30.10 | 3.52 |
| Left Arm | 35.56 | 0.55 | 46.06 | 0.31 | -35.83 | 3.56 | 35.00 | 3.01 | -14.85 | 4.22 | 27.83 | 4.08 |
| Right Arm | 36.61 | 0.61 | 46.33 | 0.38 | -36.75 | 3.21 | 35.17 | 2.16 | -17.04 | 4.83 | 29.72 | 4.54 |
| Left Leg | 39.46 | 0.69 | 46.58 | 0.46 | -41.06 | 4.61 | 39.89 | 3.31 | -20.07 | 3.35 | 34.39 | 3.70 |
| Right Leg | 39.01 | 1.05 | 46.44 | 0.36 | -39.19 | 2.93 | 37.94 | 1.99 | -19.01 | 2.29 | 32.51 | 2.35 |
| Chest | 39.14 | 0.94 | 46.78 | 0.37 | -37.97 | 3.38 | 36.14 | 3.04 | -19.19 | 5.81 | 33.49 | 5.24 |
| Abdomen | 37.21 | 0.57 | 46.42 | 0.25 | -34.64 | 4.27 | 33.14 | 2.87 | -13.27 | 3.84 | 26.64 | 4.20 |

**Note.** All means are averages across N = 9 participants.

Supplementary Table S2. Univariate analysis through voxelwise robust regression.

| Structure | Region | Parcel | Network | Volume | MNI Coordinates X | Y | Z | Experienced Pain (Hot-Warm) T | Activated (n) | Deactivated (n) | Imagined Pain (Imagine-Warm) T | Activated (n) | Deactivated (n) | Pain Conjunction Activated (n) | Deactivated (n) |
|---|---|---|---|---|---|---|---|---|---|---|---|---|---|---|---|
| Amygdala L | MTL AMY BL L | AMY BL CeM L | subctx | 880 | -21 | -4 | -20 | -3 | 0 | 6 | 9.01 | 1 | 0 | 0 | 0 |
| Amygdala L | MTL AMY CeM L | AMY BL CeM L | subctx | 520 | -20 | -5 | -16 | 5.02 | 1 | 4 | 5.62 | 2 | 0 | 0 | 0 |
| Amygdala L | MTL AMY La L | AMY La L | subctx | 736 | -28 | -3 | -23 | -3.57 | 0 | 6 | -3.01 | 0 | 1 | 0 | 1 |
| Amygdala R | MTL AMY BL R | AMY BL CeM R | subctx | 816 | 22 | -4 | -19 | -3.91 | 0 | 5 | 10.31 | 0 | 0 | 0 | 0 |
| Amygdala R | MTL AMY CeM R | AMY BL CeM R | subctx | 608 | 20 | -5 | -16 | 4.1 | 1 | 3 | 3.21 | 2 | 0 | 0 | 0 |
| Amygdala R | MTL AMY La R | AMY La R | subctx | 784 | 28 | -2 | -24 | -5.03 | 0 | 5 | -1.96 | 0 | 0 | 0 | 0 |
| CAU L | BG CAU DA L | aCAU L | subctx | 1832 | -14 | 19 | 6 | -4.05 | 0 | 2 | -3.34 | 1 | 1 | 0 | 0 |
| CAU L | BG CAU VA L | aCAU L | subctx | 1144 | -10 | 12 | 5 | 6.72 | 1 | 1 | 3.21 | 1 | 1 | 1 | 0 |
| CAU L | BG CAU body L | pCAU L | subctx | 1168 | -13 | 6 | 17 | 8.06 | 2 | 0 | 1.92 | 2 | 1 | 0 | 0 |
| CAU L | BG CAU tail L | pCAU L | subctx | 1408 | -16 | -9 | 23 | 6.98 | 2 | 0 | 5.86 | 1 | 0 | 1 | 0 |
| CAU R | BG CAU DA R | aCAU R | subctx | 2184 | 14 | 20 | 7 | 6.32 | 1 | 2 | -7.66 | 2 | 1 | 0 | 0 |
| CAU R | BG CAU VA R | aCAU R | subctx | 1008 | 10 | 13 | 4 | 4.65 | 4 | 1 | -5.02 | 1 | 1 | 1 | 0 |
| CAU R | BG CAU body R | pCAU R | subctx | 1248 | 13 | 7 | 17 | 10.04 | 4 | 0 | -5.47 | 3 | 1 | 1 | 0 |
| CAU R | BG CAU tail R | pCAU R | subctx | 1336 | 16 | -9 | 23 | 4.74 | 4 | 0 | 3.29 | 3 | 1 | 2 | 0 |
| Cblm cortex L | Cblm CrusI L | Cblm CrusI CrusII VIIIb L | subctx | 15848 | -36 | -68 | -32 | 8.54 | 5 | 0 | -6.83 | 5 | 1 | 3 | 0 |
| Cblm cortex L | Cblm CrusII L | Cblm CrusI CrusII VIIIb L | subctx | 11888 | -27 | -74 | -42 | -7.5 | 4 | 0 | -5 | 1 | 3 | 0 | 0 |
| Cblm cortex L | Cblm I IV L | Cblm I IV V VI L | subctx | 3480 | -8 | -45 | -16 | -6.94 | 2 | 0 | 3.15 | 3 | 0 | 2 | 0 |
| Cblm cortex L | Cblm IX L | Cblm VIIIa VIIb IX X L | subctx | 4072 | -7 | -55 | -49 | -15.84 | 1 | 2 | -11.38 | 0 | 4 | 0 | 0 |
| Cblm cortex L | Cblm V L | Cblm I IV V VI L | subctx | 5016 | -15 | -50 | -19 | -3.13 | 3 | 0 | 3.63 | 3 | 0 | 2 | 0 |
| Cblm cortex L | Cblm VI L | Cblm I IV V VI L | subctx | 11192 | -24 | -59 | -25 | 11.01 | 5 | 0 | -4.5 | 3 | 1 | 2 | 0 |
| Cblm cortex L | Cblm VIIIa L | Cblm VIIIa VIIb IX X L | subctx | 6384 | -25 | -59 | -52 | 12.39 | 6 | 0 | -6.76 | 2 | 0 | 0 | 0 |
| Cblm cortex L | Cblm VIIIb L | Cblm VIIIa VIIb IX X L | subctx | 5368 | -18 | -51 | -55 | -9.83 | 2 | 0 | -8.19 | 0 | 2 | 0 | 0 |
| Cblm cortex L | Cblm VIIb L | Cblm CrusI CrusII VIIIb L | subctx | 5552 | -26 | -67 | -50 | 6 | 3 | 1 | 3.94 | 2 | 2 | 1 | 0 |
| Cblm cortex L | Cblm X L | Cblm VIIIa VIIb IX X L | subctx | 904 | -21 | -37 | -45 | 1.63 | 1 | 0 | -6.14 | 0 | 2 | 0 | 0 |
| Cblm cortex R | Cblm CrusI R | Cblm CrusI CrusII VIIIb R | subctx | 16136 | 37 | -67 | -33 | -7 | 4 | 1 | 4.5 | 5 | 0 | 3 | 0 |
| Cblm cortex R | Cblm CrusII R | Cblm CrusI CrusII VIIIb R | subctx | 11832 | 26 | -76 | -42 | -5.03 | 1 | 1 | 2.83 | 3 | 1 | 1 | 0 |
| Cblm cortex R | Cblm I IV R | Cblm I IV V VI R | subctx | 3672 | 10 | -44 | -17 | 4.44 | 3 | 0 | 3.08 | 2 | 1 | 1 | 0 |

| Structure | Region | Parcel | Network | Volume | MNI Coordinates X | Y | Z | Experienced Pain (Hot-Warm) T | Activated (n) | Deactivated (n) | Imagined Pain (Imagine-Warm) T | Activated (n) | Deactivated (n) | Pain Conjunction Activated (n) | Deactivated (n) |
|---|---|---|---|---|---|---|---|---|---|---|---|---|---|---|---|
| Cblm cortex R | Cblm IX R | Cblm VIIIa VIIb IX X R | subctx | 4416 | 7 | -55 | -49 | -21.47 | 0 | 3 | -5.83 | 0 | 2 | 0 | 1 |
| Cblm cortex R | Cblm V R | Cblm I IV V VI R | subctx | 5336 | 15 | -51 | -20 | 5.09 | 4 | 0 | 3.78 | 4 | 0 | 2 | 0 |
| Cblm cortex R | Cblm VI R | Cblm I IV V VI R | subctx | 9816 | 24 | -59 | -26 | 6.53 | 5 | 0 | 11.34 | 5 | 0 | 3 | 0 |
| Cblm cortex R | Cblm VIIIa R | Cblm VIIIa VIIb IX X R | subctx | 6464 | 24 | -59 | -53 | 7.66 | 4 | 0 | 6.85 | 3 | 1 | 1 | 0 |
| Cblm cortex R | Cblm VIIIb R | Cblm VIIIa VIIb IX X R | subctx | 5144 | 17 | -52 | -56 | -4.5 | 2 | 0 | -3.42 | 1 | 3 | 0 | 0 |
| Cblm cortex R | Cblm VIIb R | Cblm CrusI CrusII VIIIb R | subctx | 6216 | 27 | -66 | -51 | 6.63 | 4 | 0 | 3.97 | 4 | 0 | 1 | 0 |
| Cblm cortex R | Cblm X R | Cblm VIIIa VIIb IX X R | subctx | 920 | 22 | -37 | -46 | -4.27 | 1 | 0 | -5.19 | 1 | 2 | 0 | 0 |
| Cblm vermis | Cblm Vermis CrusII | Cblm vermis | subctx | 504 | 1 | -75 | -31 | 1.61 | 2 | 1 | 1.72 | 1 | 1 | 0 | 1 |
| Cblm vermis | Cblm Vermis IX | Cblm vermis | subctx | 976 | -0 | -57 | -38 | -2.83 | 2 | 0 | -2.7 | 1 | 0 | 0 | 0 |
| Cblm vermis | Cblm Vermis VI | Cblm vermis | subctx | 2600 | 1 | -71 | -20 | 2.99 | 5 | 0 | 3.9 | 5 | 0 | 2 | 0 |
| Cblm vermis | Cblm Vermis VIIIa | Cblm vermis | subctx | 1464 | 0 | -68 | -37 | -2.05 | 1 | 1 | 1.32 | 1 | 1 | 0 | 1 |
| Cblm vermis | Cblm Vermis VIIIb | Cblm vermis | subctx | 848 | 0 | -65 | -41 | -1.39 | 3 | 1 | -1.12 | 1 | 1 | 0 | 0 |
| Cblm vermis | Cblm Vermis VIIb | Cblm vermis | subctx | 216 | 0 | -70 | -31 | -1.94 | 1 | 1 | 0.97 | 1 | 0 | 0 | 0 |
| Cblm vermis | Cblm Vermis X | Cblm vermis | subctx | 352 | 0 | -49 | -36 | -5.35 | 0 | 0 | -2.24 | 1 | 1 | 0 | 0 |
| GP L | BG GPe L | GPe L | subctx | 1648 | -20 | -4 | -2 | 6.26 | 3 | 0 | 5.56 | 2 | 0 | 2 | 0 |
| GP L | BG GPi L | GPi L | subctx | 864 | -18 | -6 | -4 | 5.23 | 3 | 0 | 4.71 | 2 | 0 | 1 | 0 |
| GP R | BG GPe R | GPe R | subctx | 1200 | 20 | -2 | -1 | 8.54 | 3 | 0 | 2.5 | 2 | 0 | 2 | 0 |
| GP R | BG GPi R | GPi R | subctx | 1056 | 19 | -5 | -4 | 5.56 | 2 | 0 | 4.68 | 1 | 0 | 1 | 0 |
| Hippocampal Formation L | MTL Hipp CA1 L | Hipp L | subctx | 2840 | -29 | -24 | -14 | -11.03 | 0 | 7 | 5.15 | 0 | 0 | 0 | 0 |
| Hippocampal Formation L | MTL Hipp CA23 L | Hipp L | subctx | 1344 | -25 | -20 | -14 | -9.45 | 0 | 6 | 7.04 | 1 | 0 | 0 | 0 |
| Hippocampal Formation L | MTL Hipp DG L | Hipp L | subctx | 872 | -27 | -26 | -12 | -5.11 | 0 | 5 | 2.64 | 0 | 0 | 0 | 0 |
| Hippocampal Formation L | MTL Hipp Subiculum L | Subiculum L | subctx | 1856 | -21 | -19 | -19 | -4.72 | 0 | 5 | 2.72 | 0 | 1 | 0 | 1 |
| Hippocampal Formation R | MTL Hipp CA1 R | Hipp R | subctx | 2960 | 28 | -23 | -14 | -7.06 | 0 | 6 | -6.93 | 1 | 0 | 0 | 0 |
| Hippocampal Formation R | MTL Hipp CA23 R | Hipp R | subctx | 1336 | 25 | -18 | -15 | -4.98 | 0 | 6 | 5.94 | 2 | 0 | 0 | 0 |
| Hippocampal Formation R | MTL Hipp DG R | Hipp R | subctx | 560 | 29 | -24 | -13 | -9.34 | 0 | 7 | 2.05 | 0 | 0 | 0 | 0 |
| Hippocampal Formation R | MTL Hipp Subiculum R | Subiculum R | subctx | 1928 | 22 | -18 | -19 | -6.37 | 0 | 6 | -4.81 | 0 | 1 | 0 | 0 |
| Medulla | BStem RObPaMg | Medullary Raphe (Serotonergic) | subctx | 584 | -0 | -37 | -53 | 3.65 | 4 | 0 | 2.99 | 0 | 0 | 0 | 0 |
| Medulla L | BStem OC L | Olivary complex L | subctx | 832 | -7 | -37 | -53 | 2.85 | 3 | 0 | 4.34 | 0 | 0 | 0 | 0 |
| Medulla L | BStem PCRtA L | Medullary reticular formation L | subctx | 112 | -9 | -39 | -42 | 1.31 | 2 | 1 | -1.94 | 0 | 0 | 0 | 0 |

| | | | | | MNI Coordinates | | | Experienced Pain (Hot-Warm) | | | Imagined Pain (Imagine-Warm) | | | Pain Conjunction | |
|---|---|---|---|---|---|---|---|---|---|---|---|---|---|---|---|
| Structure | Region | Parcel | Network | Volume | X | Y | Z | T | Activated (n) | Deactivated (n) | T | Activated (n) | Deactivated (n) | Activated (n) | Deactivated (n) |
| Medulla L | BStem Shen Med L | Shen Med L | subctx | 1952 | -4 | -41 | -59 | -5.54 | 2 | 2 | 3.33 | 0 | 0 | 0 | 0 |
| Medulla L | BStem VSM L | Cranial nuclei L | subctx | 208 | -3 | -45 | -54 | 2.28 | 2 | 0 | 2.22 | 1 | 0 | 0 | 0 |
| Medulla L | BStem Ve L | Cranial nuclei L | subctx | 456 | -7 | -42 | -39 | -4.99 | 1 | 1 | -3.08 | 0 | 0 | 0 | 0 |
| Medulla L | BStem iMRt L | Medullary reticular formation L | subctx | 632 | -5 | -44 | -59 | 2.29 | 2 | 0 | 3.48 | 0 | 0 | 0 | 0 |
| Medulla L | BStem sMRt L | Medullary reticular formation L | subctx | 152 | -6 | -42 | -47 | -2.33 | 3 | 1 | 1.2 | 0 | 0 | 0 | 0 |
| Medulla R | BStem OC R | Olivary complex R | subctx | 784 | 6 | -36 | -53 | 3.01 | 5 | 0 | 4.18 | 0 | 0 | 0 | 0 |
| Medulla R | BStem PCRtA R | Medullary reticular formation R | subctx | 168 | 9 | -38 | -42 | 2.43 | 2 | 0 | 1.73 | 0 | 0 | 0 | 0 |
| Medulla R | BStem Shen Med R | Shen Med R | subctx | 2104 | 5 | -41 | -60 | -5.16 | 1 | 0 | 8.21 | 0 | 0 | 0 | 0 |
| Medulla R | BStem VSM R | Cranial nuclei R | subctx | 160 | 2 | -44 | -54 | 2.3 | 2 | 0 | 3.15 | 0 | 0 | 0 | 0 |
| Medulla R | BStem Ve R | Cranial nuclei R | subctx | 344 | 6 | -42 | -40 | -4.88 | 0 | 1 | -1.58 | 0 | 1 | 0 | 1 |
| Medulla R | BStem iMRt R | Medullary reticular formation R | subctx | 752 | 4 | -44 | -59 | -3.8 | 2 | 0 | 6.3 | 0 | 0 | 0 | 0 |
| Medulla R | BStem sMRt R | Medullary reticular formation R | subctx | 160 | 6 | -42 | -48 | -3.09 | 4 | 1 | -3.55 | 0 | 1 | 0 | 1 |
| Midbrain | BStem CLi RLi | Rostral Raphe (Serotonergic) | subctx | 264 | -1 | -24 | -9 | 6.83 | 5 | 0 | 1.29 | 0 | 0 | 0 | 0 |
| Midbrain | BStem DR B7 | Rostral Raphe (Serotonergic) | subctx | 112 | -0 | -34 | -19 | 1.86 | 2 | 0 | 1.03 | 0 | 0 | 0 | 0 |
| Midbrain | BStem MnR | Rostral Raphe (Serotonergic) | subctx | 120 | 0 | -31 | -21 | 2.18 | 2 | 0 | -1.71 | 0 | 0 | 0 | 0 |
| Midbrain | BStem PAG | PAG | subctx | 712 | 0 | -32 | -10 | 3.06 | 4 | 0 | 2.25 | 2 | 2 | 0 | 0 |
| Midbrain L | BStem IC L | Tectum L | subctx | 240 | -6 | -38 | -10 | -4.03 | 0 | 0 | 1.92 | 2 | 1 | 0 | 0 |
| Midbrain L | BStem MiTg PBG L | Tectum L | subctx | 96 | -9 | -32 | -10 | 1.72 | 2 | 0 | 1.46 | 0 | 0 | 0 | 0 |
| Midbrain L | BStem RN L | RN L | subctx | 416 | -5 | -19 | -9 | 4.78 | 3 | 0 | 1.18 | 0 | 0 | 0 | 0 |
| Midbrain L | BStem SC L | Tectum L | subctx | 248 | -5 | -33 | -4 | -2.25 | 1 | 0 | -0.78 | 1 | 2 | 0 | 0 |
| Midbrain L | BStem SN L | VTA PBP SN L | subctx | 552 | -9 | -17 | -13 | 4.62 | 3 | 0 | 2.53 | 1 | 0 | 1 | 0 |
| Midbrain L | BStem STH L | STH L | subctx | 224 | -10 | -11 | -7 | 4.37 | 2 | 0 | 2.88 | 1 | 0 | 1 | 0 |
| Midbrain L | BStem Shen Midb Lc | Shen Midb Lc | subctx | 4848 | -8 | -19 | -16 | -5.16 | 5 | 0 | 6.81 | 2 | 0 | 0 | 0 |
| Midbrain L | BStem Shen Midb Lrd | Shen Midb Lrd | subctx | 2264 | -13 | -21 | -6 | 9.15 | 2 | 0 | 9.66 | 3 | 0 | 1 | 0 |
| Midbrain L | BStem VTA PBP L | VTA PBP SN L | subctx | 160 | -7 | -18 | -12 | 3.99 | 3 | 0 | 1.47 | 1 | 0 | 1 | 0 |
| Midbrain L | BStem isRt L | Rostral reticular formation L | subctx | 160 | -6 | -29 | -8 | 2.64 | 1 | 0 | -0.61 | 1 | 2 | 0 | 0 |
| Midbrain L | BStem mRt L | Rostral reticular formation L | subctx | 688 | -8 | -25 | -3 | 5.47 | 5 | 0 | 1.86 | 2 | 1 | 1 | 0 |
| Midbrain R | BStem IC R | Tectum R | subctx | 296 | 5 | -37 | -9 | -4.24 | 0 | 0 | 2.78 | 1 | 2 | 0 | 0 |
| Midbrain R | BStem MiTg PBG R | Tectum R | subctx | 80 | 9 | -31 | -10 | 1.77 | 3 | 0 | -2.71 | 0 | 0 | 0 | 0 |

| | | | | | MNI Coordinates | | | Experienced Pain (Hot-Warm) | | | Imagined Pain (Imagine-Warm) | | | Pain Conjunction | |
|---|---|---|---|---|---|---|---|---|---|---|---|---|---|---|---|
| Structure | Region | Parcel | Network | Volume | X | Y | Z | T | Activated (n) | Deactivated (n) | T | Activated (n) | Deactivated (n) | Activated (n) | Deactivated (n) |
| Midbrain R | BStem RN R | RN R | subctx | 392 | 5 | -19 | -9 | 4.49 | 4 | 0 | -1.96 | 1 | 0 | 1 | 0 |
| Midbrain R | BStem SC R | Tectum R | subctx | 192 | 4 | -33 | -4 | -4.46 | 1 | 0 | -2.18 | 1 | 3 | 0 | 0 |
| Midbrain R | BStem SN R | VTA PBP SN R | subctx | 560 | 10 | -17 | -13 | 4.58 | 5 | 0 | 4.15 | 1 | 0 | 1 | 0 |
| Midbrain R | BStem STH R | STH R | subctx | 232 | 11 | -11 | -7 | 5.44 | 4 | 0 | 4.46 | 1 | 0 | 1 | 0 |
| Midbrain R | BStem Shen Midb Rcd | Shen Midb Rcd | subctx | 5392 | 9 | -21 | -16 | 5.91 | 3 | 0 | 4.74 | 1 | 0 | 0 | 0 |
| Midbrain R | BStem Shen Midb Rrd | Shen Midb Rrd | subctx | 1936 | 13 | -21 | -5 | 5.32 | 4 | 0 | 3.99 | 2 | 0 | 1 | 0 |
| Midbrain R | BStem VTA PBP R | VTA PBP SN R | subctx | 152 | 7 | -18 | -13 | 4.14 | 5 | 0 | 1.19 | 2 | 0 | 2 | 0 |
| Midbrain R | BStem isRt R | Rostral reticular formation R | subctx | 120 | 6 | -28 | -8 | 3.21 | 4 | 0 | -1.6 | 1 | 1 | 0 | 0 |
| Midbrain R | BStem mRt R | Rostral reticular formation R | subctx | 720 | 8 | -24 | -4 | 3.47 | 6 | 0 | -2.09 | 3 | 1 | 1 | 0 |
| PUT L | BG PUT DA L | aPUT L | subctx | 1912 | -25 | 6 | 0 | 19.35 | 6 | 0 | 4.14 | 3 | 1 | 3 | 0 |
| PUT L | BG PUT DP L | pPUT L | subctx | 1952 | -27 | -2 | 8 | 7.96 | 4 | 0 | 5.4 | 3 | 0 | 2 | 0 |
| PUT L | BG PUT VA L | aPUT L | subctx | 2504 | -21 | 10 | -7 | 10.33 | 4 | 2 | 5.84 | 4 | 0 | 3 | 0 |
| PUT L | BG PUT VP L | pPUT L | subctx | 2736 | -31 | -11 | -1 | 8.65 | 3 | 0 | 6.22 | 3 | 0 | 2 | 0 |
| PUT R | BG PUT DA R | aPUT R | subctx | 2048 | 26 | 7 | -0 | 6.86 | 5 | 0 | 3.02 | 4 | 0 | 4 | 0 |
| PUT R | BG PUT DP R | pPUT R | subctx | 1584 | 27 | -2 | 6 | 6.17 | 4 | 0 | 2.85 | 3 | 0 | 3 | 0 |
| PUT R | BG PUT VA R | aPUT R | subctx | 2624 | 20 | 12 | -7 | 8.92 | 4 | 0 | 5.49 | 3 | 0 | 3 | 0 |
| PUT R | BG PUT VP R | pPUT R | subctx | 2368 | 31 | -10 | -2 | 14.22 | 5 | 0 | 6.04 | 3 | 0 | 2 | 0 |
| Pons L | BStem LC+ L | LC+ L | subctx | 176 | -7 | -37 | -35 | 1.49 | 2 | 0 | 3.64 | 0 | 0 | 0 | 0 |
| Pons L | BStem LDTg CGPn L | Cholinergic nuclei L | subctx | 488 | -2 | -40 | -33 | 2.7 | 1 | 1 | 2.86 | 0 | 0 | 0 | 0 |
| Pons L | BStem LPB L | Parabrachial nuclei L | subctx | 288 | -8 | -38 | -21 | 1.45 | 1 | 0 | 3.9 | 1 | 0 | 0 | 0 |
| Pons L | BStem MPB L | Parabrachial nuclei L | subctx | 216 | -6 | -37 | -24 | 1.24 | 2 | 0 | 2.24 | 0 | 0 | 0 | 0 |
| Pons L | BStem PTg L | Cholinergic nuclei L | subctx | 144 | -8 | -30 | -14 | 2.18 | 4 | 0 | 1.23 | 0 | 0 | 0 | 0 |
| Pons L | BStem PnO PnC L | L PnO PnC L | subctx | 680 | -3 | -35 | -33 | -5.88 | 2 | 0 | -1.79 | 0 | 1 | 0 | 0 |
| Pons L | BStem Shen Pons Lcd | Shen Pons Lcd | subctx | 2520 | -9 | -32 | -40 | -8.96 | 1 | 3 | -3.23 | 0 | 1 | 0 | 0 |
| Pons L | BStem Shen Pons Lrd | Shen Pons Lrd | subctx | 1744 | -9 | -36 | -28 | -7.36 | 1 | 0 | 13.53 | 1 | 0 | 0 | 0 |
| Pons L | BStem Shen Pons Lv | Shen Pons Lv | subctx | 4568 | -7 | -21 | -35 | -14.04 | 0 | 3 | 3.13 | 0 | 0 | 0 | 0 |
| Pons R | BStem LC+ R | LC+ R | subctx | 176 | 7 | -38 | -35 | 2.35 | 3 | 0 | -1.59 | 1 | 0 | 1 | 0 |
| Pons R | BStem LDTg CGPn R | Cholinergic nuclei R | subctx | 440 | 2 | -40 | -31 | 1.66 | 2 | 1 | -2.14 | 1 | 1 | 1 | 1 |
| Pons R | BStem LPB R | Parabrachial nuclei R | subctx | 328 | 7 | -38 | -21 | 1.31 | 2 | 0 | 1.87 | 1 | 0 | 0 | 0 |
| Pons R | BStem MPB R | Parabrachial nuclei R | subctx | 216 | 5 | -38 | -24 | 1.39 | 2 | 0 | 2.05 | 1 | 0 | 0 | 0 |
| Pons R | BStem PTg R | Cholinergic nuclei R | subctx | 160 | 7 | -30 | -15 | 3.01 | 4 | 0 | 2.25 | 0 | 0 | 0 | 0 |
| Pons R | BStem PnO PnC R | R PnO PnC R | subctx | 616 | 3 | -35 | -33 | -2.39 | 4 | 0 | -3.99 | 1 | 0 | 1 | 0 |

| Structure | Region | Parcel | Network | Volume | MNI Coordinates X | Y | Z | Experienced Pain (Hot-Warm) T | Activated (n) | Deactivated (n) | Imagined Pain (Imagine-Warm) T | Activated (n) | Deactivated (n) | Pain Conjunction Activated (n) | Deactivated (n) |
|---|---|---|---|---|---|---|---|---|---|---|---|---|---|---|---|
| Pons R | BStem Shen Pons Rcd | Shen Pons Rcd | subctx | 2488 | 9 | -36 | -36 | -11.64 | 2 | 0 | -3.94 | 0 | 0 | 0 | 0 |
| Pons R | BStem Shen Pons Rcv | Shen Pons Rcv | subctx | 1920 | 6 | -24 | -43 | -5.24 | 0 | 2 | -1.98 | 0 | 0 | 0 | 0 |
| Pons R | BStem Shen Pons Rrv | Shen Pons Rrv | subctx | 4224 | 10 | -20 | -30 | -5.74 | 0 | 2 | -4.19 | 0 | 2 | 0 | 0 |
| Thal Anterior L | Thal AV L | Thal anterior L | subctx | 760 | -4 | -4 | 10 | 4.62 | 4 | 0 | 1.9 | 2 | 0 | 2 | 0 |
| Thal Anterior R | Thal AV R | Thal anterior R | subctx | 872 | 4 | -3 | 9 | 5.32 | 4 | 0 | 1.98 | 0 | 0 | 0 | 0 |
| Thal Intralaminar L | Thal CL L | Thal Intralaminar-anterodorsal L | subctx | 160 | -2 | -11 | 13 | 3.65 | 4 | 0 | 1.5 | 2 | 0 | 2 | 0 |
| Thal Intralaminar L | Thal cIL L | Thal Intralaminar-posterior L | subctx | 448 | -10 | -20 | 1 | 3.35 | 6 | 0 | 3.36 | 2 | 0 | 2 | 0 |
| Thal Intralaminar L | Thal rIL L | Thal Intralaminar-antoventral L | subctx | 216 | -3 | -10 | -1 | 3.29 | 5 | 0 | 1.17 | 0 | 0 | 0 | 0 |
| Thal Intralaminar R | Thal CL R | Thal Intralaminar-anterodorsal R | subctx | 112 | 4 | -12 | 13 | 3.5 | 6 | 0 | 0.43 | 1 | 1 | 1 | 0 |
| Thal Intralaminar R | Thal cIL R | Thal Intralaminar-posterior R | subctx | 432 | 10 | -19 | 1 | 6.06 | 6 | 0 | 2.92 | 3 | 0 | 2 | 0 |
| Thal Intralaminar R | Thal rIL R | Thal Intralaminar-antoventral R | subctx | 208 | 4 | -9 | -1 | 3.12 | 3 | 0 | 1.67 | 0 | 0 | 0 | 0 |
| Thal Lateral L | Thal LD L | Thal dorsal L | subctx | 336 | -5 | -15 | 18 | 5.15 | 2 | 0 | 1.12 | 0 | 0 | 0 | 0 |
| Thal Lateral L | Thal LP L | Thal dorsal L | subctx | 392 | -10 | -16 | 18 | 3.79 | 3 | 0 | 1.83 | 1 | 0 | 1 | 0 |
| Thal Lateral R | Thal LD R | Thal dorsal R | subctx | 344 | 5 | -15 | 17 | 4.35 | 4 | 0 | -1.69 | 0 | 1 | 0 | 0 |
| Thal Lateral R | Thal LP R | Thal dorsal R | subctx | 440 | 11 | -15 | 18 | 4.92 | 5 | 0 | 1.45 | 1 | 0 | 0 | 0 |
| Thal Medial L | Thal MD L | Thal Medial L | subctx | 1480 | -4 | -15 | 6 | 3.62 | 4 | 0 | 2.81 | 2 | 0 | 2 | 0 |
| Thal Medial R | Thal MD R | Thal Medial R | subctx | 1480 | 5 | -14 | 6 | 3.72 | 5 | 0 | 1.91 | 2 | 1 | 2 | 0 |
| Thal Posterior L | Thal LGN L | Thal LGN L | subctx | 848 | -24 | -27 | -3 | 4.36 | 1 | 1 | 4.83 | 0 | 3 | 0 | 0 |
| Thal Posterior L | Thal MGN L | Thal MGN L | subctx | 424 | -16 | -29 | -3 | -2.96 | 1 | 1 | 2.96 | 0 | 1 | 0 | 0 |
| Thal Posterior L | Thal PuA L | Thal Pulvinar L | subctx | 376 | -11 | -21 | 11 | 3.06 | 4 | 0 | 2.96 | 1 | 0 | 1 | 0 |
| Thal Posterior L | Thal PuL L | Thal Pulvinar L | subctx | 728 | -19 | -27 | 7 | -4.02 | 2 | 0 | 3.39 | 1 | 0 | 0 | 0 |
| Thal Posterior L | Thal PuM L | Thal Pulvinar L | subctx | 3272 | -11 | -29 | 9 | -6.2 | 3 | 0 | 14.5 | 1 | 0 | 0 | 0 |
| Thal Posterior R | Thal LGN R | Thal LGN R | subctx | 752 | 24 | -26 | -3 | 2.98 | 2 | 0 | 3.67 | 1 | 2 | 0 | 0 |
| Thal Posterior R | Thal MGN R | Thal MGN R | subctx | 408 | 16 | -27 | -2 | -5.22 | 1 | 1 | -2.57 | 1 | 1 | 1 | 0 |
| Thal Posterior R | Thal PuA R | Thal Pulvinar R | subctx | 408 | 12 | -20 | 11 | 3.65 | 5 | 0 | 1.63 | 1 | 0 | 1 | 0 |
| Thal Posterior R | Thal PuL R | Thal Pulvinar R | subctx | 712 | 19 | -25 | 7 | -3.95 | 3 | 1 | 2.57 | 0 | 1 | 0 | 0 |
| Thal Posterior R | Thal PuM R | Thal Pulvinar R | subctx | 3096 | 12 | -28 | 9 | -6.02 | 4 | 0 | -2.68 | 1 | 0 | 0 | 0 |
| Thal Ventral L | Thal VA L | Thal Lateral-rostral L | subctx | 864 | -8 | -4 | 4 | 4.91 | 4 | 0 | 1.69 | 3 | 0 | 3 | 0 |
| Thal Ventral L | Thal VL L | Thal Lateral-rostral L | subctx | 2016 | -13 | -11 | 9 | 6.93 | 5 | 0 | 3.12 | 3 | 0 | 3 | 0 |
| Thal Ventral L | Thal VPL VPM L | Thal Lateral-caudal L | subctx | 1432 | -18 | -20 | 5 | 4.26 | 5 | 0 | 4.16 | 4 | 0 | 3 | 0 |
| Thal Ventral R | Thal VA R | Thal Lateral-rostral R | subctx | 864 | 9 | -3 | 4 | 10.16 | 6 | 0 | 2.94 | 2 | 0 | 2 | 0 |

| Structure | Region | Parcel | Network | Volume | MNI Coordinates X | Y | Z | Experienced Pain (Hot-Warm) T | Activated (n) | Deactivated (n) | Imagined Pain (Imagine-Warm) T | Activated (n) | Deactivated (n) | Pain Conjunction Activated (n) | Deactivated (n) |
|---|---|---|---|---|---|---|---|---|---|---|---|---|---|---|---|
| Thal Ventral R | Thal VL R | Thal Lateral-rostral R | subctx | 2040 | 14 | -10 | 9 | 7.03 | 5 | 0 | 2.52 | 2 | 1 | 2 | 0 |
| Thal Ventral R | Thal VPL VPM R | Thal Lateral-caudal R | subctx | 1368 | 18 | -19 | 5 | 5.3 | 5 | 0 | 2.92 | 3 | 0 | 2 | 0 |
| VStriatum L | BG BST SLEA L | VStriatum L | subctx | 608 | -7 | 4 | -1 | -2.55 | 1 | 3 | -5.23 | 2 | 1 | 1 | 0 |
| VStriatum L | BG NAc L | VStriatum L | subctx | 2576 | -8 | 13 | -7 | -4.28 | 0 | 4 | -9.42 | 0 | 2 | 0 | 2 |
| VStriatum R | BG BST SLEA R | VStriatum R | subctx | 816 | 7 | 5 | -0 | 5.32 | 3 | 2 | -3.97 | 2 | 1 | 2 | 0 |
| VStriatum R | BG NAc R | VStriatum R | subctx | 2552 | 7 | 14 | -7 | -4.49 | 0 | 3 | -6.73 | 0 | 1 | 0 | 1 |
| auditory association cortex L | Ctx A4 L | auditory transition area L | Cortex_SomatomotorB | 4032 | -63 | -23 | 10 | 5.82 | 5 | 3 | -6.89 | 2 | 2 | 1 | 2 |
| auditory association cortex L | Ctx A5 L | auditory transition area L | Cortex_Temporal_Parietal | 3704 | -63 | -13 | -1 | -5.26 | 5 | 3 | -4.46 | 2 | 3 | 2 | 1 |
| auditory association cortex L | Ctx STGa L | auditory transition area L | Cortex_Temporal_Parietal | 3064 | -51 | 13 | -14 | 8.06 | 4 | 1 | 3 | 2 | 3 | 1 | 2 |
| auditory association cortex L | Ctx STSda L | anterior AAC L | Cortex_Default_ModeB | 2448 | -55 | -7 | -9 | -5.64 | 1 | 3 | -3.87 | 0 | 6 | 0 | 4 |
| auditory association cortex L | Ctx STSdp L | posterior AAC L | Cortex_Default_ModeB | 2656 | -52 | -31 | 2 | -5.15 | 0 | 4 | -4.46 | 2 | 3 | 0 | 2 |
| auditory association cortex L | Ctx STSva L | anterior AAC L | Cortex_Default_ModeB | 2384 | -51 | -10 | -15 | -7.53 | 0 | 5 | -3.82 | 1 | 3 | 0 | 2 |
| auditory association cortex L | Ctx STSvp L | posterior AAC L | Cortex_Default_ModeB | 4080 | -54 | -34 | -4 | -3.54 | 0 | 4 | -5.44 | 1 | 4 | 0 | 1 |
| auditory association cortex L | Ctx TA2 L | auditory transition area L | Cortex_Temporal_Parietal | 1544 | -50 | 0 | -5 | 11.99 | 9 | 1 | -1.68 | 2 | 2 | 2 | 0 |
| auditory association cortex R | Ctx A4 R | auditory transition area R | Cortex_SomatomotorB | 3864 | 65 | -14 | 7 | 8.28 | 5 | 3 | -9.08 | 1 | 3 | 1 | 2 |
| auditory association cortex R | Ctx A5 R | auditory transition area R | Cortex_Temporal_Parietal | 4616 | 63 | -12 | -2 | -8.57 | 2 | 4 | -4.95 | 0 | 4 | 2 | 1 |
| auditory association cortex R | Ctx STGa R | auditory transition area R | Cortex_Temporal_Parietal | 2688 | 51 | 16 | -17 | -10.3 | 7 | 1 | -4.66 | 2 | 1 | 2 | 1 |
| auditory association cortex R | Ctx STSda R | anterior AAC R | Cortex_Temporal_Parietal | 3160 | 52 | -2 | -14 | -11.18 | 0 | 5 | -12.89 | 0 | 4 | 0 | 3 |
| auditory association cortex R | Ctx STSdp R | posterior AAC R | Cortex_Temporal_Parietal | 3632 | 49 | -27 | -2 | -4.87 | 2 | 1 | -3.87 | 1 | 3 | 0 | 1 |
| auditory association cortex R | Ctx STSva R | anterior AAC R | Cortex_Temporal_Parietal | 2384 | 54 | -10 | -17 | -6.91 | 0 | 5 | -6.04 | 0 | 6 | 0 | 4 |
| auditory association cortex R | Ctx STSvp R | posterior AAC R | Cortex_Default_ModeB | 3240 | 57 | -30 | -5 | -4.08 | 2 | 2 | -5.43 | 1 | 4 | 0 | 2 |
| auditory association cortex R | Ctx TA2 R | auditory transition area R | Cortex_SomatomotorB | 2040 | 51 | 2 | -5 | 6.74 | 7 | 0 | -9.66 | 3 | 2 | 3 | 0 |
| auditory early L | Ctx A1 L | A1 L | Cortex_SomatomotorB | 1448 | -43 | -24 | 12 | 6.76 | 7 | 1 | -3.35 | 0 | 2 | 0 | 1 |
| auditory early L | Ctx LBelt L | auditory+ L | Cortex_SomatomotorB | 1360 | -44 | -27 | 8 | 3.39 | 2 | 2 | -3.63 | 0 | 2 | 0 | 1 |
| auditory early L | Ctx MBelt L | auditory+ L | Cortex_SomatomotorB | 1872 | -45 | -16 | 4 | 6.44 | 7 | 2 | 4.61 | 0 | 2 | 0 | 0 |
| auditory early L | Ctx PBelt L | auditory+ L | Cortex_SomatomotorB | 2504 | -53 | -26 | 9 | 4.14 | 4 | 3 | -4.05 | 0 | 2 | 0 | 1 |
| auditory early L | Ctx RI L | auditory+ L | Cortex_SomatomotorB | 2488 | -39 | -34 | 20 | -3.47 | 2 | 3 | 8.13 | 0 | 2 | 0 | 2 |
| auditory early R | Ctx A1 R | A1 R | Cortex_SomatomotorB | 888 | 43 | -21 | 11 | 5.89 | 7 | 1 | -2.57 | 0 | 2 | 0 | 1 |
| auditory early R | Ctx LBelt R | auditory+ R | Cortex_SomatomotorB | 1624 | 48 | -24 | 10 | 4.25 | 2 | 2 | -3.07 | 0 | 2 | 0 | 1 |
| auditory early R | Ctx MBelt R | auditory+ R | Cortex_SomatomotorB | 1440 | 46 | -15 | 5 | 13.86 | 8 | 0 | -3.5 | 0 | 3 | 0 | 1 |
| auditory early R | Ctx PBelt R | auditory+ R | Cortex_SomatomotorB | 1856 | 57 | -18 | 8 | 7.69 | 6 | 2 | -2.74 | 0 | 3 | 0 | 1 |

| Structure | Region | Parcel | Network | Volume | MNI Coordinates X | Y | Z | Experienced Pain (Hot-Warm) T | Activated (n) | Deactivated (n) | Imagined Pain (Imagine-Warm) T | Activated (n) | Deactivated (n) | Pain Conjunction Activated (n) | Deactivated (n) |
|---|---|---|---|---|---|---|---|---|---|---|---|---|---|---|---|
| auditory early R | Ctx RI R | auditory+ R | Cortex_SomatomotorB | 2056 | 40 | -30 | 20 | 4.61 | 3 | 3 | -4.19 | 0 | 3 | 0 | 2 |
| cingulate ACC mPFC L | Ctx 25 L | infralimbic L | Cortex_Default_ModeA | 2304 | -5 | 24 | -11 | -10.86 | 0 | 5 | -5 | 0 | 2 | 0 | 2 |
| cingulate ACC mPFC L | Ctx 33pr L | midcingulate L | Cortex_Ventral_AttentionB | 1752 | -4 | 11 | 28 | 7.83 | 4 | 1 | -2.62 | 0 | 3 | 0 | 1 |
| cingulate ACC mPFC L | Ctx 8BM L | dmPFC L | Cortex_Fronto_ParietalB | 5048 | -6 | 31 | 45 | 4.6 | 2 | 2 | 2.63 | 2 | 3 | 1 | 0 |
| cingulate ACC mPFC L | Ctx 9m L | dmPFC L | Cortex_Default_ModeB | 8768 | -6 | 53 | 26 | -5.26 | 0 | 3 | -3.06 | 1 | 4 | 0 | 1 |
| cingulate ACC mPFC L | Ctx a24 L | vmPFC L | Cortex_Default_ModeA | 3192 | -5 | 41 | -2 | -3.89 | 0 | 4 | -3.34 | 0 | 1 | 0 | 1 |
| cingulate ACC mPFC L | Ctx a24pr L | midcingulate L | Cortex_Ventral_AttentionB | 1856 | -5 | 17 | 32 | 9.97 | 6 | 0 | 4.74 | 1 | 1 | 1 | 0 |
| cingulate ACC mPFC L | Ctx a32pr L | midcingulate L | Cortex_Ventral_AttentionB | 2360 | -10 | 29 | 30 | 4.12 | 3 | 1 | -3.12 | 1 | 2 | 1 | 0 |
| cingulate ACC mPFC L | Ctx d32 L | dmPFC L | Cortex_Default_ModeA | 3152 | -8 | 40 | 27 | -6.1 | 0 | 3 | -3.4 | 1 | 5 | 0 | 1 |
| cingulate ACC mPFC L | Ctx p24 L | midcingulate L | Cortex_Default_ModeA | 3352 | -5 | 36 | 15 | 2.61 | 2 | 2 | -3.19 | 0 | 2 | 0 | 1 |
| cingulate ACC mPFC L | Ctx p24pr L | midcingulate L | Cortex_Ventral_AttentionA | 1976 | -4 | -4 | 40 | 7.51 | 6 | 0 | 2.58 | 2 | 1 | 1 | 0 |
| cingulate ACC mPFC L | Ctx p32 L | vmPFC L | Cortex_Default_ModeA | 1664 | -13 | 49 | 1 | -4.63 | 0 | 2 | -4.88 | 1 | 2 | 0 | 1 |
| cingulate ACC mPFC L | Ctx p32pr L | midcingulate L | Cortex_Ventral_AttentionA | 2920 | -9 | 14 | 40 | 8.06 | 6 | 0 | 3.16 | 4 | 1 | 2 | 0 |
| cingulate ACC mPFC L | Ctx s32 L | vmPFC L | Cortex_Default_ModeA | 880 | -8 | 35 | -12 | -3.9 | 0 | 4 | -12.09 | 0 | 1 | 0 | 1 |
| cingulate ACC mPFC R | Ctx 25 R | infralimbic R | Cortex_Limbic | 1904 | 3 | 23 | -10 | -14.6 | 0 | 6 | -3.19 | 0 | 3 | 0 | 3 |
| cingulate ACC mPFC R | Ctx 33pr R | midcingulate R | Cortex_Ventral_AttentionB | 2168 | 5 | 12 | 27 | 5.71 | 4 | 0 | -2.83 | 0 | 2 | 0 | 0 |
| cingulate ACC mPFC R | Ctx 8BM R | dmPFC R | Cortex_Fronto_ParietalB | 6232 | 7 | 27 | 48 | 5.39 | 4 | 0 | -11.8 | 1 | 3 | 1 | 0 |
| cingulate ACC mPFC R | Ctx 9m R | dmPFC R | Cortex_Default_ModeA | 10400 | 8 | 53 | 23 | -8.08 | 0 | 5 | -7.72 | 0 | 5 | 0 | 1 |
| cingulate ACC mPFC R | Ctx a24 R | vmPFC R | Cortex_Default_ModeA | 2936 | 6 | 38 | 0 | -4.29 | 1 | 4 | -3.55 | 1 | 2 | 1 | 1 |
| cingulate ACC mPFC R | Ctx a24pr R | midcingulate R | Cortex_Ventral_AttentionB | 2024 | 5 | 17 | 34 | 4.99 | 7 | 0 | 3.39 | 1 | 2 | 2 | 0 |
| cingulate ACC mPFC R | Ctx a32pr R | midcingulate R | Cortex_Ventral_AttentionB | 1696 | 12 | 26 | 32 | -5.16 | 4 | 0 | 5.07 | 1 | 1 | 1 | 0 |
| cingulate ACC mPFC R | Ctx d32 R | dmPFC R | Cortex_Ventral_AttentionB | 3848 | 10 | 38 | 28 | -3.57 | 2 | 2 | -11.62 | 0 | 4 | 0 | 0 |
| cingulate ACC mPFC R | Ctx p24 R | midcingulate R | Cortex_Ventral_AttentionB | 3048 | 6 | 36 | 17 | -2.6 | 3 | 2 | -2.9 | 0 | 3 | 1 | 1 |
| cingulate ACC mPFC R | Ctx p24pr R | midcingulate R | Cortex_Ventral_AttentionA | 2584 | 6 | -4 | 41 | 6.24 | 6 | 0 | 3.28 | 1 | 2 | 1 | 0 |
| cingulate ACC mPFC R | Ctx p32 R | vmPFC R | Cortex_Default_ModeA | 2040 | 11 | 49 | -1 | -4.3 | 1 | 4 | -4.45 | 0 | 2 | 0 | 0 |
| cingulate ACC mPFC R | Ctx p32pr R | midcingulate R | Cortex_Ventral_AttentionB | 2632 | 12 | 13 | 41 | 4.68 | 6 | 0 | 2.95 | 3 | 0 | 2 | 0 |
| cingulate ACC mPFC R | Ctx s32 R | vmPFC R | Cortex_Default_ModeA | 1296 | 6 | 36 | -13 | -7.48 | 0 | 5 | -5.13 | 0 | 3 | 0 | 2 |

| Structure | Region | Parcel | Network | Volume | MNI Coordinates X | Y | Z | Experienced Pain (Hot-Warm) T | Activated (n) | Deactivated (n) | Imagined Pain (Imagine-Warm) T | Activated (n) | Deactivated (n) | Pain Conjunction Activated (n) | Deactivated (n) |
|---|---|---|---|---|---|---|---|---|---|---|---|---|---|---|---|
| cingulate dlPFC L | Ctx 46 L | BA46 L | Cortex_Ventral_AttentionB | 7480 | -37 | 37 | 32 | 10.43 | 6 | 0 | 4.69 | 4 | 2 | 5 | 0 |
| cingulate dlPFC L | Ctx 8Ad L | Superior lateral BA8 L | Cortex_Default_ModeA | 6368 | -24 | 26 | 43 | -12.18 | 0 | 6 | -6.17 | 0 | 4 | 0 | 2 |
| cingulate dlPFC L | Ctx 8Av L | Inferior lateral BA8 L | Cortex_Default_ModeB | 5984 | -38 | 17 | 52 | -4.31 | 0 | 4 | -3.63 | 2 | 1 | 0 | 0 |
| cingulate dlPFC L | Ctx 8BL L | Superior lateral BA8 L | Cortex_Default_ModeB | 4520 | -12 | 35 | 55 | -5.82 | 0 | 4 | -2.96 | 1 | 1 | 0 | 0 |
| cingulate dlPFC L | Ctx 8C L | Inferior lateral BA8 L | Cortex_Fronto_ParietalA | 7160 | -41 | 14 | 38 | -5 | 1 | 3 | 5.12 | 1 | 1 | 1 | 0 |
| cingulate dlPFC L | Ctx 9 46d L | BA46 L | Cortex_Ventral_AttentionB | 6784 | -29 | 45 | 23 | 5.04 | 4 | 1 | 6.43 | 5 | 0 | 3 | 0 |
| cingulate dlPFC L | Ctx 9a L | BA9 L | Cortex_Default_ModeB | 5032 | -20 | 55 | 27 | -6 | 1 | 5 | -2.27 | 4 | 1 | 2 | 2 |
| cingulate dlPFC L | Ctx 9p L | BA9 L | Cortex_Default_ModeB | 4520 | -20 | 45 | 40 | -8.44 | 4 | 2 | -3 | 2 | 2 | 2 | 1 |
| cingulate dlPFC L | Ctx SFL L | Superior lateral BA8 L | Cortex_Default_ModeB | 5616 | -8 | 16 | 65 | -7.79 | 2 | 2 | -3.05 | 4 | 2 | 3 | 0 |
| cingulate dlPFC L | Ctx a9 46v L | BA46 L | Cortex_Ventral_AttentionB | 4632 | -38 | 50 | 10 | 3.53 | 4 | 2 | 5.47 | 3 | 1 | 3 | 0 |
| cingulate dlPFC L | Ctx i6 8 L | Inferior lateral BA8 L | Cortex_Dorsal_AttentionB | 2496 | -32 | 5 | 60 | -3.66 | 0 | 3 | 2.38 | 3 | 0 | 0 | 0 |
| cingulate dlPFC L | Ctx p9 46v L | BA46 L | Cortex_Fronto_ParietalA | 4432 | -44 | 28 | 28 | 4.08 | 3 | 3 | 3.66 | 5 | 0 | 3 | 1 |
| cingulate dlPFC L | Ctx s6 8 L | Superior lateral BA8 L | Cortex_Default_ModeA | 2088 | -23 | 22 | 58 | -4.16 | 0 | 4 | -2.37 | 2 | 1 | 0 | 1 |
| cingulate dlPFC R | Ctx 46 R | BA46 R | Cortex_Ventral_AttentionB | 8176 | 36 | 38 | 31 | 9.74 | 6 | 0 | -5.08 | 1 | 3 | 2 | 0 |
| cingulate dlPFC R | Ctx 8Ad R | Superior lateral BA8 R | Cortex_Default_ModeA | 5296 | 24 | 27 | 45 | -7.8 | 0 | 5 | -13.61 | 0 | 4 | 0 | 3 |
| cingulate dlPFC R | Ctx 8Av R | Inferior lateral BA8 R | Cortex_Fronto_ParietalB | 7128 | 40 | 19 | 49 | 4.65 | 1 | 3 | -8.59 | 0 | 4 | 1 | 1 |
| cingulate dlPFC R | Ctx 8BL R | Superior lateral BA8 R | Cortex_Default_ModeB | 7312 | 13 | 38 | 53 | -6.31 | 1 | 4 | -10.95 | 0 | 5 | 0 | 2 |
| cingulate dlPFC R | Ctx 8C R | Inferior lateral BA8 R | Cortex_Fronto_ParietalA | 5992 | 38 | 16 | 36 | -5.29 | 2 | 3 | -9.34 | 0 | 2 | 0 | 1 |
| cingulate dlPFC R | Ctx 9 46d R | BA46 R | Cortex_Ventral_AttentionB | 7896 | 29 | 48 | 24 | 8.95 | 6 | 0 | -4.32 | 3 | 2 | 2 | 0 |
| cingulate dlPFC R | Ctx 9a R | BA9 R | Cortex_Default_ModeA | 4088 | 20 | 61 | 22 | -6.94 | 0 | 3 | -4.58 | 1 | 3 | 0 | 0 |
| cingulate dlPFC R | Ctx 9p R | BA9 R | Cortex_Default_ModeB | 3936 | 20 | 48 | 36 | -5.14 | 4 | 1 | -3.07 | 3 | 2 | 3 | 0 |
| cingulate dlPFC R | Ctx SFL R | Superior lateral BA8 R | Cortex_Default_ModeB | 3792 | 8 | 13 | 67 | 2.92 | 3 | 3 | -4.13 | 5 | 1 | 3 | 0 |
| cingulate dlPFC R | Ctx a9 46v R | BA46 R | Cortex_Ventral_AttentionB | 3664 | 40 | 52 | 10 | 5.15 | 6 | 2 | -7.57 | 2 | 2 | 2 | 1 |
| cingulate dlPFC R | Ctx i6 8 R | Inferior lateral BA8 R | Cortex_Fronto_ParietalB | 2384 | 35 | 9 | 60 | -4.27 | 2 | 2 | -4.66 | 0 | 3 | 0 | 1 |
| cingulate dlPFC R | Ctx p9 46v R | BA46 R | Cortex_Fronto_ParietalA | 5560 | 46 | 31 | 29 | 8.61 | 3 | 1 | -11.59 | 0 | 4 | 0 | 0 |
| cingulate dlPFC R | Ctx s6 8 R | Superior lateral BA8 R | Cortex_Default_ModeA | 3024 | 22 | 18 | 60 | -3.43 | 0 | 4 | -2.91 | 0 | 4 | 0 | 1 |
| cingulate posterior L | Ctx 23c L | anterior precuneus L | Cortex_Ventral_AttentionA | 2856 | -12 | -31 | 42 | -7.47 | 3 | 3 | -5.4 | 1 | 3 | 0 | 2 |
| cingulate posterior L | Ctx 23d L | anterior precuneus L | Cortex_Default_ModeA | 1728 | -3 | -22 | 39 | 5.1 | 2 | 2 | -9.92 | 0 | 3 | 0 | 1 |
| cingulate posterior L | Ctx 31a L | anterior precuneus L | Cortex_Default_ModeA | 1912 | -4 | -38 | 44 | -7.57 | 1 | 4 | -6.04 | 1 | 4 | 0 | 2 |
| cingulate posterior L | Ctx 31pd L | precuneus L | Cortex_Default_ModeA | 2144 | -10 | -52 | 36 | -15.2 | 0 | 5 | -6.02 | 0 | 3 | 0 | 1 |
| cingulate posterior L | Ctx 31pv L | precuneus L | Cortex_Default_ModeA | 1328 | -11 | -46 | 32 | -12.21 | 0 | 4 | -4.26 | 0 | 3 | 0 | 1 |
| cingulate posterior L | Ctx 7m L | precuneus L | Cortex_Default_ModeA | 2952 | -4 | -62 | 37 | -19.48 | 1 | 4 | -3.91 | 0 | 4 | 0 | 2 |
| cingulate posterior L | Ctx DVT L | parietal occipital sulcus L | Cortex_Visual_Peripheral | 2944 | -20 | -71 | 32 | -8.19 | 0 | 4 | -5.4 | 0 | 2 | 0 | 1 |

| | | | | | MNI Coordinates | | | Experienced Pain (Hot-Warm) | | | Imagined Pain (Imagine-Warm) | | | Pain Conjunction | |
|---|---|---|---|---|---|---|---|---|---|---|---|---|---|---|---|
| Structure | Region | Parcel | Network | Volume | X | Y | Z | T | Activated (n) | Deactivated (n) | T | Activated (n) | Deactivated (n) | Activated (n) | Deactivated (n) |
| cingulate posterior L | Ctx PCV L | PCV L | Cortex_Default_ModeA | 3280 | -6 | -51 | 51 | -12.21 | 1 | 3 | -6.75 | 1 | 3 | 0 | 2 |
| cingulate posterior L | Ctx POS1 L | parietal occipital sulcus L | Cortex_Default_ModeC | 3504 | -13 | -57 | 16 | -11.73 | 1 | 5 | -5.84 | 0 | 2 | 0 | 1 |
| cingulate posterior L | Ctx POS2 L | parietal occipital sulcus L | Cortex_Fronto_ParietalC | 5736 | -10 | -71 | 40 | 11.11 | 5 | 0 | -6.73 | 2 | 1 | 1 | 1 |
| cingulate posterior L | Ctx ProS L | parietal occipital sulcus L | Cortex_Visual_Peripheral | 2944 | -24 | -55 | 7 | -4.07 | 1 | 3 | 3.88 | 2 | 2 | 0 | 1 |
| cingulate posterior L | Ctx RSC L | RSC L | Cortex_Default_ModeC | 5144 | -5 | -38 | 18 | -14.66 | 2 | 2 | -7.98 | 1 | 2 | 0 | 1 |
| cingulate posterior L | Ctx d23ab L | precuneus L | Cortex_Default_ModeA | 1960 | -4 | -43 | 31 | -5.24 | 1 | 4 | -5.02 | 0 | 4 | 0 | 1 |
| cingulate posterior L | Ctx v23ab L | precuneus L | Cortex_Default_ModeA | 1848 | -4 | -58 | 18 | -6.18 | 0 | 4 | -3.81 | 0 | 3 | 0 | 2 |
| cingulate posterior R | Ctx 23c R | anterior precuneus R | Cortex_Ventral_AttentionA | 3184 | 12 | -33 | 44 | -5.07 | 2 | 3 | -3.96 | 1 | 3 | 0 | 1 |
| cingulate posterior R | Ctx 23d R | anterior precuneus R | Cortex_Default_ModeA | 2064 | 4 | -23 | 40 | 3.13 | 3 | 1 | -3.55 | 0 | 3 | 0 | 1 |
| cingulate posterior R | Ctx 31a R | anterior precuneus R | Cortex_Default_ModeA | 1760 | 7 | -40 | 43 | -14.54 | 0 | 4 | -5.16 | 1 | 4 | 0 | 2 |
| cingulate posterior R | Ctx 31pd R | precuneus R | Cortex_Default_ModeA | 1416 | 12 | -52 | 37 | -10.59 | 1 | 3 | -3.61 | 0 | 4 | 0 | 1 |
| cingulate posterior R | Ctx 31pv R | precuneus R | Cortex_Default_ModeA | 1448 | 12 | -44 | 33 | -4.33 | 1 | 3 | -3.29 | 0 | 4 | 0 | 1 |
| cingulate posterior R | Ctx 7m R | precuneus R | Cortex_Default_ModeA | 3496 | 5 | -62 | 37 | -13.69 | 1 | 5 | -4.18 | 0 | 4 | 0 | 3 |
| cingulate posterior R | Ctx DVT R | parietal occipital sulcus R | Cortex_Fronto_ParietalC | 3312 | 23 | -66 | 31 | -3.06 | 1 | 3 | -4.69 | 0 | 4 | 0 | 1 |
| cingulate posterior R | Ctx PCV R | PCV R | Cortex_Fronto_ParietalC | 3592 | 7 | -52 | 52 | -10.32 | 0 | 3 | -4.83 | 1 | 3 | 0 | 2 |
| cingulate posterior R | Ctx POS1 R | parietal occipital sulcus R | Cortex_Default_ModeC | 3392 | 15 | -55 | 18 | -9.27 | 1 | 4 | -11.57 | 0 | 4 | 0 | 2 |
| cingulate posterior R | Ctx POS2 R | parietal occipital sulcus R | Cortex_Fronto_ParietalC | 6144 | 13 | -68 | 40 | 4.25 | 4 | 2 | -4.88 | 0 | 3 | 0 | 1 |
| cingulate posterior R | Ctx ProS R | parietal occipital sulcus R | Cortex_Visual_Peripheral | 2120 | 25 | -48 | 6 | -7.37 | 1 | 2 | 4.98 | 2 | 2 | 0 | 1 |
| cingulate posterior R | Ctx RSC R | RSC R | Cortex_Default_ModeA | 5368 | 6 | -37 | 19 | 15.72 | 3 | 2 | -10.67 | 0 | 5 | 0 | 1 |
| cingulate posterior R | Ctx d23ab R | precuneus R | Cortex_Default_ModeA | 1424 | 5 | -41 | 33 | 10.7 | 1 | 3 | -3.69 | 0 | 4 | 0 | 1 |
| cingulate posterior R | Ctx v23ab R | precuneus R | Cortex_Default_ModeA | 1880 | 5 | -54 | 20 | -6.45 | 0 | 4 | -4.27 | 0 | 4 | 0 | 2 |
| cingulate ventral frontal L | Ctx 10d L | BA10 L | Cortex_Default_ModeA | 6120 | -11 | 64 | 10 | -7.05 | 1 | 4 | -5.31 | 0 | 2 | 0 | 1 |
| cingulate ventral frontal L | Ctx 10pp L | BA10 L | Cortex_Limbic | 2704 | -15 | 61 | -13 | -3.69 | 2 | 2 | 3.9 | 1 | 0 | 0 | 0 |
| cingulate ventral frontal L | Ctx 10r L | BA10 L | Cortex_Default_ModeA | 2568 | -7 | 52 | -7 | -4.9 | 0 | 6 | -3.06 | 0 | 2 | 0 | 1 |
| cingulate ventral frontal L | Ctx 10v L | BA10 L | Cortex_Limbic | 6248 | -5 | 52 | -16 | -6.99 | 0 | 7 | -7.82 | 0 | 1 | 0 | 1 |
| cingulate ventral frontal L | Ctx 11l L | BA11 L | Cortex_Ventral_AttentionB | 4712 | -27 | 47 | -13 | 7.6 | 4 | 2 | 3.2 | 2 | 0 | 1 | 0 |
| cingulate ventral frontal L | Ctx 13l L | BA13 L | Cortex_Limbic | 3648 | -23 | 30 | -18 | 3.5 | 1 | 3 | 5.9 | 4 | 0 | 0 | 0 |
| cingulate ventral frontal L | Ctx 47l L | BA47 L | Cortex_Default_ModeB | 3600 | -46 | 30 | -10 | -3.74 | 2 | 3 | 3.17 | 2 | 1 | 1 | 1 |
| cingulate ventral frontal L | Ctx 47m L | BA47 L | Cortex_Default_ModeB | 912 | -36 | 33 | -14 | -2.13 | 0 | 2 | 1.95 | 2 | 0 | 0 | 0 |
| cingulate ventral frontal L | Ctx 47s L | BA47 L | Cortex_Default_ModeB | 4056 | -33 | 22 | -20 | -3.62 | 1 | 3 | 3.65 | 1 | 1 | 0 | 0 |
| cingulate ventral frontal L | Ctx OFC L | OFC L | Cortex_Limbic | 6640 | -12 | 31 | -21 | -6.44 | 0 | 3 | -8.45 | 0 | 1 | 0 | 1 |

| Structure | Region | Parcel | Network | Volume | MNI Coordinates | | | Experienced Pain (Hot-Warm) | | | Imagined Pain (Imagine-Warm) | | | Pain Conjunction | |
|---|---|---|---|---|---|---|---|---|---|---|---|---|---|---|---|
| | | | | | X | Y | Z | T | Activated (n) | Deactivated (n) | T | Activated (n) | Deactivated (n) | Activated (n) | Deactivated (n) |
| cingulate ventral frontal L | Ctx a10p L | BA10 L | Cortex_Fronto_ParietalB | 4288 | -26 | 52 | -3 | -4.38 | 1 | 1 | -4.3 | 1 | 2 | 0 | 0 |
| cingulate ventral frontal L | Ctx a47r L | BA47 L | Cortex_Fronto_ParietalB | 5272 | -39 | 46 | -9 | -3.31 | 1 | 3 | -2.37 | 3 | 2 | 0 | 1 |
| cingulate ventral frontal L | Ctx p10p L | BA10 L | Cortex_Fronto_ParietalB | 3544 | -24 | 60 | 6 | 5.39 | 0 | 4 | -5.8 | 2 | 2 | 0 | 1 |
| cingulate ventral frontal L | Ctx p47r L | BA47 L | Cortex_Fronto_ParietalB | 3480 | -43 | 42 | 2 | 6.2 | 2 | 1 | 2.21 | 3 | 0 | 1 | 0 |
| cingulate ventral frontal L | Ctx pOFC L | OFC L | Cortex_Limbic | 4008 | -13 | 12 | -20 | -3.74 | 0 | 3 | -4.4 | 1 | 1 | 0 | 1 |
| cingulate ventral frontal R | Ctx 10d R | BA10 R | Cortex_Default_ModeA | 4520 | 10 | 66 | 6 | -11.75 | 0 | 6 | -3.79 | 0 | 4 | 0 | 3 |
| cingulate ventral frontal R | Ctx 10pp R | BA10 R | Cortex_Limbic | 3688 | 14 | 61 | -15 | -7.27 | 1 | 4 | -5.91 | 1 | 2 | 0 | 1 |
| cingulate ventral frontal R | Ctx 10r R | BA10 R | Cortex_Default_ModeA | 2280 | 8 | 51 | -8 | -3.3 | 1 | 3 | -3.39 | 0 | 2 | 0 | 1 |
| cingulate ventral frontal R | Ctx 10v R | BA10 R | Cortex_Default_ModeA | 4208 | 5 | 53 | -15 | -7.59 | 1 | 6 | -5.53 | 0 | 2 | 0 | 1 |
| cingulate ventral frontal R | Ctx 11l R | BA11 R | Cortex_Fronto_ParietalB | 5240 | 26 | 47 | -14 | 7.71 | 3 | 1 | -5.43 | 2 | 1 | 1 | 0 |
| cingulate ventral frontal R | Ctx 13l R | BA13 R | Cortex_Limbic | 1784 | 22 | 29 | -16 | -2.69 | 0 | 2 | -3.93 | 2 | 0 | 0 | 0 |
| cingulate ventral frontal R | Ctx 47l R | BA47 R | Cortex_Default_ModeB | 3344 | 46 | 32 | -13 | -4.03 | 3 | 2 | 1.51 | 3 | 2 | 1 | 0 |
| cingulate ventral frontal R | Ctx 47m R | BA47 R | Cortex_Default_ModeB | 1416 | 34 | 34 | -15 | 4.14 | 2 | 2 | 3.81 | 2 | 0 | 0 | 0 |
| cingulate ventral frontal R | Ctx 47s R | BA47 R | Cortex_Default_ModeB | 4136 | 31 | 23 | -20 | -3.78 | 4 | 1 | -4.64 | 2 | 1 | 1 | 0 |
| cingulate ventral frontal R | Ctx OFC R | OFC R | Cortex_Limbic | 7544 | 10 | 34 | -22 | -9.69 | 0 | 4 | -7.31 | 0 | 1 | 0 | 1 |
| cingulate ventral frontal R | Ctx a10p R | BA10 R | Cortex_Fronto_ParietalB | 2600 | 26 | 60 | -7 | 4.4 | 3 | 0 | -8.25 | 1 | 3 | 1 | 0 |
| cingulate ventral frontal R | Ctx a47r R | BA47 R | Cortex_Fronto_ParietalB | 5712 | 39 | 52 | -7 | 2.93 | 4 | 1 | -10.93 | 1 | 4 | 1 | 0 |
| cingulate ventral frontal R | Ctx p10p R | BA10 R | Cortex_Fronto_ParietalB | 3760 | 26 | 59 | 6 | -3.74 | 3 | 1 | -5.02 | 0 | 3 | 0 | 0 |
| cingulate ventral frontal R | Ctx p47r R | BA47 R | Cortex_Ventral_AttentionB | 2960 | 45 | 44 | -3 | 10.57 | 4 | 1 | -2.3 | 3 | 3 | 1 | 0 |
| cingulate ventral frontal R | Ctx pOFC R | OFC R | Cortex_Limbic | 4704 | 14 | 14 | -17 | -5.16 | 0 | 3 | 2.45 | 1 | 0 | 0 | 0 |
| cingulate vlPFC L | Ctx 44 L | BA44 L | Cortex_Default_ModeB | 3440 | -52 | 15 | 14 | 11.88 | 4 | 1 | 7.43 | 3 | 1 | 2 | 0 |
| cingulate vlPFC L | Ctx 45 L | BA45 L | Cortex_Default_ModeB | 4952 | -50 | 26 | 3 | 4.06 | 3 | 1 | -3.44 | 1 | 1 | 1 | 0 |
| cingulate vlPFC L | Ctx IFJa L | IFJ L | Cortex_Fronto_ParietalA | 2704 | -39 | 13 | 26 | 2.67 | 2 | 2 | 6.6 | 3 | 0 | 2 | 0 |
| cingulate vlPFC L | Ctx IFJp L | IFJ L | Cortex_Fronto_ParietalA | 2088 | -38 | 3 | 31 | 4.56 | 1 | 3 | 4.62 | 2 | 2 | 0 | 2 |
| cingulate vlPFC L | Ctx IFSa L | IFS L | Cortex_Fronto_ParietalA | 3376 | -47 | 33 | 12 | 6.07 | 4 | 2 | 4.01 | 5 | 0 | 2 | 0 |
| cingulate vlPFC L | Ctx IFSp L | IFS L | Cortex_Fronto_ParietalA | 2160 | -49 | 22 | 23 | 4.03 | 1 | 2 | 2.41 | 4 | 1 | 0 | 0 |
| cingulate vlPFC R | Ctx 44 R | BA44 R | Cortex_Default_ModeB | 4208 | 54 | 19 | 11 | 6.86 | 6 | 0 | -4.54 | 3 | 2 | 2 | 0 |
| cingulate vlPFC R | Ctx 45 R | BA45 R | Cortex_Default_ModeB | 3376 | 51 | 28 | 2 | 4.76 | 5 | 0 | -6.25 | 2 | 3 | 1 | 1 |
| cingulate vlPFC R | Ctx IFJa R | IFJ R | Cortex_Fronto_ParietalA | 2056 | 43 | 18 | 24 | 10.6 | 3 | 1 | -3.2 | 1 | 3 | 1 | 1 |
| cingulate vlPFC R | Ctx IFJp R | IFJ R | Cortex_Fronto_ParietalA | 1592 | 38 | 7 | 27 | 6.53 | 2 | 2 | -2.07 | 1 | 2 | 1 | 1 |
| cingulate vlPFC R | Ctx IFSa R | IFS R | Cortex_Ventral_AttentionB | 4080 | 48 | 38 | 6 | 8.42 | 5 | 1 | -2.77 | 3 | 2 | 3 | 1 |

| Structure | Region | Parcel | Network | Volume | MNI Coordinates | | | Experienced Pain (Hot-Warm) | | | Imagined Pain (Imagine-Warm) | | | Pain Conjunction | |
|---|---|---|---|---|---|---|---|---|---|---|---|---|---|---|---|
| | | | | | X | Y | Z | T | Activated (n) | Deactivated (n) | T | Activated (n) | Deactivated (n) | Activated (n) | Deactivated (n) |
| cingulate vlPFC R | Ctx IFSp R | IFS R | Cortex_Fronto_ParietalA | 2672 | 49 | 28 | 19 | 6.07 | 3 | 1 | 7.26 | 0 | 3 | 1 | 0 |
| hypothalamus L | hypothalamus anterior and tubular inferior L | hypothalamus L | subctx | 624 | -4 | -0 | -16 | -1.65 | 0 | 3 | -1.43 | 0 | 1 | 0 | 1 |
| hypothalamus L | hypothalamus anterior and tubular superior L | hypothalamus L | subctx | 576 | -3 | -1 | -9 | -2.61 | 0 | 5 | -1.21 | 0 | 1 | 0 | 1 |
| hypothalamus L | hypothalamus posterior L | hypothalamus L | subctx | 248 | -2 | -8 | -14 | 2.09 | 1 | 1 | 0.91 | 0 | 0 | 0 | 0 |
| hypothalamus R | hypothalamus anterior and tubular inferior R | hypothalamus R | subctx | 632 | 5 | -0 | -16 | -2.01 | 0 | 3 | -1.27 | 0 | 1 | 0 | 1 |
| hypothalamus R | hypothalamus anterior and tubular superior R | hypothalamus R | subctx | 536 | 4 | -1 | -9 | -2.87 | 0 | 4 | 1.67 | 0 | 0 | 0 | 0 |
| hypothalamus R | hypothalamus posterior R | hypothalamus R | subctx | 232 | 3 | -8 | -15 | -1.32 | 1 | 1 | 2.58 | 0 | 0 | 0 | 0 |
| insula anterior L | Ctx AAIC L | anterior agranular insula L | Cortex_Ventral_AttentionB | 1824 | -33 | 14 | -11 | 8.53 | 3 | 0 | 3.37 | 0 | 3 | 0 | 0 |
| insula anterior L | Ctx AVI L | anterior agranular insula L | Cortex_Ventral_AttentionB | 2432 | -29 | 25 | -2 | 6.37 | 5 | 0 | 4.34 | 2 | 1 | 1 | 0 |
| insula anterior L | Ctx MI L | anterior agranular insula L | Cortex_Ventral_AttentionA | 3072 | -35 | 12 | 2 | 10.14 | 9 | 1 | 3.01 | 4 | 1 | 3 | 1 |
| insula anterior L | Ctx Pir L | PIR L | Cortex_Limbic | 2528 | -31 | 4 | -17 | 12 | 3 | 2 | 14.83 | 0 | 1 | 0 | 0 |
| insula anterior R | Ctx AAIC R | anterior agranular insula R | Cortex_Ventral_AttentionA | 2192 | 34 | 17 | -11 | 4.9 | 6 | 1 | -3.21 | 2 | 1 | 2 | 1 |
| insula anterior R | Ctx AVI R | anterior agranular insula R | Cortex_Ventral_AttentionB | 2752 | 31 | 26 | -3 | 17.18 | 7 | 0 | 3.42 | 3 | 2 | 2 | 0 |
| insula anterior R | Ctx MI R | anterior agranular insula R | Cortex_Ventral_AttentionA | 2128 | 36 | 12 | 2 | 17.98 | 9 | 1 | 2.33 | 5 | 1 | 5 | 1 |
| insula anterior R | Ctx Pir R | PIR R | Cortex_Ventral_AttentionA | 3864 | 31 | 6 | -18 | 11.46 | 5 | 2 | -2.74 | 1 | 2 | 0 | 0 |
| insula operculum L | Ctx FOP2 L | posterior operculum L | Cortex_SomatomotorB | 1064 | -42 | -4 | 15 | 9.26 | 8 | 0 | 4.94 | 1 | 2 | 1 | 0 |
| insula operculum L | Ctx FOP3 L | posterior operculum L | Cortex_Ventral_AttentionA | 1296 | -34 | 3 | 14 | 15.36 | 9 | 1 | 3.8 | 2 | 1 | 2 | 1 |
| insula operculum L | Ctx FOP4 L | anterior operculum L | Cortex_Ventral_AttentionA | 3752 | -41 | 13 | 7 | 12.87 | 8 | 0 | 3.64 | 3 | 0 | 2 | 0 |
| insula operculum L | Ctx FOP5 L | anterior operculum L | Cortex_Ventral_AttentionA | 2376 | -35 | 28 | 6 | 6.03 | 7 | 0 | 1.72 | 3 | 2 | 2 | 0 |
| insula operculum R | Ctx FOP2 R | posterior operculum R | Cortex_SomatomotorB | 1056 | 40 | -1 | 16 | 8.74 | 6 | 0 | 2.38 | 0 | 2 | 0 | 0 |
| insula operculum R | Ctx FOP3 R | posterior operculum R | Cortex_Ventral_AttentionA | 872 | 33 | 8 | 12 | 10.97 | 9 | 1 | 2.35 | 4 | 0 | 4 | 1 |
| insula operculum R | Ctx FOP4 R | anterior operculum R | Cortex_Ventral_AttentionA | 2296 | 38 | 15 | 10 | 15.13 | 9 | 1 | 3.2 | 3 | 0 | 3 | 1 |

| Structure | Region | Parcel | Network | Volume | MNI Coordinates X | Y | Z | Experienced Pain (Hot-Warm) T | Activated (n) | Deactivated (n) | Imagined Pain (Imagine-Warm) T | Activated (n) | Deactivated (n) | Pain Conjunction Activated (n) | Deactivated (n) |
|---|---|---|---|---|---|---|---|---|---|---|---|---|---|---|---|
| insula operculum R | Ctx FOP5 R | anterior operculum R | Cortex_Ventral_AttentionB | 2696 | 37 | 29 | 6 | 13.97 | 7 | 0 | 2.53 | 2 | 3 | 1 | 0 |
| insula posterior L | Ctx 52 L | posterior granular insula L | Cortex_SomatomotorB | 1152 | -38 | -21 | 0 | 15.85 | 7 | 1 | 9.45 | 1 | 2 | 1 | 0 |
| insula posterior L | Ctx Ig L | posterior granular insula L | Cortex_SomatomotorB | 1616 | -35 | -14 | 15 | 6.73 | 8 | 0 | 5.15 | 1 | 2 | 1 | 0 |
| insula posterior L | Ctx PI L | inferior dysgranular insula L | Cortex_Limbic | 1400 | -43 | -4 | -13 | 10.27 | 8 | 1 | 5.82 | 1 | 2 | 1 | 1 |
| insula posterior L | Ctx PoI1 L | inferior dysgranular insula L | Cortex_Ventral_AttentionA | 1960 | -38 | -10 | -4 | 14.6 | 9 | 1 | 2.07 | 2 | 1 | 2 | 0 |
| insula posterior L | Ctx PoI2 L | inferior dysgranular insula L | Cortex_Ventral_AttentionA | 3296 | -40 | -1 | -1 | 11.75 | 9 | 1 | 3.01 | 3 | 2 | 2 | 0 |
| insula posterior R | Ctx 52 R | posterior granular insula R | Cortex_SomatomotorB | 800 | 39 | -19 | 1 | 6.18 | 7 | 1 | 4.07 | 0 | 2 | 0 | 0 |
| insula posterior R | Ctx Ig R | posterior granular insula R | Cortex_SomatomotorB | 1432 | 36 | -11 | 13 | 12.57 | 8 | 0 | 4.36 | 1 | 2 | 0 | 0 |
| insula posterior R | Ctx PI R | inferior dysgranular insula R | Cortex_Ventral_AttentionA | 1528 | 44 | -3 | -12 | 7 | 8 | 1 | -9.8 | 0 | 3 | 0 | 0 |
| insula posterior R | Ctx PoI1 R | inferior dysgranular insula R | Cortex_Ventral_AttentionA | 2048 | 38 | -8 | -6 | 11.18 | 9 | 1 | -6.2 | 0 | 2 | 0 | 0 |
| insula posterior R | Ctx PoI2 R | inferior dysgranular insula R | Cortex_Ventral_AttentionA | 3640 | 40 | -0 | -0 | 13.17 | 9 | 1 | -2.35 | 2 | 2 | 3 | 1 |
| parietal TPOJ L | Ctx PSL L | PSL L | Cortex_Ventral_AttentionA | 2768 | -56 | -44 | 23 | 2.98 | 2 | 4 | -3.83 | 3 | 3 | 1 | 2 |
| parietal TPOJ L | Ctx STV L | STV L | Cortex_Temporal_Parietal | 2368 | -61 | -50 | 17 | -8.01 | 1 | 4 | -3.72 | 3 | 2 | 0 | 3 |
| parietal TPOJ L | Ctx TPOJ1 L | TPOJ L | Cortex_Temporal_Parietal | 2976 | -52 | -45 | 10 | -5.04 | 0 | 5 | -5.03 | 2 | 3 | 0 | 2 |
| parietal TPOJ L | Ctx TPOJ2 L | TPOJ L | Cortex_Dorsal_AttentionA | 3152 | -52 | -61 | 12 | -7.77 | 1 | 6 | -6.86 | 1 | 4 | 1 | 3 |
| parietal TPOJ L | Ctx TPOJ3 L | TPOJ L | Cortex_Dorsal_AttentionA | 1264 | -42 | -71 | 19 | -6.51 | 0 | 4 | -4.3 | 0 | 3 | 0 | 2 |
| parietal TPOJ R | Ctx PSL R | PSL R | Cortex_Temporal_Parietal | 3128 | 64 | -35 | 26 | 15.18 | 7 | 0 | -5.17 | 2 | 2 | 1 | 2 |
| parietal TPOJ R | Ctx STV R | STV R | Cortex_Temporal_Parietal | 3384 | 59 | -42 | 19 | -2.36 | 3 | 3 | -4.76 | 2 | 3 | 2 | 2 |
| parietal TPOJ R | Ctx TPOJ1 R | TPOJ R | Cortex_Temporal_Parietal | 6104 | 53 | -42 | 10 | -6.9 | 2 | 3 | -9.04 | 2 | 2 | 1 | 2 |
| parietal TPOJ R | Ctx TPOJ2 R | TPOJ R | Cortex_Dorsal_AttentionA | 3376 | 53 | -57 | 8 | -10.59 | 1 | 4 | -10.29 | 1 | 2 | 1 | 1 |
| parietal TPOJ R | Ctx TPOJ3 R | TPOJ R | Cortex_Default_ModeC | 2696 | 42 | -62 | 17 | -7.02 | 0 | 4 | -14 | 0 | 3 | 0 | 2 |
| parietal inferior lobule L | Ctx IP0 L | posterior IPL L | Cortex_Dorsal_AttentionA | 2192 | -31 | -75 | 27 | -5.17 | 0 | 4 | -4.79 | 0 | 6 | 0 | 3 |
| parietal inferior lobule L | Ctx IP1 L | superior IPL L | Cortex_Fronto_ParietalA | 3040 | -31 | -68 | 40 | -3.94 | 0 | 4 | 5.17 | 1 | 1 | 0 | 1 |
| parietal inferior lobule L | Ctx IP2 L | superior IPL L | Cortex_Fronto_ParietalA | 3728 | -39 | -51 | 39 | 4.75 | 3 | 1 | 3.49 | 4 | 0 | 1 | 0 |
| parietal inferior lobule L | Ctx PF L | anterior IPL L | Cortex_Ventral_AttentionA | 7520 | -58 | -38 | 39 | 11.05 | 6 | 0 | -5.25 | 4 | 1 | 3 | 0 |
| parietal inferior lobule L | Ctx PFm L | middle IPL L | Cortex_Fronto_ParietalB | 9640 | -49 | -56 | 44 | -7.17 | 2 | 2 | 3.54 | 4 | 0 | 1 | 0 |
| parietal inferior lobule L | Ctx PFop L | anterior IPL L | Cortex_Dorsal_AttentionB | 3080 | -62 | -23 | 27 | 8.35 | 9 | 1 | -1.96 | 4 | 2 | 4 | 0 |
| parietal inferior lobule L | Ctx PFt L | anterior IPL L | Cortex_Dorsal_AttentionB | 3664 | -53 | -27 | 38 | -3.81 | 3 | 3 | 3.59 | 2 | 3 | 2 | 1 |

| | | | | | MNI Coordinates | | | Experienced Pain (Hot-Warm) | | | Imagined Pain (Imagine-Warm) | | | Pain Conjunction | |
|---|---|---|---|---|---|---|---|---|---|---|---|---|---|---|---|
| Structure | Region | Parcel | Network | Volume | X | Y | Z | T | Activated (n) | Deactivated (n) | T | Activated (n) | Deactivated (n) | Activated (n) | Deactivated (n) |
| parietal inferior lobule L | Ctx PGi L | inferior IPL L | Cortex_Default_ModeB | 7832 | -45 | -59 | 24 | -22.86 | 0 | 6 | 3.81 | 2 | 0 | 0 | 0 |
| parietal inferior lobule L | Ctx PGp L | posterior IPL L | Cortex_Visual_Central | 3016 | -37 | -84 | 25 | -7.46 | 0 | 4 | -15.25 | 0 | 6 | 0 | 4 |
| parietal inferior lobule L | Ctx PGs L | inferior IPL L | Cortex_Default_ModeA | 5256 | -40 | -74 | 39 | -10.14 | 0 | 4 | -12.2 | 1 | 2 | 0 | 1 |
| parietal inferior lobule R | Ctx IP0 R | posterior IPL R | Cortex_Dorsal_AttentionA | 1936 | 34 | -69 | 31 | -6.72 | 0 | 3 | -4.2 | 0 | 5 | 0 | 3 |
| parietal inferior lobule R | Ctx IP1 R | superior IPL R | Cortex_Dorsal_AttentionA | 4000 | 35 | -63 | 44 | 7.52 | 1 | 3 | -6.49 | 0 | 4 | 0 | 2 |
| parietal inferior lobule R | Ctx IP2 R | superior IPL R | Cortex_Fronto_ParietalA | 3648 | 43 | -43 | 43 | 6.54 | 4 | 2 | -5.55 | 0 | 3 | 0 | 0 |
| parietal inferior lobule R | Ctx PF R | anterior IPL R | Cortex_Ventral_AttentionA | 5704 | 60 | -29 | 38 | 16.91 | 7 | 0 | -5.65 | 1 | 3 | 2 | 1 |
| parietal inferior lobule R | Ctx PFm R | middle IPL R | Cortex_Fronto_ParietalB | 10704 | 53 | -47 | 45 | 12.37 | 4 | 1 | -6.16 | 1 | 3 | 0 | 0 |
| parietal inferior lobule R | Ctx PFop R | anterior IPL R | Cortex_Ventral_AttentionA | 2608 | 63 | -18 | 26 | 7.85 | 8 | 0 | -4.15 | 0 | 2 | 0 | 1 |
| parietal inferior lobule R | Ctx PFt R | anterior IPL R | Cortex_Dorsal_AttentionB | 2968 | 56 | -21 | 40 | 7.6 | 1 | 5 | 3.1 | 1 | 2 | 1 | 1 |
| parietal inferior lobule R | Ctx PGi R | inferior IPL R | Cortex_Default_ModeC | 8568 | 50 | -57 | 26 | -10.99 | 0 | 5 | -6.84 | 0 | 3 | 0 | 2 |
| parietal inferior lobule R | Ctx PGp R | posterior IPL R | Cortex_Default_ModeC | 4872 | 42 | -78 | 27 | -11.62 | 0 | 5 | -7.44 | 0 | 5 | 0 | 3 |
| parietal inferior lobule R | Ctx PGs R | inferior IPL R | Cortex_Default_ModeA | 4424 | 45 | -67 | 44 | 8.95 | 1 | 2 | -5.1 | 0 | 4 | 0 | 1 |
| parietal superior lobule L | Ctx 7AL L | anterior BA7 L | Cortex_Dorsal_AttentionB | 3264 | -20 | -51 | 69 | -5.58 | 1 | 4 | -4.71 | 1 | 3 | 0 | 2 |
| parietal superior lobule L | Ctx 7Am L | anterior BA7 L | Cortex_Dorsal_AttentionB | 4384 | -8 | -59 | 63 | -4.09 | 1 | 4 | -5.04 | 1 | 2 | 0 | 2 |
| parietal superior lobule L | Ctx 7PC L | postcentral BA7 L | Cortex_Dorsal_AttentionB | 4136 | -36 | -48 | 63 | -12.5 | 0 | 5 | -6.6 | 1 | 3 | 0 | 2 |
| parietal superior lobule L | Ctx 7PL L | posterior BA7 L | Cortex_Dorsal_AttentionA | 1952 | -14 | -72 | 57 | -3.29 | 0 | 5 | 2.53 | 2 | 2 | 0 | 2 |
| parietal superior lobule L | Ctx 7Pm L | posterior BA7 L | Cortex_Fronto_ParietalC | 2016 | -5 | -68 | 53 | -4.94 | 2 | 2 | 4.31 | 0 | 3 | 0 | 2 |
| parietal superior lobule L | Ctx AIP L | ventral intraparietal L | Cortex_Fronto_ParietalA | 5032 | -37 | -40 | 44 | -3.23 | 1 | 4 | 3.97 | 2 | 0 | 0 | 0 |
| parietal superior lobule L | Ctx LIPd L | ventral intraparietal L | Cortex_Fronto_ParietalA | 2240 | -28 | -55 | 41 | 2.78 | 0 | 2 | 3.19 | 4 | 1 | 0 | 0 |
| parietal superior lobule L | Ctx LIPv L | dorsal intraparietal L | Cortex_Dorsal_AttentionA | 2848 | -30 | -58 | 56 | -8.43 | 0 | 5 | -5.67 | 1 | 3 | 0 | 2 |
| parietal superior lobule L | Ctx MIP L | MIP L | Cortex_Dorsal_AttentionA | 3160 | -23 | -64 | 47 | -3.84 | 0 | 4 | 4.31 | 2 | 2 | 0 | 1 |
| parietal superior lobule L | Ctx VIP L | dorsal intraparietal L | Cortex_Dorsal_AttentionA | 2776 | -23 | -64 | 63 | -6.32 | 0 | 5 | -3.98 | 1 | 3 | 0 | 3 |
| parietal superior lobule R | Ctx 7AL R | anterior BA7 R | Cortex_Dorsal_AttentionB | 3016 | 23 | -50 | 69 | -5.89 | 2 | 3 | -5.29 | 0 | 3 | 0 | 2 |
| parietal superior lobule R | Ctx 7Am R | anterior BA7 R | Cortex_Dorsal_AttentionB | 4008 | 10 | -58 | 65 | -12.7 | 0 | 5 | -6.17 | 1 | 2 | 0 | 2 |
| parietal superior lobule R | Ctx 7PC R | postcentral BA7 R | Cortex_Dorsal_AttentionA | 4904 | 35 | -47 | 63 | -13.16 | 1 | 6 | -5.9 | 0 | 5 | 0 | 2 |
| parietal superior lobule R | Ctx 7PL R | posterior BA7 R | Cortex_Fronto_ParietalC | 1600 | 17 | -69 | 59 | -4.26 | 0 | 5 | -4.07 | 2 | 2 | 0 | 1 |

| Structure | Region | Parcel | Network | Volume | MNI Coordinates | | | Experienced Pain (Hot-Warm) | | | Imagined Pain (Imagine-Warm) | | | Pain Conjunction | |
|---|---|---|---|---|---|---|---|---|---|---|---|---|---|---|---|
| | | | | | X | Y | Z | T | Activated (n) | Deactivated (n) | T | Activated (n) | Deactivated (n) | Activated (n) | Deactivated (n) |
| parietal superior lobule R | Ctx 7Pm R | posterior BA7 R | Cortex_Fronto_ParietalC | 2088 | 7 | -67 | 55 | -7.59 | 1 | 4 | -2.85 | 0 | 3 | 0 | 2 |
| parietal superior lobule R | Ctx AIP R | ventral intraparietal R | Cortex_Fronto_ParietalA | 5640 | 37 | -38 | 45 | -8 | 1 | 6 | 2.84 | 1 | 3 | 0 | 1 |
| parietal superior lobule R | Ctx LIPd R | ventral intraparietal R | Cortex_Dorsal_AttentionA | 1128 | 30 | -51 | 42 | 2.59 | 1 | 2 | -2.32 | 1 | 2 | 0 | 1 |
| parietal superior lobule R | Ctx LIPv R | dorsal intraparietal R | Cortex_Dorsal_AttentionA | 2024 | 28 | -55 | 55 | -7.44 | 0 | 5 | -3.65 | 0 | 4 | 0 | 3 |
| parietal superior lobule R | Ctx MIP R | MIP R | Cortex_Dorsal_AttentionA | 3672 | 26 | -64 | 50 | -4.48 | 0 | 5 | -4.86 | 0 | 4 | 0 | 2 |
| parietal superior lobule R | Ctx VIP R | dorsal intraparietal R | Cortex_Dorsal_AttentionA | 1784 | 23 | -61 | 64 | -6.27 | 0 | 5 | -4.4 | 0 | 3 | 0 | 2 |
| somatomotor operculum L | Ctx 43 L | FOP1-BA43 L | Cortex_SomatomotorB | 2160 | -57 | -0 | 11 | 7.72 | 9 | 1 | 4.79 | 6 | 0 | 6 | 1 |
| somatomotor operculum L | Ctx FOP1 L | FOP1-BA43 L | Cortex_Ventral_AttentionA | 1008 | -50 | 2 | 5 | 11.32 | 8 | 0 | 3.03 | 4 | 1 | 3 | 0 |
| somatomotor operculum L | Ctx OP1 L | SII+ L | Cortex_SomatomotorB | 2704 | -48 | -21 | 21 | 8.55 | 7 | 1 | -3.73 | 0 | 3 | 0 | 1 |
| somatomotor operculum L | Ctx OP2 3 L | SII+ L | Cortex_SomatomotorB | 1552 | -40 | -16 | 20 | 7.76 | 8 | 0 | 3.48 | 0 | 2 | 0 | 0 |
| somatomotor operculum L | Ctx OP4 L | OP4 L | Cortex_SomatomotorB | 2784 | -59 | -13 | 16 | 10.94 | 9 | 1 | -5.5 | 3 | 2 | 3 | 1 |
| somatomotor operculum L | Ctx PFcm L | SII+ L | Cortex_Ventral_AttentionA | 2360 | -51 | -32 | 22 | 7.16 | 7 | 1 | -4.49 | 1 | 3 | 0 | 2 |
| somatomotor operculum R | Ctx 43 R | FOP1-BA43 R | Cortex_SomatomotorB | 3184 | 56 | 1 | 11 | 14.88 | 9 | 1 | 7.65 | 4 | 1 | 4 | 0 |
| somatomotor operculum R | Ctx FOP1 R | FOP1-BA43 R | Cortex_Ventral_AttentionA | 1216 | 47 | 4 | 7 | 9.19 | 9 | 1 | 2.87 | 4 | 1 | 4 | 1 |
| somatomotor operculum R | Ctx OP1 R | SII+ R | Cortex_SomatomotorB | 1456 | 44 | -19 | 21 | 11.42 | 7 | 1 | -1.68 | 0 | 2 | 0 | 0 |
| somatomotor operculum R | Ctx OP2 3 R | SII+ R | Cortex_SomatomotorB | 1248 | 39 | -12 | 19 | 12.78 | 9 | 1 | 2.82 | 1 | 2 | 1 | 0 |
| somatomotor operculum R | Ctx OP4 R | OP4 R | Cortex_SomatomotorB | 3664 | 58 | -11 | 16 | 8.03 | 8 | 0 | -7.64 | 0 | 2 | 0 | 0 |
| somatomotor operculum R | Ctx PFcm R | SII+ R | Cortex_SomatomotorB | 2472 | 49 | -26 | 24 | 5.71 | 7 | 1 | -2.55 | 0 | 4 | 0 | 1 |
| somatomotor paracentral lobule L | Ctx 24dd L | cingulate motor area L | Cortex_SomatomotorA | 4440 | -7 | -16 | 51 | -3.24 | 2 | 3 | 4.72 | 3 | 1 | 2 | 1 |
| somatomotor paracentral lobule L | Ctx 24dv L | cingulate motor area L | Cortex_Ventral_AttentionA | 2112 | -10 | -2 | 46 | 5.59 | 4 | 0 | 2.83 | 4 | 1 | 3 | 0 |
| somatomotor paracentral lobule L | Ctx 5L L | BA5 L | Cortex_SomatomotorA | 2680 | -13 | -46 | 72 | -3.22 | 2 | 3 | 2.33 | 0 | 2 | 0 | 1 |
| somatomotor paracentral lobule L | Ctx 5m L | BA5 L | Cortex_SomatomotorA | 2200 | -6 | -40 | 63 | -2.94 | 1 | 4 | -3.26 | 2 | 3 | 1 | 2 |
| somatomotor paracentral lobule L | Ctx 5mv L | BA5 L | Cortex_Ventral_AttentionA | 2216 | -14 | -36 | 51 | -2.09 | 2 | 3 | 2.9 | 1 | 2 | 0 | 2 |
| somatomotor paracentral lobule L | Ctx 6ma L | SMA L | Cortex_Fronto_ParietalA | 5680 | -19 | 4 | 69 | -5.38 | 3 | 1 | 4.12 | 5 | 0 | 2 | 0 |
| somatomotor paracentral lobule L | Ctx 6mp L | SMA L | Cortex_SomatomotorA | 6432 | -12 | -15 | 71 | -12.07 | 4 | 3 | 5.05 | 5 | 1 | 3 | 1 |
| somatomotor paracentral lobule L | Ctx SCEF L | SMA L | Cortex_Ventral_AttentionA | 5824 | -6 | 4 | 59 | 6.01 | 4 | 0 | 4.57 | 7 | 0 | 3 | 0 |
| somatomotor paracentral lobule R | Ctx 24dd R | cingulate motor area R | Cortex_SomatomotorA | 4864 | 8 | -17 | 53 | -4.93 | 2 | 3 | 5.95 | 3 | 1 | 2 | 1 |

| Structure | Region | Parcel | Network | Volume | MNI Coordinates X | Y | Z | Experienced Pain (Hot-Warm) T | Activated (n) | Deactivated (n) | Imagined Pain (Imagine-Warm) T | Activated (n) | Deactivated (n) | Pain Conjunction Activated (n) | Deactivated (n) |
|---|---|---|---|---|---|---|---|---|---|---|---|---|---|---|---|
| somatomotor paracentral lobule R | Ctx 24dv R | cingulate motor area R | Cortex_Ventral_AttentionA | 2336 | 11 | -4 | 47 | 5.44 | 4 | 2 | 8.46 | 3 | 1 | 2 | 0 |
| somatomotor paracentral lobule R | Ctx 5L R | BA5 R | Cortex_Dorsal_AttentionB | 2936 | 13 | -45 | 75 | -2.78 | 3 | 3 | -3.93 | 0 | 2 | 0 | 1 |
| somatomotor paracentral lobule R | Ctx 5m R | BA5 R | Cortex_SomatomotorA | 2896 | 6 | -38 | 66 | -2.87 | 1 | 4 | 4.44 | 2 | 2 | 1 | 1 |
| somatomotor paracentral lobule R | Ctx 5mv R | BA5 R | Cortex_Ventral_AttentionA | 3000 | 14 | -39 | 54 | -7.21 | 1 | 4 | 4.8 | 2 | 2 | 1 | 1 |
| somatomotor paracentral lobule R | Ctx 6ma R | SMA R | Cortex_Ventral_AttentionA | 5728 | 20 | 3 | 68 | -10.89 | 5 | 0 | 6.56 | 3 | 3 | 2 | 1 |
| somatomotor paracentral lobule R | Ctx 6mp R | SMA R | Cortex_SomatomotorA | 5808 | 17 | -12 | 68 | -6.19 | 1 | 4 | 5.07 | 4 | 1 | 1 | 1 |
| somatomotor paracentral lobule R | Ctx SCEF R | SMA R | Cortex_Ventral_AttentionA | 4776 | 7 | 3 | 59 | 7.16 | 6 | 1 | 5.56 | 6 | 0 | 4 | 0 |
| somatomotor premotor L | Ctx 55b L | 55b L | Cortex_Default_ModeB | 3016 | -48 | -1 | 50 | -4.62 | 1 | 3 | -3.94 | 4 | 1 | 3 | 0 |
| somatomotor premotor L | Ctx 6a L | superior premotor L | Cortex_Dorsal_AttentionB | 8120 | -25 | -3 | 54 | -18.74 | 1 | 3 | 6.48 | 3 | 1 | 0 | 1 |
| somatomotor premotor L | Ctx 6d L | superior premotor L | Cortex_SomatomotorA | 4200 | -34 | -14 | 67 | -10.13 | 0 | 5 | 3.9 | 1 | 3 | 0 | 1 |
| somatomotor premotor L | Ctx 6r L | inferior premotor L | Cortex_Ventral_AttentionA | 4360 | -52 | 6 | 19 | 8.77 | 6 | 0 | 7.1 | 5 | 0 | 4 | 0 |
| somatomotor premotor L | Ctx 6v L | inferior premotor L | Cortex_SomatomotorB | 2096 | -59 | 4 | 33 | -4.55 | 5 | 0 | 7.62 | 5 | 1 | 3 | 0 |
| somatomotor premotor L | Ctx FEF L | superior premotor L | Cortex_Dorsal_AttentionB | 2888 | -39 | -5 | 50 | -4.29 | 2 | 1 | 4.49 | 6 | 0 | 2 | 0 |
| somatomotor premotor L | Ctx PEF L | inferior premotor L | Cortex_Dorsal_AttentionB | 1272 | -48 | -1 | 39 | -2.54 | 1 | 2 | 3.7 | 2 | 3 | 1 | 1 |
| somatomotor premotor R | Ctx 55b R | 55b R | Cortex_Ventral_AttentionA | 1880 | 51 | 3 | 48 | -3.61 | 4 | 2 | 3.59 | 3 | 2 | 3 | 0 |
| somatomotor premotor R | Ctx 6a R | superior premotor R | Cortex_Dorsal_AttentionB | 7056 | 27 | -2 | 52 | -5.82 | 1 | 4 | 6.27 | 3 | 2 | 1 | 0 |
| somatomotor premotor R | Ctx 6d R | superior premotor R | Cortex_SomatomotorA | 3784 | 37 | -12 | 65 | -8.4 | 1 | 5 | 3.65 | 2 | 2 | 1 | 2 |
| somatomotor premotor R | Ctx 6r R | inferior premotor R | Cortex_Ventral_AttentionA | 5496 | 52 | 10 | 15 | 10.85 | 8 | 0 | -6.95 | 5 | 0 | 4 | 0 |
| somatomotor premotor R | Ctx 6v R | inferior premotor R | Cortex_SomatomotorB | 3696 | 59 | 7 | 31 | 4.86 | 5 | 1 | 4.01 | 4 | 1 | 3 | 0 |
| somatomotor premotor R | Ctx FEF R | superior premotor R | Cortex_Dorsal_AttentionB | 3128 | 43 | -1 | 51 | -4.16 | 3 | 1 | 2.8 | 5 | 1 | 2 | 1 |
| somatomotor premotor R | Ctx PEF R | inferior premotor R | Cortex_Fronto_ParietalA | 1640 | 47 | 3 | 37 | 2.2 | 4 | 1 | 2.46 | 3 | 2 | 1 | 1 |
| somatomotor primary L | Ctx 1 L | S1 L | Cortex_SomatomotorA | 8384 | -44 | -26 | 60 | -12.14 | 1 | 5 | 3.31 | 1 | 3 | 0 | 2 |
| somatomotor primary L | Ctx 2 L | S1 L | Cortex_Dorsal_AttentionB | 8488 | -36 | -34 | 54 | -11.77 | 1 | 5 | 5.8 | 3 | 2 | 0 | 2 |
| somatomotor primary L | Ctx 3a L | S1 L | Cortex_SomatomotorA | 4752 | -33 | -21 | 44 | -8.55 | 1 | 4 | 19.14 | 3 | 1 | 1 | 0 |
| somatomotor primary L | Ctx 3b L | S1 L | Cortex_SomatomotorA | 8320 | -35 | -24 | 55 | -7.49 | 1 | 4 | 6.97 | 3 | 1 | 1 | 2 |
| somatomotor primary L | Ctx 4 L | M1 L | Cortex_SomatomotorA | 14672 | -26 | -19 | 56 | -8.21 | 1 | 4 | 9.27 | 4 | 0 | 1 | 1 |
| somatomotor primary R | Ctx 1 R | S1 R | Cortex_SomatomotorA | 8464 | 46 | -22 | 58 | -17.65 | 1 | 6 | -2.23 | 2 | 2 | 1 | 2 |

| Structure | Region | Parcel | Network | Volume | MNI Coordinates | | | Experienced Pain (Hot-Warm) | | | Imagined Pain (Imagine-Warm) | | | Pain Conjunction | |
|---|---|---|---|---|---|---|---|---|---|---|---|---|---|---|---|
| | | | | | X | Y | Z | T | Activated (n) | Deactivated (n) | T | Activated (n) | Deactivated (n) | Activated (n) | Deactivated (n) |
| somatomotor primary R | Ctx 2 R | S1 R | Cortex_Dorsal_AttentionB | 8240 | 36 | -31 | 54 | -24.84 | 2 | 6 | 10.09 | 2 | 1 | 1 | 1 |
| somatomotor primary R | Ctx 3a R | S1 R | Cortex_SomatomotorA | 4648 | 32 | -20 | 46 | -5.17 | 1 | 4 | 5.17 | 3 | 1 | 1 | 1 |
| somatomotor primary R | Ctx 3b R | S1 R | Cortex_SomatomotorA | 6840 | 36 | -21 | 54 | -11.29 | 1 | 5 | 4.5 | 3 | 1 | 1 | 1 |
| somatomotor primary R | Ctx 4 R | M1 R | Cortex_SomatomotorA | 13608 | 27 | -17 | 58 | -7.9 | 1 | 5 | 7.61 | 3 | 1 | 1 | 1 |
| temporal lateral L | Ctx PHT L | middle temporal gyrus L | Cortex_Fronto_ParietalA | 5224 | -58 | -59 | 0 | -10.29 | 0 | 4 | -3.47 | 2 | 3 | 0 | 2 |
| temporal lateral L | Ctx TE1a L | temporal pole L | Cortex_Default_ModeB | 5368 | -61 | -7 | -20 | -11.26 | 0 | 5 | -4.78 | 0 | 6 | 0 | 4 |
| temporal lateral L | Ctx TE1m L | middle temporal gyrus L | Cortex_Fronto_ParietalB | 3888 | -64 | -26 | -14 | -6.58 | 0 | 4 | -3.73 | 1 | 2 | 0 | 1 |
| temporal lateral L | Ctx TE1p L | middle temporal gyrus L | Cortex_Fronto_ParietalB | 8568 | -61 | -46 | -11 | -2.91 | 1 | 3 | -3.98 | 2 | 1 | 0 | 0 |
| temporal lateral L | Ctx TE2a L | inferior temporal sulcus L | Cortex_Limbic | 7464 | -56 | -24 | -27 | 6.77 | 0 | 2 | -3.39 | 0 | 0 | 0 | 0 |
| temporal lateral L | Ctx TE2p L | inferior temporal sulcus L | Cortex_Dorsal_AttentionA | 4736 | -47 | -42 | -19 | -5.06 | 0 | 2 | 9.2 | 2 | 2 | 0 | 1 |
| temporal lateral L | Ctx TF L | inferior temporal sulcus L | Cortex_Limbic | 7104 | -42 | -24 | -26 | -8.13 | 0 | 5 | 11.78 | 1 | 1 | 0 | 0 |
| temporal lateral L | Ctx TGd L | temporal pole L | Cortex_Default_ModeB | 15600 | -40 | 10 | -32 | -14.25 | 0 | 5 | 7.92 | 1 | 2 | 0 | 0 |
| temporal lateral L | Ctx TGv L | temporal pole L | Cortex_Limbic | 5984 | -38 | -2 | -43 | -9.4 | 0 | 3 | 10.78 | 0 | 0 | 0 | 0 |
| temporal lateral R | Ctx PHT R | middle temporal gyrus R | Cortex_Dorsal_AttentionB | 4912 | 60 | -55 | -3 | -5.14 | 1 | 4 | -6.8 | 1 | 1 | 1 | 0 |
| temporal lateral R | Ctx TE1a R | temporal pole R | Cortex_Default_ModeA | 5136 | 61 | -2 | -23 | -13.38 | 0 | 5 | -9.44 | 0 | 4 | 0 | 3 |
| temporal lateral R | Ctx TE1m R | middle temporal gyrus R | Cortex_Fronto_ParietalB | 4112 | 65 | -23 | -15 | -9.7 | 0 | 4 | -3.81 | 0 | 4 | 0 | 2 |
| temporal lateral R | Ctx TE1p R | middle temporal gyrus R | Cortex_Fronto_ParietalB | 8528 | 62 | -42 | -12 | -5.59 | 0 | 2 | -4.55 | 1 | 2 | 0 | 0 |
| temporal lateral R | Ctx TE2a R | inferior temporal sulcus R | Cortex_Limbic | 8192 | 55 | -17 | -29 | -6.9 | 0 | 3 | -4.65 | 0 | 3 | 0 | 0 |
| temporal lateral R | Ctx TE2p R | inferior temporal sulcus R | Cortex_Dorsal_AttentionA | 4552 | 48 | -37 | -19 | -4.64 | 2 | 2 | -5.91 | 3 | 0 | 0 | 0 |
| temporal lateral R | Ctx TF R | inferior temporal sulcus R | Cortex_Limbic | 7096 | 41 | -22 | -27 | -6.28 | 0 | 5 | -8.25 | 0 | 2 | 0 | 0 |
| temporal lateral R | Ctx TGd R | temporal pole R | Cortex_Limbic | 13248 | 38 | 14 | -34 | -10.81 | 0 | 3 | -5.9 | 0 | 1 | 0 | 0 |
| temporal lateral R | Ctx TGv R | temporal pole R | Cortex_Limbic | 6256 | 36 | -1 | -45 | -12.5 | 0 | 3 | 3.44 | 0 | 1 | 0 | 0 |
| temporal medial L | Ctx EC L | hippocampal formation L | Cortex_Limbic | 2824 | -21 | -12 | -31 | -5.04 | 0 | 5 | 3.66 | 0 | 1 | 0 | 1 |
| temporal medial L | Ctx PHA1 L | PHA L | Cortex_Default_ModeC | 2200 | -23 | -36 | -15 | -7.58 | 0 | 5 | -3.52 | 0 | 1 | 0 | 1 |
| temporal medial L | Ctx PHA2 L | PHA L | Cortex_Default_ModeC | 688 | -31 | -38 | -11 | -7.91 | 0 | 5 | -4.35 | 0 | 4 | 0 | 3 |
| temporal medial L | Ctx PHA3 L | PHA L | Cortex_Default_ModeC | 2352 | -31 | -38 | -17 | -12.16 | 0 | 5 | -7.14 | 0 | 5 | 0 | 4 |
| temporal medial L | Ctx PeEc L | PeEc L | Cortex_Limbic | 8280 | -31 | -9 | -32 | -4 | 0 | 4 | 5.63 | 0 | 1 | 0 | 1 |
| temporal medial L | Ctx PreS L | hippocampal formation L | Cortex_Default_ModeC | 2056 | -16 | -32 | -12 | -7.67 | 0 | 4 | 3.84 | 0 | 2 | 0 | 0 |
| temporal medial R | Ctx EC R | hippocampal formation R | Cortex_Default_ModeC | 2272 | 20 | -12 | -30 | -7.59 | 0 | 5 | -5.27 | 0 | 2 | 0 | 1 |
| temporal medial R | Ctx PHA1 R | PHA R | Cortex_Visual_Peripheral | 1840 | 23 | -34 | -16 | -7.86 | 0 | 6 | -7.15 | 0 | 3 | 0 | 2 |
| temporal medial R | Ctx PHA2 R | PHA R | Cortex_Default_ModeC | 672 | 31 | -34 | -13 | -5.41 | 0 | 6 | -4.99 | 0 | 2 | 0 | 1 |
| temporal medial R | Ctx PHA3 R | PHA R | Cortex_Default_ModeC | 784 | 35 | -36 | -14 | -4.96 | 0 | 6 | -4.75 | 0 | 3 | 0 | 1 |

| Structure | Region | Parcel | Network | Volume | MNI Coordinates X | Y | Z | Experienced Pain (Hot-Warm) T | Activated (n) | Deactivated (n) | Imagined Pain (Imagine-Warm) T | Activated (n) | Deactivated (n) | Pain Conjunction Activated (n) | Deactivated (n) |
|---|---|---|---|---|---|---|---|---|---|---|---|---|---|---|---|
| temporal medial R | Ctx PeEc R | PeEc R | Cortex_Limbic | 7112 | 28 | -7 | -33 | -6.78 | 0 | 5 | -4.49 | 0 | 1 | 0 | 0 |
| temporal medial R | Ctx PreS R | hippocampal formation R | Cortex_Visual_Peripheral | 1664 | 16 | -33 | -10 | -5.58 | 0 | 5 | -9.88 | 0 | 2 | 0 | 1 |
| visual MT+ L | Ctx FST L | vMT+ L | Cortex_Fronto_ParietalA | 1792 | -47 | -66 | 3 | -5.09 | 0 | 4 | -10.94 | 0 | 5 | 0 | 4 |
| visual MT+ L | Ctx LO1 L | LO L | Cortex_Visual_Central | 1416 | -37 | -82 | 6 | -3.31 | 0 | 3 | -14.72 | 0 | 6 | 0 | 3 |
| visual MT+ L | Ctx LO2 L | LO L | Cortex_Visual_Central | 1608 | -42 | -85 | -3 | -3.73 | 0 | 4 | -6.11 | 0 | 7 | 0 | 4 |
| visual MT+ L | Ctx LO3 L | LO L | Cortex_Visual_Central | 1368 | -46 | -79 | 13 | -4 | 0 | 5 | -11.95 | 0 | 5 | 0 | 3 |
| visual MT+ L | Ctx MST L | dMT+ L | Cortex_Visual_Central | 1272 | -39 | -66 | 7 | -4.3 | 1 | 3 | -11.93 | 2 | 3 | 1 | 1 |
| visual MT+ L | Ctx MT L | dMT+ L | Cortex_Visual_Central | 1120 | -43 | -73 | 10 | -5.04 | 0 | 5 | -8.93 | 0 | 4 | 0 | 2 |
| visual MT+ L | Ctx PH L | PH L | Cortex_Dorsal_AttentionA | 5760 | -45 | -65 | -7 | -7.7 | 0 | 4 | -8.9 | 1 | 5 | 0 | 3 |
| visual MT+ L | Ctx V4t L | vMT+ L | Cortex_Visual_Central | 1176 | -47 | -78 | 1 | -3.59 | 0 | 5 | -6.43 | 0 | 6 | 0 | 4 |
| visual MT+ R | Ctx FST R | vMT+ R | Cortex_Dorsal_AttentionA | 1912 | 46 | -61 | -1 | -4.35 | 0 | 5 | -8.2 | 0 | 5 | 0 | 4 |
| visual MT+ R | Ctx LO1 R | LO R | Cortex_Visual_Central | 1320 | 37 | -78 | 6 | -6.58 | 0 | 4 | -9.94 | 0 | 5 | 0 | 3 |
| visual MT+ R | Ctx LO2 R | LO R | Cortex_Visual_Central | 1120 | 44 | -84 | -0 | -4.15 | 0 | 4 | -8.49 | 0 | 5 | 0 | 4 |
| visual MT+ R | Ctx LO3 R | LO R | Cortex_Visual_Central | 1312 | 45 | -75 | 12 | -14.8 | 0 | 5 | -13.97 | 0 | 5 | 0 | 3 |
| visual MT+ R | Ctx MST R | dMT+ R | Cortex_Dorsal_AttentionA | 1584 | 42 | -65 | 5 | -5.29 | 0 | 4 | -8.66 | 0 | 4 | 0 | 2 |
| visual MT+ R | Ctx MT R | dMT+ R | Cortex_Dorsal_AttentionA | 1464 | 50 | -70 | 8 | -4.95 | 0 | 5 | -7.62 | 0 | 3 | 0 | 2 |
| visual MT+ R | Ctx PH R | PH R | Cortex_Dorsal_AttentionA | 5216 | 47 | -63 | -10 | -10.34 | 0 | 5 | -10.52 | 0 | 6 | 0 | 3 |
| visual MT+ R | Ctx V4t R | vMT+ R | Cortex_Dorsal_AttentionA | 1152 | 50 | -75 | -1 | -4.59 | 0 | 5 | -5.63 | 0 | 6 | 0 | 4 |
| visual dorsal L | Ctx IPS1 L | ISP1 L | Cortex_Dorsal_AttentionA | 2424 | -24 | -73 | 36 | -6.06 | 0 | 4 | 4.42 | 1 | 2 | 0 | 2 |
| visual dorsal L | Ctx V3A L | supplementary V3 L | Cortex_Visual_Central | 3376 | -17 | -89 | 27 | -3.39 | 0 | 3 | -9.69 | 1 | 4 | 0 | 3 |
| visual dorsal L | Ctx V3B L | supplementary V3 L | Cortex_Visual_Central | 1520 | -26 | -79 | 18 | -3.62 | 0 | 3 | -6.36 | 0 | 5 | 0 | 3 |
| visual dorsal L | Ctx V3CD L | supplementary V3 L | Cortex_Visual_Central | 1488 | -35 | -86 | 13 | -4.15 | 0 | 4 | -6.01 | 0 | 6 | 0 | 4 |
| visual dorsal L | Ctx V6 L | V6 L | Cortex_Visual_Peripheral | 2024 | -15 | -79 | 29 | -2.68 | 1 | 3 | -2.3 | 2 | 3 | 1 | 2 |
| visual dorsal L | Ctx V6A L | V6 L | Cortex_Visual_Peripheral | 1048 | -21 | -85 | 43 | -3.52 | 0 | 4 | -3.19 | 1 | 3 | 0 | 2 |
| visual dorsal L | Ctx V7 L | V7 L | Cortex_Visual_Central | 1184 | -24 | -84 | 31 | -3.85 | 0 | 5 | -3.87 | 0 | 4 | 0 | 3 |
| visual dorsal R | Ctx IPS1 R | ISP1 R | Cortex_Dorsal_AttentionA | 2544 | 27 | -69 | 38 | -2.75 | 0 | 3 | -5.48 | 0 | 5 | 0 | 2 |
| visual dorsal R | Ctx V3A R | supplementary V3 R | Cortex_Visual_Peripheral | 3960 | 18 | -87 | 32 | -3.75 | 1 | 4 | -10.55 | 1 | 4 | 0 | 3 |
| visual dorsal R | Ctx V3B R | supplementary V3 R | Cortex_Dorsal_AttentionA | 1696 | 29 | -74 | 22 | -5.61 | 0 | 2 | -4.09 | 1 | 4 | 0 | 2 |
| visual dorsal R | Ctx V3CD R | supplementary V3 R | Cortex_Visual_Central | 1456 | 37 | -81 | 15 | -12.72 | 0 | 4 | -10.22 | 0 | 6 | 0 | 4 |
| visual dorsal R | Ctx V6 R | V6 R | Cortex_Visual_Peripheral | 2416 | 18 | -77 | 30 | -2.23 | 2 | 3 | -2.9 | 1 | 3 | 0 | 2 |
| visual dorsal R | Ctx V6A R | V6 R | Cortex_Visual_Peripheral | 1032 | 24 | -82 | 46 | -4.03 | 0 | 3 | -2.7 | 0 | 4 | 0 | 2 |
| visual dorsal R | Ctx V7 R | V7 R | Cortex_Visual_Peripheral | 1320 | 29 | -82 | 34 | -4.11 | 0 | 4 | -3.9 | 0 | 4 | 0 | 3 |
| visual early L | Ctx V1 L | striate L | Cortex_Visual_Peripheral | 20216 | -10 | -85 | 2 | 4.91 | 3 | 1 | -7.25 | 1 | 5 | 1 | 1 |
| visual early L | Ctx V2 L | extrastriate L | Cortex_Visual_Peripheral | 15816 | -9 | -79 | 4 | -7.8 | 1 | 3 | -7.62 | 1 | 5 | 0 | 3 |
| visual early L | Ctx V3 L | extrastriate L | Cortex_Visual_Central | 11552 | -15 | -85 | 8 | -3.69 | 1 | 3 | -6.67 | 1 | 5 | 1 | 2 |
| visual early L | Ctx V4 L | extrastriate L | Cortex_Visual_Central | 9128 | -27 | -86 | -0 | -3.53 | 0 | 4 | -14.21 | 0 | 6 | 0 | 3 |
| visual early R | Ctx V1 R | striate R | Cortex_Visual_Peripheral | 21760 | 13 | -81 | 4 | 5.81 | 2 | 1 | -12.41 | 1 | 5 | 1 | 0 |
| visual early R | Ctx V2 R | extrastriate R | Cortex_Visual_Central | 15488 | 12 | -75 | 5 | -5.71 | 1 | 3 | -3.51 | 1 | 5 | 0 | 2 |
| visual early R | Ctx V3 R | extrastriate R | Cortex_Visual_Central | 12256 | 19 | -85 | 9 | -4.38 | 1 | 3 | -4.77 | 1 | 5 | 0 | 2 |
| visual early R | Ctx V4 R | extrastriate R | Cortex_Visual_Central | 8088 | 29 | -83 | -0 | -4.26 | 0 | 3 | -12.95 | 0 | 6 | 0 | 2 |
| visual ventral L | Ctx FFC L | FFC L | Cortex_Dorsal_AttentionA | 5616 | -42 | -61 | -19 | -7.09 | 0 | 4 | -6.17 | 1 | 5 | 0 | 4 |

| Structure | Region | Parcel | Network | Volume | MNI Coordinates | | | Experienced Pain (Hot-Warm) | | | Imagined Pain (Imagine-Warm) | | | Pain Conjunction | |
|---|---|---|---|---|---|---|---|---|---|---|---|---|---|---|---|
| | | | | | X | Y | Z | T | Activated (n) | Deactivated (n) | T | Activated (n) | Deactivated (n) | Activated (n) | Deactivated (n) |
| visual ventral L | Ctx PIT L | FFC L | Cortex_Visual_Central | 2096 | -38 | -84 | -15 | -4.15 | 1 | 3 | 4.9 | 2 | 4 | 0 | 1 |
| visual ventral L | Ctx V8 L | VVC L | Cortex_Visual_Central | 1904 | -31 | -75 | -12 | -3.13 | 0 | 2 | -5 | 1 | 6 | 0 | 1 |
| visual ventral L | Ctx VMV1 L | VMV L | Cortex_Visual_Peripheral | 1792 | -19 | -55 | -6 | -4.12 | 1 | 3 | -2.1 | 1 | 3 | 0 | 3 |
| visual ventral L | Ctx VMV2 L | VMV L | Cortex_Visual_Peripheral | 1664 | -29 | -55 | -4 | -6.92 | 1 | 2 | 10.71 | 1 | 3 | 0 | 2 |
| visual ventral L | Ctx VMV3 L | VMV L | Cortex_Visual_Peripheral | 1312 | -28 | -61 | -11 | 2.65 | 1 | 2 | -4.3 | 1 | 6 | 0 | 2 |
| visual ventral L | Ctx VVC L | VVC L | Cortex_Visual_Central | 3032 | -31 | -52 | -16 | -5.85 | 0 | 4 | -3.85 | 2 | 5 | 0 | 4 |
| visual ventral R | Ctx FFC R | FFC R | Cortex_Visual_Central | 6080 | 41 | -54 | -20 | -4.72 | 0 | 5 | -8.12 | 2 | 3 | 0 | 4 |
| visual ventral R | Ctx PIT R | FFC R | Cortex_Visual_Central | 3136 | 41 | -80 | -13 | -5.88 | 0 | 3 | -6.04 | 3 | 3 | 0 | 3 |
| visual ventral R | Ctx V8 R | VVC R | Cortex_Visual_Central | 2448 | 31 | -72 | -12 | -3.16 | 1 | 3 | -6.16 | 1 | 5 | 0 | 3 |
| visual ventral R | Ctx VMV1 R | VMV R | Cortex_Visual_Peripheral | 2320 | 19 | -55 | -7 | -4.36 | 1 | 2 | -6.46 | 1 | 3 | 0 | 1 |
| visual ventral R | Ctx VMV2 R | VMV R | Cortex_Visual_Peripheral | 1808 | 29 | -51 | -4 | -5.57 | 1 | 2 | 4.63 | 1 | 3 | 0 | 1 |
| visual ventral R | Ctx VMV3 R | VMV R | Cortex_Visual_Peripheral | 1864 | 29 | -59 | -8 | -6.99 | 1 | 3 | -3.35 | 1 | 4 | 0 | 3 |
| visual ventral R | Ctx VVC R | VVC R | Cortex_Visual_Central | 3792 | 29 | -45 | -18 | -7.49 | 0 | 6 | -4.97 | 1 | 5 | 0 | 3 |

**Notes.** Voxelwise group-level robust-regression contrasts for experienced pain (Hot–Warm) and imagined pain (Imagine–Warm), summarized by CANLab-2024 atlas parcel (N = 9 participants; FDR q < .05). For each parcel we list its Structure, Region, Parcel, and Network, its volume (mm³), and its MNI centroid (X, Y, Z). Within each contrast, T is the peak (maximum-magnitude) group-level voxelwise *t* within the parcel. Activated (n) and Deactivated (n) give the number of participants (of 9) with a significant positive or negative effect within the parcel, computed from each participant's FDR-thresholded voxelwise map: a participant is counted as *activated* when the parcel's significant voxels have a positive peak or a positive mean, and as *deactivated* when the parcel's significant voxels have both a negative peak and a negative mean. Pain Conjunction reports the number of participants with a significant effect of the same sign within the parcel for both contrasts (Activated = positive in both Hot–Warm and Imagine–Warm; Deactivated = negative in both). Parcels are grouped by anatomical structure. n: Number of subjects exhibiting a significant effect. † $p < .10$, * $p < 0.05$, ** $p < 0.01$ and *** $p < .001$.

### Supplementary Table S3: Pattern expression of participant-specific nociceptive body site selective SVM weight maps in a priori pain regions

| | MNI Coordinates | | | Hot-Rest (Target Body Site) | | Hot-Warm (Target Body Site) | | Hot-Warm x Target – Other Body Site | | Imagine-Rest (Target Body Site) | | Imagine-Warm (Target Body Site) | | Imagine-Warm x Target – Other Body Site | |
|---|---|---|---|---|---|---|---|---|---|---|---|---|---|---|---|
| Region | X | Y | Z | T | n | T | n | T | n | T | n | T | n | T | n |
| Amygdala L | -22.5 | -4.5 | -20.5 | 3.75** | 7 | 3.65** | 7 | 1.96† | 5 | -1.87† | 0 | 1.27 | 3 | 0.410 | 1 |
| Amygdala R | 23.5 | -2.5 | -20.5 | | | | | | | | | | | | |
| aMCC MPFC | 1.5 | 15.5 | 33.5 | 4.67** | 8 | 4.36** | 7 | 1.27 | 2 | -1.83 | 1 | 0.94 | 2 | -1.51 | 0 |
| aIns L | -40.5 | 23.5 | 3.5 | 7.45*** | 9 | 7.28*** | 9 | 0.79 | 3 | -2.10+ | 0 | 1.59 | 3 | -0.89 | 1 |
| aIns R | 39.5 | 25.5 | 3.5 | | | | | | | | | | | | |
| mIns L | -38.5 | 3.5 | 3.5 | 9.27*** | 9 | 9.38*** | 9 | 0.26 | 2 | -3.91** | 0 | 0.19 | 1 | -1.31 | 1 |
| mIns R | 37.5 | 3.5 | 3.5 | | | | | | | | | | | | |
| dpIns L | -40.5 | -22.5 | 19.5 | 10.74*** | 9 | 1.00*** | 9 | 5.36*** | 8 | -6.80*** | 0 | -0.11 | 2 | -0.55 | 0 |
| dpIns R | 39.5 | -18.5 | 17.5 | | | | | | | | | | | | |
| S1 L | -34.5 | -28.5 | 55.5 | 6.76*** | 9 | 6.81*** | 9 | 4.44** | 8 | -2.63* | 0 | 1.01 | 4 | -0.41 | 2 |
| S1 R | 35.5 | -24.5 | 55.5 | | | | | | | | | | | | |
| S2 L | 55.5 | -16.5 | 19.5 | 10.53*** | 9 | 10.22*** | 9 | 2.63** | 6 | -4.46** | 0 | -0.72 | 1 | -1.40 | 0 |
| S2 R | -56.5 | -22.5 | 19.5 | | | | | | | | | | | | |
| Hypothalamus | -0.5 | -2.5 | -12.5 | 2.08+ | 3 | 2.12† | 4 | 0.81 | 2 | -0.43 | 0 | 0.66 | 1 | 0.91 | 2 |
| Thal VPLM L | 17.5 | -18.5 | 5.5 | 2.75* | 5 | 2.75* | 5 | 1.05 | 3 | -1.67 | 0 | 1.23 | 2 | -0.72 | 0 |
| Thal VPLM R | -18.5 | -20.5 | 5.5 | | | | | | | | | | | | |
| Thal IL | -0.5 | -12.5 | -0.5 | 4.33** | 8 | 4.66** | 8 | 2.12+ | 4 | -2.08* | 0 | 1.46 | 3 | -0.76 | 0 |
| Thal MD | -0.5 | -14.5 | 5.5 | 3.92** | 7 | 3.84** | 7 | 1.49 | 2 | -2.77** | 0 | 0.94 | 2 | -2.10* | 0 |
| PAG | -0.5 | -32.5 | -10.5 | 2.14* | 4 | 2.43* | 6 | 0.82 | 3 | 0.06 | 0 | 1.29 | 0 | 0.73 | 1 |
| Parabrachial nuclei L | -6.5 | -36.5 | -22.5 | 1.85+ | 3 | 2.10† | 3 | 2.41* | 7 | -1.00 | 0 | 0.51 | 2 | 0.50 | 3 |
| Parabrachial nuclei R | 7.5 | -38.5 | -22.5 | | | | | | | | | | | | |
| RVM | 3.5 | -34.5 | -42.5 | 0.10 | 1 | -0.06 | 1 | -2.05† | 0 | -0.16 | 2 | -0.25 | 1 | -0.70 | 0 |

**Notes.** n: Number of subjects with significant pattern expression. aMCC MPFC: anterior midcingulate cortex/medial prefrontal cortex, aIns: anterior insula, mIns: medial insula, dpIns: dorsoposterior insula, S1: primary somatosensory cortex, S2: secondary somatosensory cortex, Thal: thalamus, VPLM: medial ventral posterolateral, IL: intralaminar, MD: mediodorsal, PAG: periaqueductal gray, RVM: rostral ventral medulla. † $p$ < .10, * $p$ < 0.05, ** $p$ < 0.01 and *** $p$ < .001.

Supplementary Table S4: Pattern expression of participant-specific nociceptive body site selective SVM weight maps in 131 CANlab 2024 atlas parcels

| Parcel | Region | Pain Region | MNI Coordinates | | | Hot-Rest (Target Body Site) | | Hot-Warm (Target Body Site) | | Hot-Warm x Target – Other Body Site | | Imagine-Rest (Target Body Site) | | Imagine-Warm (Target Body Site) | | Imagine-Warm x Target – Other Body Site | |
|---|---|---|---|---|---|---|---|---|---|---|---|---|---|---|---|---|---|
| | | | X | Y | Z | T | n | T | n | T | n | T | n | T | n | T | n |
| 55b L | somatomotor premotor L | | -48.5 | -0.5 | 49.5 | 4.51** | 8 | 4.69** | 8 | 2.18 | 4 | -0.46 | 2 | 1.96 | 4 | -1.12 | 0 |
| 55b R | somatomotor premotor R | | 51.5 | 3.5 | 47.5 | | | | | | | | | | | | |
| aCAU L | CAU L | | -12.5 | 15.5 | 5.5 | 3.17* | 7 | 3.13* | 7 | 2.19 | 3 | -0.46 | 0 | 1.43 | 1 | 0.21 | 0 |
| aCAU R | CAU R | | 13.5 | 17.5 | 5.5 | | | | | | | | | | | | |
| AMY BL CeM L | Amygdala L | Amygdala | -20.5 | -4.5 | -18.5 | 3.85* | 7 | 3.84* | 7 | 2.37 | 5 | -1.7 | 0 | 1.65 | 2 | 0.86 | 0 |
| AMY BL CeM R | Amygdala R | Amygdala | 21.5 | -4.5 | -18.5 | | | | | | | | | | | | |
| AMY La L | Amygdala L | Amygdala | -28.5 | -2.5 | -22.5 | 2.14† | 4 | 2.16† | 5 | 1.5 | 0 | -1.19 | 0 | -0.01 | 0 | -0.62 | 0 |
| AMY La R | Amygdala R | Amygdala | 27.5 | -2.5 | -24.5 | | | | | | | | | | | | |
| anterior AAC L | auditory association cortex L | | -52.5 | -8.5 | -12.5 | 2.98* | 4 | 2.95* | 4 | 1.43 | 2 | -0.14 | 1 | 2.54 | 3 | 0.39 | 0 |
| anterior AAC R | auditory association cortex R | | 53.5 | -4.5 | -14.5 | | | | | | | | | | | | |
| anterior agranular insula L | insula anterior L | aIns/mIns | -32.5 | 17.5 | -2.5 | 7.9*** | 9 | 7.96*** | 9 | 0.28 | 1 | -2.78 | 0 | 0.65 | 2 | -1.06 | 0 |
| anterior agranular insula R | insula anterior R | aIns/mIns | 33.5 | 19.5 | -4.5 | | | | | | | | | | | | |
| anterior BA7 L | parietal superior lobule L | | -12.5 | -54.5 | 65.5 | 5.16** | 9 | 5.11** | 9 | 4.45* | 8 | -3.6† | 0 | 0.72 | 0 | 0.6 | 0 |
| anterior BA7 R | parietal superior lobule R | | 15.5 | -54.5 | 67.5 | | | | | | | | | | | | |
| anterior IPL L | parietal inferior lobule L | | -58.5 | -32.5 | 35.5 | 7.94*** | 9 | 8.39*** | 9 | 1.37 | 4 | -3.13 | 0 | 0.33 | 2 | -1.12 | 0 |
| anterior IPL R | parietal inferior lobule R | | 59.5 | -24.5 | 35.5 | | | | | | | | | | | | |
| anterior operculum L | insula operculum L | aIns | -38.5 | 19.5 | 7.5 | 9.45*** | 9 | 9.51*** | 9 | -0.05 | 1 | -2.38 | 0 | 1.3 | 2 | -1.31 | 0 |
| anterior operculum R | insula operculum R | aIns | 37.5 | 21.5 | 7.5 | | | | | | | | | | | | |
| anterior precuneus L | cingulate posterior L | | -6.5 | -30.5 | 41.5 | 3.73* | 6 | 3.71* | 6 | 1.28 | 2 | -3.97* | 0 | 0.22 | 2 | -1.44 | 0 |
| anterior precuneus R | cingulate posterior R | | 9.5 | -32.5 | 41.5 | | | | | | | | | | | | |
| aPUT L | PUT L | | -22.5 | 9.5 | -4.5 | 3.35* | 6 | 3.2* | 5 | 1.78 | 3 | -0.61 | 1 | 1.13 | 1 | 0.12 | 1 |
| aPUT R | PUT R | | 23.5 | 9.5 | -4.5 | | | | | | | | | | | | |
| auditory transition area L | auditory association cortex L | | -58.5 | -8.5 | -0.5 | 9.07*** | 9 | 8.9*** | 9 | 0.99 | 1 | -1.86 | 1 | 1.58 | 3 | -1.03 | 0 |
| auditory transition area R | auditory association cortex R | | 59.5 | -4.5 | -2.5 | | | | | | | | | | | | |
| auditory+ L | auditory early L | dpIns | -44.5 | -28.5 | 9.5 | 9.64*** | 9 | 9.4*** | 9 | 3.07† | 6 | -5.65* | 0 | -0.22 | 1 | -0.63 | 0 |
| auditory+ R | auditory early R | dpIns | 47.5 | -22.5 | 11.5 | | | | | | | | | | | | |
| BA10 L | cingulate ventral frontal L | | -14.5 | 57.5 | -2.5 | 3.48* | 7 | 3.4* | 7 | 0.78 | 2 | -0.94 | 1 | 1.78 | 2 | 0.06 | 0 |

| Parcel | Region | Pain Region | MNI Coordinates | | | Hot-Rest (Target Body Site) | | Hot-Warm (Target Body Site) | | Hot-Warm x Target – Other Body Site | | Imagine-Rest (Target Body Site) | | Imagine-Warm (Target Body Site) | | Imagine-Warm x Target – Other Body Site | |
|---|---|---|---|---|---|---|---|---|---|---|---|---|---|---|---|---|---|
| | | | X | Y | Z | T | n | T | n | T | n | T | n | T | n | T | n |
| BA10 R | cingulate ventral frontal R | | 13.5 | 59.5 | -4.5 | | | | | | | | | | | | |
| BA11 L | cingulate ventral frontal L | | -26.5 | 47.5 | -12.5 | 5.48** | 9 | 5.83** | 9 | 2.45 | 5 | -1.14 | 0 | 1.98 | 1 | 0.41 | 0 |
| BA11 R | cingulate ventral frontal R | | 25.5 | 47.5 | -14.5 | | | | | | | | | | | | |
| BA13 L | cingulate ventral frontal L | | -22.5 | 29.5 | -18.5 | 4.28** | 8 | 4.48** | 8 | 2.08 | 3 | -2.25 | 0 | 0.35 | 1 | -0.03 | 0 |
| BA13 R | cingulate ventral frontal R | | 21.5 | 29.5 | -16.5 | | | | | | | | | | | | |
| BA44 L | cingulate vlPFC L | | -52.5 | 15.5 | 13.5 | 5.84** | 9 | 6.08** | 9 | 0.15 | 3 | -1.05 | 0 | 1.96 | 4 | -0.99 | 0 |
| BA44 R | cingulate vlPFC R | | 53.5 | 19.5 | 11.5 | | | | | | | | | | | | |
| BA45 L | cingulate vlPFC L | aIns | -50.5 | 25.5 | 3.5 | 5.13** | 8 | 5.1** | 8 | 0.8 | 2 | -1.41 | 0 | 1.44 | 3 | -0.78 | 0 |
| BA45 R | cingulate vlPFC R | aIns | 51.5 | 27.5 | 1.5 | | | | | | | | | | | | |
| BA46 L | cingulate dlPFC L | | -36.5 | 39.5 | 23.5 | 4.43** | 7 | 4.7** | 7 | 0.79 | 2 | -1.47 | 0 | 1.54 | 4 | -0.73 | 0 |
| BA46 R | cingulate dlPFC R | | 37.5 | 41.5 | 25.5 | | | | | | | | | | | | |
| BA47 L | cingulate ventral frontal L | | -40.5 | 35.5 | -10.5 | 4.16* | 7 | 4.35** | 8 | 1.85 | 4 | -2.22 | 0 | 1.98 | 3 | -0.36 | 0 |
| BA47 R | cingulate ventral frontal R | | 39.5 | 37.5 | -10.5 | | | | | | | | | | | | |
| BA5 L | somatomotor paracentral lobule L | S1 | -12.5 | -40.5 | 63.5 | 6.39** | 9 | 6.6** | 9 | 6.32*** | 9 | -4.23* | 0 | 0.09 | 1 | 0.45 | 0 |
| BA5 R | somatomotor paracentral lobule R | S1 | 11.5 | -40.5 | 65.5 | | | | | | | | | | | | |
| BA9 L | cingulate dlPFC L | | -20.5 | 49.5 | 33.5 | 3.33* | 6 | 3.41* | 6 | 1.83 | 4 | -0.02 | 1 | 1.86 | 4 | 0.22 | 1 |
| BA9 R | cingulate dlPFC R | | 19.5 | 55.5 | 29.5 | | | | | | | | | | | | |
| Cblm CrusI CrusII VIIIb L | Cblm cortex L | | -30.5 | -70.5 | -38.5 | 3.79* | 5 | 3.57* | 5 | 1.16 | 1 | -0.67 | 0 | 2.18 | 3 | -0.23 | 0 |
| Cblm CrusI CrusII VIIIb R | Cblm cortex R | | 31.5 | -70.5 | -38.5 | | | | | | | | | | | | |
| Cblm I IV V VI L | Cblm cortex L | | -18.5 | -54.5 | -22.5 | 3.84* | 5 | 3.75* | 5 | 3.47† | 5 | -0.58 | 2 | 2.5 | 4 | -0.72 | 0 |
| Cblm I IV V VI R | Cblm cortex R | | 19.5 | -54.5 | -22.5 | | | | | | | | | | | | |
| Cblm vermis | Cblm vermis | | -0.5 | -66.5 | -30.5 | 2.73* | 5 | 2.8* | 5 | 1.09 | 0 | -1.49 | 0 | 1.68 | 2 | -0.48 | 0 |
| Cblm VIIIa VIIb IX X L | Cblm cortex L | | -18.5 | -54.5 | -50.5 | 5.15** | 8 | 5.2** | 8 | 2.15 | 4 | -1.75 | 0 | 2.19 | 4 | 0.45 | 0 |
| Cblm VIIIa VIIb IX X R | Cblm cortex R | | 17.5 | -54.5 | -52.5 | | | | | | | | | | | | |
| Cholinergic nuclei L | Pons L | | -2.5 | -38.5 | -28.5 | 1.54 | 2 | 1.23 | 2 | 0.55 | 0 | 0.11 | 0 | 0.36 | 1 | 0.26 | 0 |
| Cholinergic nuclei R | Pons R | | 3.5 | -36.5 | -26.5 | | | | | | | | | | | | |
| cingulate motor area L | somatomotor paracentral lobule L | | -8.5 | -12.5 | 49.5 | 6.69*** | 9 | 6.17** | 9 | 3.57† | 5 | -2.9 | 0 | 0.63 | 2 | -0.66 | 0 |
| cingulate motor area R | somatomotor paracentral lobule R | | 9.5 | -12.5 | 51.5 | | | | | | | | | | | | |
| Cranial nuclei L | Medulla L | | -6.5 | -42.5 | -42.5 | 1.58 | 2 | 1.6 | 3 | 0.64 | 0 | -0.59 | 0 | 0.36 | 0 | -0.36 | 0 |
| Cranial nuclei R | Medulla R | | 5.5 | -42.5 | -44.5 | | | | | | | | | | | | |

| Parcel | Region | Pain Region | MNI Coordinates X | Y | Z | Hot-Rest (Target Body Site) T | n | Hot-Warm (Target Body Site) T | n | Hot-Warm x Target – Other Body Site T | n | Imagine-Rest (Target Body Site) T | n | Imagine-Warm (Target Body Site) T | n | Imagine-Warm x Target – Other Body Site T | n |
|---|---|---|---|---|---|---|---|---|---|---|---|---|---|---|---|---|---|
| dmPFC L | cingulate ACC mPFC L | | -6.5 | 43.5 | 31.5 | 2.81* | 6 | 2.74* | 6 | 1.03 | 3 | -1.69 | 0 | 0.92 | 3 | -0.19 | 0 |
| dmPFC R | cingulate ACC mPFC R | | 7.5 | 43.5 | 31.5 | | | | | | | | | | | | |
| dMT+ L | visual MT+ L | | -40.5 | -68.5 | 7.5 | 3.06* | 5 | 3.04* | 5 | 1.13 | 2 | -0.79 | 0 | 1.92 | 2 | 0.25 | 0 |
| dMT+ R | visual MT+ R | | 45.5 | -66.5 | 7.5 | | | | | | | | | | | | |
| dorsal intraparietal L | parietal superior lobule L | | -26.5 | -60.5 | 59.5 | 5.14** | 9 | 5.12** | 9 | 4.16** | 9 | -2.1 | 0 | 1.48 | 2 | 0.59 | 0 |
| dorsal intraparietal R | parietal superior lobule R | | 25.5 | -58.5 | 59.5 | | | | | | | | | | | | |
| early auditory ctx a1 L | auditory early L | | -42.5 | -24.5 | 11.5 | 9.08*** | 9 | 8.51*** | 9 | 3.83* | 8 | -4.86* | 0 | -0.13 | 1 | 1.28 | 0 |
| early auditory ctx a1 R | auditory early R | | 43.5 | -20.5 | 11.5 | | | | | | | | | | | | |
| extrastriate L | visual early L | | -14.5 | -80.5 | 1.5 | 3.16* | 5 | 3.23* | 5 | 1.74 | 2 | -0.57 | 1 | 1.98 | 3 | -0.36 | 0 |
| extrastriate R | visual early R | | 17.5 | -80.5 | 1.5 | | | | | | | | | | | | |
| FFC L | visual ventral L | | -40.5 | -68.5 | -18.5 | 3.24* | 6 | 2.98* | 5 | 1.25 | 2 | -0.9 | 0 | 1.82 | 2 | -0.41 | 0 |
| FFC R | visual ventral R | | 41.5 | -62.5 | -18.5 | | | | | | | | | | | | |
| FOP1-BA43 L | somatomotor operculum L | | -54.5 | 1.5 | 9.5 | 10.46*** | 9 | 10.43*** | 9 | 1.28 | 3 | -2.76 | 0 | 0.64 | 2 | -1.13 | 0 |
| FOP1-BA43 R | somatomotor operculum R | | 53.5 | 1.5 | 9.5 | | | | | | | | | | | | |
| GPe L | GP L | | -20.5 | -2.5 | -2.5 | 0.86 | 3 | 0.61 | 3 | -1.4 | 0 | -0.48 | 0 | -0.15 | 0 | -0.98 | 0 |
| GPe R | GP R | | 19.5 | -0.5 | -0.5 | | | | | | | | | | | | |
| GPi L | GP L | | -18.5 | -6.5 | -4.5 | 0.28 | 2 | 0.19 | 2 | -1.05 | 0 | 0.11 | 0 | 0.09 | 1 | -0.34 | 0 |
| GPi R | GP R | | 19.5 | -4.5 | -4.5 | | | | | | | | | | | | |
| Hipp L | Hippocampal Formation L | | -26.5 | -22.5 | -14.5 | 2.75* | 3 | 2.74* | 4 | 2.11 | 3 | -2.33 | 0 | 0.48 | 0 | -0.02 | 0 |
| Hipp R | Hippocampal Formation R | | 27.5 | -20.5 | -14.5 | | | | | | | | | | | | |
| hippocampal formation L | temporal medial L | | -18.5 | -20.5 | -22.5 | 4.09** | 7 | 4.63** | 8 | 2.01 | 4 | -1.8 | 0 | 1.65 | 0 | -0.18 | 0 |
| hippocampal formation R | temporal medial R | | 19.5 | -20.5 | -22.5 | | | | | | | | | | | | |
| hypothalamus L | hypothalamus L | hypothalamus | -2.5 | -2.5 | -12.5 | 2.08† | 3 | 2.12† | 4 | 0.81 | 1 | -0.43 | 0 | 0.66 | 0 | 0.91 | 0 |
| hypothalamus R | hypothalamus R | hypothalamus | 3.5 | -2.5 | -12.5 | | | | | | | | | | | | |
| IFJ L | cingulate vlPFC L | | -38.5 | 7.5 | 27.5 | 3.59* | 6 | 3.75* | 6 | 1.19 | 2 | -0.94 | 0 | 1.45 | 4 | -1.01 | 0 |
| IFJ R | cingulate vlPFC R | | 41.5 | 13.5 | 25.5 | | | | | | | | | | | | |
| IFS L | cingulate vlPFC L | | -48.5 | 27.5 | 15.5 | 4.37** | 8 | 4.38** | 7 | 0.39 | 2 | -1.69 | 0 | 1.41 | 4 | -1.34 | 0 |
| IFS R | cingulate vlPFC R | | 47.5 | 33.5 | 11.5 | | | | | | | | | | | | |
| inferior dysgranular insula L | insula posterior L | mIns | -40.5 | -4.5 | -4.5 | 6.93*** | 9 | 6.91*** | 9 | 0.41 | 1 | -4.56* | 0 | -0.84 | 1 | -1.62 | 0 |
| inferior dysgranular insula R | insula posterior R | mIns | 41.5 | -2.5 | -4.5 | | | | | | | | | | | | |

| | | | MNI Coordinates | | | Hot-Rest (Target Body Site) | | Hot-Warm (Target Body Site) | | Hot-Warm x Target – Other Body Site | | Imagine-Rest (Target Body Site) | | Imagine-Warm (Target Body Site) | | Imagine-Warm x Target – Other Body Site | |
|---|---|---|---|---|---|---|---|---|---|---|---|---|---|---|---|---|---|
| Parcel | Region | Pain Region | X | Y | Z | T | n | T | n | T | n | T | n | T | n | T | n |
| inferior IPL L | parietal inferior lobule L | | -42.5 | -64.5 | 29.5 | 2.41† | 5 | 2.59* | 5 | 1.25 | 2 | -2.02 | 0 | 0.94 | 2 | -0.42 | 0 |
| inferior IPL R | parietal inferior lobule R | | 47.5 | -60.5 | 31.5 | | | | | | | | | | | | |
| Inferior lateral BA8 L | cingulate dlPFC L | | -38.5 | 13.5 | 47.5 | 3.15* | 5 | 3.3* | 5 | 2.61 | 5 | -2.58 | 0 | 0.86 | 2 | -0.91 | 0 |
| Inferior lateral BA8 R | cingulate dlPFC R | | 39.5 | 15.5 | 45.5 | | | | | | | | | | | | |
| inferior premotor L | somatomotor premotor L | | -52.5 | 3.5 | 25.5 | 6.74*** | 9 | 7.08*** | 9 | 0.58 | 2 | -0.85 | 0 | 1.88 | 4 | -1.97 | 0 |
| inferior premotor R | somatomotor premotor R | | 53.5 | 7.5 | 23.5 | | | | | | | | | | | | |
| inferior temporal sulcus L | temporal lateral L | | -48.5 | -28.5 | -24.5 | 2.56* | 5 | 2.41† | 4 | 1.22 | 2 | -1.31 | 0 | 1.45 | 2 | -0.58 | 0 |
| inferior temporal sulcus R | temporal lateral R | | 49.5 | -22.5 | -26.5 | | | | | | | | | | | | |
| infralimbic L | cingulate ACC mPFC L | | -4.5 | 23.5 | -10.5 | 1.73 | 3 | 1.57 | 3 | 0.04 | 1 | -0.89 | 0 | 0.21 | 0 | -0.5 | 0 |
| infralimbic R | cingulate ACC mPFC R | | 3.5 | 23.5 | -10.5 | | | | | | | | | | | | |
| ISP1 L | visual dorsal L | | -24.5 | -72.5 | 35.5 | 2.8* | 5 | 2.92* | 5 | 1.5 | 2 | -1.45 | 1 | 0.6 | 3 | -0.08 | 0 |
| ISP1 R | visual dorsal R | | 27.5 | -68.5 | 37.5 | | | | | | | | | | | | |
| PnO PnC L | Pons L | RVM | -2.5 | -34.5 | -32.5 | -0.11 | 0 | -0.36 | 0 | -0.97 | 0 | -0.25 | 0 | -0.48 | 0 | -0.51 | 0 |
| PnO PnC R | Pons R | RVM | 3.5 | -34.5 | -32.5 | | | | | | | | | | | | |
| LC+ L | Pons L | | -6.5 | -36.5 | -34.5 | 0.39 | 2 | 0.16 | 1 | -0.13 | 1 | -0.67 | 0 | -0.46 | 0 | -0.31 | 0 |
| LC+ R | Pons R | RVM | 7.5 | -36.5 | -34.5 | | | | | | | | | | | | |
| LO L | visual MT+ L | | -40.5 | -82.5 | 5.5 | 2.62* | 5 | 2.61* | 5 | 2.33 | 4 | 0.06 | 1 | 1.92 | 3 | 0.25 | 0 |
| LO R | visual MT+ R | | 41.5 | -78.5 | 5.5 | | | | | | | | | | | | |
| M1 L | somatomotor primary L | | -26.5 | -20.5 | 55.5 | 6.39** | 9 | 6.51** | 9 | 4.91* | 9 | -1.92 | 0 | 1.26 | 3 | 0.33 | 0 |
| M1 R | somatomotor primary R | | 25.5 | -18.5 | 57.5 | | | | | | | | | | | | |
| Medullary Raphe (Serotonergic) | Medulla | RVM | -0.5 | -36.5 | -52.5 | -0.31 | 0 | -0.17 | 1 | -0.52 | 0 | 1.02 | 0 | 0.68 | 1 | 0.31 | 0 |
| Medullary reticular formation L | Medulla L | | -6.5 | -42.5 | -54.5 | 1.8 | 4 | 2.23† | 6 | 0.57 | 0 | -0.53 | 0 | 0.66 | 0 | 0.67 | 0 |
| Medullary reticular formation R | Medulla R | RVM | 5.5 | -42.5 | -54.5 | | | | | | | | | | | | |
| midcingulate L | cingulate ACC mPFC L | aMCC MPFC | -6.5 | 19.5 | 29.5 | 4.48** | 7 | 4.23** | 7 | 1.63 | 2 | -2.12 | 1 | 0.94 | 2 | -1.14 | 0 |
| midcingulate R | cingulate ACC mPFC R | aMCC MPFC | 7.5 | 17.5 | 31.5 | | | | | | | | | | | | |
| middle IPL L | parietal inferior lobule L | | -48.5 | -56.5 | 43.5 | 3.63* | 5 | 3.94* | 5 | 1.82 | 1 | -2.55 | 0 | 0.66 | 3 | -1.02 | 0 |
| middle IPL R | parietal inferior lobule R | | 53.5 | -46.5 | 45.5 | | | | | | | | | | | | |

| | | | MNI Coordinates | | | Hot-Rest (Target Body Site) | | Hot-Warm (Target Body Site) | | Hot-Warm x Target – Other Body Site | | Imagine-Rest (Target Body Site) | | Imagine-Warm (Target Body Site) | | Imagine-Warm x Target – Other Body Site | |
|---|---|---|---|---|---|---|---|---|---|---|---|---|---|---|---|---|---|
| Parcel | Region | Pain Region | X | Y | Z | T | n | T | n | T | n | T | n | T | n | T | n |
| middle temporal gyrus L | temporal lateral L | | -60.5 | -44.5 | -8.5 | 3.92* | 6 | 3.81* | 6 | 1.76 | 3 | -2.76 | 0 | 0.6 | 1 | -0.09 | 0 |
| middle temporal gyrus R | temporal lateral R | | 61.5 | -40.5 | -10.5 | | | | | | | | | | | | |
| MIP L | parietal superior lobule L | | -22.5 | -64.5 | 47.5 | 3.83* | 7 | 4.44** | 7 | 2.65 | 4 | -0.89 | 0 | 2.34 | 3 | 1.86 | 0 |
| MIP R | parietal superior lobule R | | 25.5 | -64.5 | 49.5 | | | | | | | | | | | | |
| OFC L | cingulate ventral frontal L | | -12.5 | 23.5 | -20.5 | 3.68* | 6 | 3.84* | 6 | 2.44 | 4 | -2.42 | 0 | 1.3 | 3 | 0.53 | 0 |
| OFC R | cingulate ventral frontal R | | 11.5 | 25.5 | -20.5 | | | | | | | | | | | | |
| Olivary complex L | Medulla L | | -6.5 | -36.5 | -52.5 | 1.34 | 2 | 1.42 | 2 | 0.44 | 2 | -1.63 | 0 | 0.11 | 1 | 0.27 | 0 |
| Olivary complex R | Medulla R | RVM | 5.5 | -36.5 | -52.5 | | | | | | | | | | | | |
| OP4 L | somatomotor operculum L | S2 | -58.5 | -12.5 | 15.5 | 10.77*** | 9 | 11.57*** | 9 | 1.97 | 5 | -3.94† | 0 | -0.93 | 1 | -1.19 | 0 |
| OP4 R | somatomotor operculum R | S2 | 57.5 | -10.5 | 15.5 | | | | | | | | | | | | |
| PAG | Midbrain | PAG | -0.5 | -32.5 | -10.5 | 2.14† | 4 | 2.43* | 5 | 0.82 | 2 | 0.06 | 0 | 1.29 | 0 | 0.73 | 0 |
| Parabrachial nuclei L | Pons L | Parabrachial nuclei | -6.5 | -36.5 | -22.5 | 1.67 | 3 | 1.85 | 3 | 1.83 | 2 | -0.57 | 0 | 0.75 | 2 | 0.38 | 0 |
| Parabrachial nuclei R | Pons R | Parabrachial nuclei | 7.5 | -38.5 | -22.5 | | | | | | | | | | | | |
| parietal occipital sulcus L | cingulate posterior L | | -14.5 | -64.5 | 25.5 | 3.71* | 5 | 3.79* | 5 | 1.28 | 3 | -1.57 | 0 | 1.54 | 2 | -0.6 | 0 |
| parietal occipital sulcus R | cingulate posterior R | | 17.5 | -62.5 | 27.5 | | | | | | | | | | | | |
| pCAU L | CAU L | | -14.5 | -2.5 | 19.5 | 2.63* | 4 | 2.64* | 4 | 1.45 | 1 | -0.82 | 0 | 1.31 | 1 | -0.5 | 0 |
| pCAU R | CAU R | | 13.5 | -0.5 | 19.5 | | | | | | | | | | | | |
| PCV L | cingulate posterior L | | -6.5 | -50.5 | 51.5 | 4.08** | 7 | 4.42** | 8 | 2.21 | 3 | -1.32 | 0 | 1.49 | 3 | 0.12 | 0 |
| PCV R | cingulate posterior R | | 7.5 | -52.5 | 51.5 | | | | | | | | | | | | |
| PeEc L | temporal medial L | | -30.5 | -8.5 | -32.5 | 4.02** | 8 | 4** | 7 | 1.33 | 3 | -1.42 | 0 | 0.83 | 1 | 0.45 | 0 |
| PeEc R | temporal medial R | | 27.5 | -6.5 | -32.5 | | | | | | | | | | | | |
| PH RPH L | visual MT+ L | | -44.5 | -64.5 | -6.5 | 3.3* | 5 | 3.03* | 5 | 2.1 | 5 | -1.66 | 0 | 1.13 | 2 | 0.39 | 0 |
| PH RPH R | visual MT+ R | | 47.5 | -62.5 | -10.5 | | | | | | | | | | | | |
| PHA L | temporal medial L | | -28.5 | -36.5 | -14.5 | 3.46* | 5 | 3.45* | 5 | 0.91 | 1 | -0.03 | 0 | 2.29 | 4 | -0.32 | 0 |
| PHA R | temporal medial R | | 27.5 | -34.5 | -14.5 | | | | | | | | | | | | |
| PIR L | insula anterior L | | -30.5 | 3.5 | -16.5 | 7.04*** | 9 | 7.64*** | 9 | 1.74 | 3 | -3.34† | 0 | -0.34 | 1 | -1.39 | 0 |
| PIR R | insula anterior R | | 31.5 | 5.5 | -18.5 | | | | | | | | | | | | |
| postcentral BA7 L | parietal superior lobule L | | -36.5 | -48.5 | 63.5 | 6.91*** | 9 | 6.91*** | 9 | 3.93* | 8 | -2.93 | 0 | 1.29 | 2 | -0.43 | 0 |
| postcentral BA7 R | parietal superior lobule R | | 35.5 | -46.5 | 63.5 | | | | | | | | | | | | |

| Parcel | Region | Pain Region | MNI Coordinates | | | Hot-Rest (Target Body Site) | | Hot-Warm (Target Body Site) | | Hot-Warm x Target – Other Body Site | | Imagine-Rest (Target Body Site) | | Imagine-Warm (Target Body Site) | | Imagine-Warm x Target – Other Body Site | |
|---|---|---|---|---|---|---|---|---|---|---|---|---|---|---|---|---|---|
| | | | X | Y | Z | T | n | T | n | T | n | T | n | T | n | T | n |
| posterior AAC L | auditory association cortex L | | -52.5 | -32.5 | -2.5 | 3.09* | 5 | 3.24* | 6 | 1.63 | 2 | -1.61 | 0 | 2.21 | 3 | 0.38 | 0 |
| posterior AAC R | auditory association cortex R | | 53.5 | -28.5 | -2.5 | | | | | | | | | | | | |
| posterior BA7 L | parietal superior lobule L | | -8.5 | -70.5 | 55.5 | 4.23** | 7 | 4.27** | 6 | 2.72 | 4 | -1.41 | 0 | 1.88 | 3 | 0.5 | 0 |
| posterior BA7 R | parietal superior lobule R | | 11.5 | -68.5 | 57.5 | | | | | | | | | | | | |
| posterior granular insula L | insula posterior L | dpIns | -36.5 | -16.5 | 9.5 | 11.07*** | 9 | 10.95*** | 9 | 4.48* | 8 | -6.88* | 0 | -0.76 | 0 | 0.37 | 0 |
| posterior granular insula R | insula posterior R | dpIns | 37.5 | -14.5 | 9.5 | | | | | | | | | | | | |
| posterior IPL L | parietal inferior lobule L | | -34.5 | -80.5 | 25.5 | 3.37* | 5 | 3.1* | 5 | 2.04 | 3 | -1.37 | 0 | 0.86 | 2 | -0.08 | 0 |
| posterior IPL R | parietal inferior lobule R | | 39.5 | -76.5 | 27.5 | | | | | | | | | | | | |
| posterior operculum L | insula operculum L | mIns | -38.5 | -0.5 | 13.5 | 11.85*** | 9 | 11.53*** | 9 | 1.44 | 2 | -3.51† | 0 | 0.56 | 1 | -0.19 | 0 |
| posterior operculum R | insula operculum R | mIns | 37.5 | 3.5 | 15.5 | | | | | | | | | | | | |
| pPUT L | PUT L | | -28.5 | -6.5 | 3.5 | 3.22* | 6 | 3.06* | 6 | 0.71 | 2 | -1.37 | 1 | 0.5 | 1 | -0.98 | 0 |
| pPUT R | PUT R | | 29.5 | -6.5 | 1.5 | | | | | | | | | | | | |
| precuneus proper L | cingulate posterior L | | -6.5 | -52.5 | 31.5 | 2.58* | 5 | 2.69* | 5 | -0.08 | 1 | -1.85 | 0 | 0.62 | 1 | -0.84 | 0 |
| precuneus proper R | cingulate posterior R | | 7.5 | -52.5 | 33.5 | | | | | | | | | | | | |
| PSL L | parietal TPOJ L | | -56.5 | -44.5 | 23.5 | 6.73*** | 8 | 6.26** | 8 | 1.58 | 2 | -3.35† | 0 | 1.08 | 3 | -0.4 | 0 |
| PSL R | parietal TPOJ R | | 63.5 | -34.5 | 25.5 | | | | | | | | | | | | |
| RN L | Midbrain L | | -4.5 | -18.5 | -8.5 | 0.47 | 0 | 0.1 | 0 | 0.37 | 0 | -0.88 | 0 | -0.76 | 0 | -0.79 | 0 |
| RN R | Midbrain R | | 5.5 | -18.5 | -8.5 | | | | | | | | | | | | |
| Rostral Raphe (Serotonergic) | Midbrain | | -0.5 | -28.5 | -12.5 | 1.23 | 1 | 1.39 | 2 | 0.91 | 0 | 0.23 | 0 | 0.74 | 0 | 0.58 | 0 |
| Rostral reticular formation L | Midbrain L | | -8.5 | -26.5 | -4.5 | 2.44* | 5 | 2.55* | 4 | 1.39 | 3 | -1.03 | 0 | 0.55 | 1 | 0.45 | 0 |
| Rostral reticular formation R | Midbrain R | | 7.5 | -24.5 | -4.5 | | | | | | | | | | | | |
| RSC L | cingulate posterior L | | -4.5 | -42.5 | 19.5 | 3.23* | 6 | 3.32* | 6 | 0.92 | 3 | -1.85 | 0 | 1.23 | 1 | -0.99 | 0 |
| RSC R | cingulate posterior R | | 5.5 | -40.5 | 21.5 | | | | | | | | | | | | |
| S1 L | somatomotor primary L | S1 | -38.5 | -26.5 | 53.5 | 6.72*** | 9 | 6.78** | 9 | 4.21* | 7 | -2.45 | 0 | 1.03 | 2 | -0.47 | 0 |
| S1 R | somatomotor primary R | S1 | 37.5 | -24.5 | 53.5 | | | | | | | | | | | | |
| Shen Med L | Medulla L | | -2.5 | -38.5 | -60.5 | 2.54* | 5 | 2.91* | 6 | 0.56 | 0 | -0.54 | 1 | 0.76 | 1 | 0.22 | 0 |
| Shen Med R | Medulla R | RVM | 7.5 | -40.5 | -60.5 | | | | | | | | | | | | |

| Parcel | Region | Pain Region | MNI Coordinates X | Y | Z | Hot-Rest (Target Body Site) T | n | Hot-Warm (Target Body Site) T | n | Hot-Warm x Target – Other Body Site T | n | Imagine-Rest (Target Body Site) T | n | Imagine-Warm (Target Body Site) T | n | Imagine-Warm x Target – Other Body Site T | n |
|---|---|---|---|---|---|---|---|---|---|---|---|---|---|---|---|---|---|
| Shen Midb Lc | Midbrain L | | -8.5 | -18.5 | -18.5 | 3.98** | 7 | 4.57** | 9 | 3.02† | 4 | -2.18 | 0 | 0.73 | 0 | 0.16 | 0 |
| Shen Midb Lrd | Midbrain L | | -12.5 | -20.5 | -6.5 | 2.64* | 5 | 3.12* | 6 | 0.73 | 1 | -2.3 | 0 | -0.29 | 0 | -0.7 | 0 |
| Shen Midb Rcd | Midbrain R | | 11.5 | -20.5 | -16.5 | 2.69* | 5 | 2.8* | 5 | 2.51 | 4 | -1.34 | 0 | 1.3 | 2 | 1.26 | 0 |
| Shen Midb Rrd | Midbrain R | | 13.5 | -20.5 | -6.5 | 3.22* | 6 | 3.48* | 6 | 2.93 | 5 | -2.66 | 0 | -0.26 | 0 | 0.15 | 0 |
| Shen Pons Lcd | Pons L | RVM | -8.5 | -32.5 | -40.5 | 1.82 | 4 | 2.38† | 7 | 0.87 | 2 | -0.98 | 0 | 0.7 | 1 | 0.45 | 0 |
| Shen Pons Lrd | Pons L | | -8.5 | -36.5 | -28.5 | 3* | 7 | 3.03* | 7 | 2.21 | 4 | -1.19 | 0 | 1.05 | 1 | -0.84 | 0 |
| Shen Pons Lv | Pons L | | -6.5 | -20.5 | -34.5 | 2† | 4 | 2.51* | 7 | 1.56 | 1 | -2.06 | 0 | -0.35 | 0 | -0.82 | 0 |
| Shen Pons Rcd | Pons R | RVM | 9.5 | -34.5 | -36.5 | 1.02 | 2 | 0.62 | 2 | -1.16 | 0 | -1.43 | 0 | -0.81 | 1 | -1.28 | 0 |
| Shen Pons Rcv | Pons R | RVM | 5.5 | -24.5 | -42.5 | 1.47 | 3 | 1.6 | 3 | 0.56 | 0 | -0.69 | 0 | 0.28 | 0 | 0.19 | 0 |
| Shen Pons Rrv | Pons R | | 9.5 | -20.5 | -30.5 | 1.88 | 4 | 1.79 | 4 | 0.72 | 1 | -1.46 | 0 | -0.52 | 0 | 0.06 | 0 |
| SII+ L | somatomotor operculum L | S2/dpIns | -46.5 | -24.5 | 21.5 | 11.08*** | 9 | 10.11*** | 9 | 4.87** | 8 | -6.28* | 0 | -0.34 | 1 | -1.17 | 0 |
| SII+ R | somatomotor operculum R | S2/dpIns | 45.5 | -20.5 | 21.5 | | | | | | | | | | | | |
| SMA L | somatomotor paracentral lobule L | | -12.5 | -2.5 | 65.5 | 5.71** | 9 | 5.83** | 9 | 5.64*** | 9 | -0.44 | 1 | 2.22 | 4 | -0.42 | 0 |
| SMA R | somatomotor paracentral lobule R | | 15.5 | -2.5 | 65.5 | | | | | | | | | | | | |
| STH L | Midbrain L | | -10.5 | -10.5 | -6.5 | 0.94 | 0 | 1.54 | 1 | -0.53 | 0 | -0.37 | 0 | 0.59 | 0 | -0.66 | 0 |
| STH R | Midbrain R | | 11.5 | -10.5 | -6.5 | | | | | | | | | | | | |
| striate ctx L | visual early L | | -10.5 | -84.5 | 1.5 | 3.02* | 5 | 3.31* | 5 | 1.51 | 3 | -2.24 | 0 | 1.33 | 2 | -0.58 | 0 |
| striate ctx R | visual early R | | 13.5 | -80.5 | 3.5 | | | | | | | | | | | | |
| STV L | parietal TPOJ L | | -60.5 | -50.5 | 17.5 | 4.84** | 8 | 4.74** | 8 | 1.97 | 4 | -0.75 | 0 | 2.15 | 4 | 0.55 | 0 |
| STV R | parietal TPOJ R | | 59.5 | -42.5 | 19.5 | | | | | | | | | | | | |
| Subiculum L | Hippocampal Formation L | | -20.5 | -20.5 | -18.5 | 3.76* | 6 | 3.86* | 6 | 1.73 | 2 | -1.86 | 0 | 1.18 | 2 | 0.27 | 0 |
| Subiculum R | Hippocampal Formation R | | 21.5 | -20.5 | -18.5 | | | | | | | | | | | | |
| superior IPL L | parietal inferior lobule L | | -34.5 | -58.5 | 39.5 | 4.2** | 7 | 4.46** | 7 | 1.71 | 2 | -2.92 | 0 | 0.78 | 2 | -1.26 | 0 |
| superior IPL R | parietal inferior lobule R | | 39.5 | -54.5 | 43.5 | | | | | | | | | | | | |
| Superior lateral BA8 L | cingulate dlPFC L | | -16.5 | 25.5 | 53.5 | 2.73* | 5 | 2.74* | 5 | 0.68 | 3 | 0.02 | 2 | 1.41 | 3 | -0.87 | 1 |
| Superior lateral BA8 R | cingulate dlPFC R | | 17.5 | 27.5 | 55.5 | | | | | | | | | | | | |
| superior premotor L | somatomotor premotor L | | -30.5 | -6.5 | 57.5 | 5.58** | 9 | 5.91** | 9 | 2.99 | 6 | -1.66 | 0 | 1.36 | 2 | -0.94 | 0 |
| superior premotor R | somatomotor premotor R | | 33.5 | -4.5 | 55.5 | | | | | | | | | | | | |
| supplementary V3 L | visual dorsal L | | -22.5 | -86.5 | 21.5 | 2.66* | 5 | 2.59* | 5 | 1.12 | 1 | 0.17 | 1 | 1.78 | 3 | 0 | 0 |
| supplementary V3 R | visual dorsal R | | 25.5 | -82.5 | 25.5 | | | | | | | | | | | | |

| | | | MNI Coordinates | | | Hot-Rest (Target Body Site) | | Hot-Warm (Target Body Site) | | Hot-Warm x Target – Other Body Site | | Imagine-Rest (Target Body Site) | | Imagine-Warm (Target Body Site) | | Imagine-Warm x Target – Other Body Site | |
|---|---|---|---|---|---|---|---|---|---|---|---|---|---|---|---|---|---|
| Parcel | Region | Pain Region | X | Y | Z | T | n | T | n | T | n | T | n | T | n | T | n |
| Tectum L | Midbrain L | | -6.5 | -34.5 | -6.5 | 2.71* | 5 | 3.22* | 6 | 1.14 | 1 | -2.16 | 0 | 0.44 | 0 | -0.5 | 0 |
| Tectum R | Midbrain R | | 5.5 | -34.5 | -6.5 | | | | | | | | | | | | |
| temporal pole L | temporal lateral L | | -44.5 | 3.5 | -32.5 | 3.63* | 7 | 3.63* | 7 | 2.39 | 3 | -1.62 | 1 | 1.8 | 2 | 0.28 | 0 |
| temporal pole R | temporal lateral R | | 41.5 | 7.5 | -34.5 | | | | | | | | | | | | |
| Thal anterior L | Thal Anterior L | | -4.5 | -4.5 | 9.5 | 2.75* | 3 | 2.7* | 3 | 1.97 | 3 | -1.7 | 0 | 1.1 | 2 | 0.33 | 0 |
| Thal anterior R | Thal Anterior R | | 3.5 | -2.5 | 9.5 | | | | | | | | | | | | |
| Thal dorsal L | Thal Lateral L | | -8.5 | -16.5 | 17.5 | 2.5* | 3 | 2.63* | 4 | 2.33 | 2 | -0.73 | 0 | 1.6 | 1 | 1 | 0 |
| Thal dorsal R | Thal Lateral R | | 7.5 | -14.5 | 17.5 | | | | | | | | | | | | |
| Thal Intralaminar-anterodorsal L | Thal Intralaminar L | Thal IL | -0.5 | -10.5 | 13.5 | 2.69* | 5 | 2.45* | 3 | 0.51 | 1 | -0.58 | 0 | 0.69 | 2 | -0.97 | 0 |
| Thal Intralaminar-anterodorsal R | Thal Intralaminar R | Thal IL | 3.5 | -12.5 | 13.5 | | | | | | | | | | | | |
| Thal Intralaminar-antoventral L | Thal Intralaminar L | Thal IL | -2.5 | -10.5 | -0.5 | 2.52* | 5 | 3.15* | 6 | 2.64 | 5 | -1.09 | 0 | 1.05 | 1 | 1.12 | 0 |
| Thal Intralaminar-antoventral R | Thal Intralaminar R | Thal IL | 3.5 | -8.5 | -0.5 | | | | | | | | | | | | |
| Thal Intralaminar-posterior L | Thal Intralaminar L | Thal IL | -10.5 | -20.5 | 1.5 | 2.68* | 4 | 2.86* | 5 | 0.38 | 1 | -1.89 | 0 | 0.29 | 1 | -1.05 | 0 |
| Thal Intralaminar-posterior R | Thal Intralaminar R | Thal IL | 9.5 | -18.5 | 1.5 | | | | | | | | | | | | |
| Thal Lateral-caudal L | Thal Ventral L | Thal VPLM | -18.5 | -20.5 | 5.5 | 2.75* | 5 | 2.73* | 5 | 1.05 | 1 | -1.67 | 0 | 1.23 | 1 | -0.72 | 0 |
| Thal Lateral-caudal R | Thal Ventral R | Thal VPLM | 17.5 | -18.5 | 5.5 | | | | | | | | | | | | |
| Thal Lateral-rostral L | Thal Ventral L | | -10.5 | -8.5 | 7.5 | 2.53* | 4 | 2.42† | 3 | 0.64 | 2 | -3.54* | 0 | 0.53 | 1 | -2.44 | 0 |
| Thal Lateral-rostral R | Thal Ventral R | | 11.5 | -8.5 | 7.5 | | | | | | | | | | | | |
| Thal LGN L | Thal Posterior L | | -24.5 | -26.5 | -2.5 | 3.07* | 7 | 3.86** | 7 | 2.84† | 7 | -1.89 | 0 | 0.25 | 0 | 0.82 | 0 |
| Thal LGN R | Thal Posterior R | | 23.5 | -26.5 | -2.5 | | | | | | | | | | | | |
| Thal Medial L | Thal Medial L | Thal MD | -4.5 | -14.5 | 5.5 | 3.92* | 7 | 3.84* | 7 | 1.49 | 1 | -2.77† | 0 | 0.94 | 2 | -2.1 | 0 |
| Thal Medial R | Thal Medial R | Thal MD | 5.5 | -14.5 | 5.5 | | | | | | | | | | | | |
| Thal MGN L | Thal Posterior L | | -16.5 | -28.5 | -2.5 | 1.32 | 2 | 1.75 | 2 | 1.4 | 1 | -1.46 | 0 | 0.47 | 1 | 0.41 | 0 |
| Thal MGN R | Thal Posterior R | | 15.5 | -26.5 | -2.5 | | | | | | | | | | | | |
| Thal Pulvinar L | Thal Posterior L | | -12.5 | -28.5 | 9.5 | 3.71* | 6 | 3.65* | 6 | 2.9 | 5 | -2.54 | 0 | 1.47 | 2 | -0.6 | 0 |
| Thal Pulvinar R | Thal Posterior R | | 13.5 | -26.5 | 9.5 | | | | | | | | | | | | |

| Parcel | Region | Pain Region | MNI Coordinates X | Y | Z | Hot-Rest (Target Body Site) T | n | Hot-Warm (Target Body Site) T | n | Hot-Warm x Target – Other Body Site T | n | Imagine-Rest (Target Body Site) T | n | Imagine-Warm (Target Body Site) T | n | Imagine-Warm x Target – Other Body Site T | n |
|---|---|---|---|---|---|---|---|---|---|---|---|---|---|---|---|---|---|
| TPOJ L | parietal TPOJ L | | -50.5 | -56.5 | 11.5 | 3.95* | 5 | 3.81* | 6 | 1.33 | 2 | -1.36 | 0 | 2.07 | 3 | -0.53 | 0 |
| TPOJ R | parietal TPOJ R | | 51.5 | -50.5 | 11.5 | | | | | | | | | | | | |
| V6 L | visual dorsal L | | -16.5 | -80.5 | 33.5 | 2.83* | 5 | 2.69* | 5 | 1.17 | 3 | -1.01 | 1 | 1.46 | 2 | -0.49 | 0 |
| V6 R | visual dorsal R | | 19.5 | -78.5 | 35.5 | | | | | | | | | | | | |
| V7 L | visual dorsal L | | -24.5 | -84.5 | 31.5 | 2.82* | 5 | 2.8* | 5 | 1.45 | 2 | -0.09 | 0 | 1.54 | 3 | 0.22 | 0 |
| V7 R | visual dorsal R | | 29.5 | -82.5 | 33.5 | | | | | | | | | | | | |
| ventral intraparietal L | parietal superior lobule L | | -34.5 | -44.5 | 43.5 | 7*** | 9 | 7.36*** | 9 | 1.92 | 3 | -3.46† | 0 | 0.63 | 0 | -1.2 | 0 |
| ventral intraparietal R | parietal superior lobule R | | 35.5 | -40.5 | 43.5 | | | | | | | | | | | | |
| vmPFC L | cingulate ACC mPFC L | | -8.5 | 41.5 | -2.5 | 3.11* | 6 | 3.52* | 6 | 0.6 | 2 | -0.76 | 1 | 1.16 | 1 | -0.39 | 0 |
| vmPFC R | cingulate ACC mPFC R | | 7.5 | 41.5 | -2.5 | | | | | | | | | | | | |
| vMT+ L | visual MT+ L | | -46.5 | -70.5 | 1.5 | 3.73* | 5 | 3.4* | 5 | 2.09 | 3 | 0.24 | 1 | 2.55 | 3 | 0.47 | 0 |
| vMT+ R | visual MT+ R | | 47.5 | -66.5 | -0.5 | | | | | | | | | | | | |
| VMV L | visual ventral L | | -24.5 | -56.5 | -6.5 | 3.36* | 5 | 3.38* | 5 | 1.7 | 4 | -0.64 | 0 | 2.17 | 5 | 0.14 | 0 |
| VMV R | visual ventral R | | 25.5 | -54.5 | -6.5 | | | | | | | | | | | | |
| VStriatum L | VStriatum L | | -8.5 | 11.5 | -6.5 | 3.03* | 5 | 2.93* | 5 | 1.49 | 2 | -1.32 | 0 | 1.36 | 3 | -0.02 | 0 |
| VStriatum R | VStriatum R | | 7.5 | 11.5 | -4.5 | | | | | | | | | | | | |
| VTA PBP SN L | Midbrain L | | -8.5 | -16.5 | -12.5 | 2.07† | 5 | 2.16† | 5 | 1.22 | 0 | -1.63 | 0 | -0.35 | 0 | -0.57 | 0 |
| VTA PBP SN R | Midbrain R | | 9.5 | -16.5 | -12.5 | | | | | | | | | | | | |
| VVC L | visual ventral L | | -30.5 | -60.5 | -14.5 | 3.31* | 5 | 3.26* | 5 | 1.25 | 3 | 1.18 | 2 | 2.96 | 4 | -0.09 | 0 |
| VVC R | visual ventral R | | 29.5 | -54.5 | -16.5 | | | | | | | | | | | | |

**Notes.** Left (L) and Right (R) hemispheric regions are listed separately to define MNI XYZ coordinates separately, but were tested as a single region for pattern expression across body sites. Pain region column corresponds to voxel overlap within a-priori regions of interest in regions shown in Supplementary Table 3. P-values from all tests were FDR-corrected across 131 regional tests. n: Number of subjects with significant pattern expression post-FDR correction across 131 regions. aMCC MPFC: anterior midcingulate cortex/medial prefrontal cortex, aIns: anterior insula, mIns: medial insula, dpIns: dorsoposterior insula, S1: primary somatosensory cortex, S2: secondary somatosensory cortex, Thal: thalamus, VPLM: medial ventral posterolateral, IL: intralaminar, MD: mediodorsal, PAG: periaqueductal gray, RVM: rostral ventral medulla. †$p < .10$, *$p < 0.05$, **$p < 0.01$ and ***$p < .001$.

### Supplementary Table S5: Representational similarity analysis by region

| | Hot > Warm | | | Hot > Imagine | | | Imagine > Warm | | | (Hot, Imagine) > (Hot, Warm) | | | (Hot, Imagine) > (Imagine, Warm) | | | Target Body Site > Other Body Site | | |
|---|---|---|---|---|---|---|---|---|---|---|---|---|---|---|---|---|---|---|
| ROI | T | Hot > Warm (n) | Hot < Warm (n) | T | Hot > Imagine (n) | Hot < Imagine (n) | T | Imagine > Warm (n) | Imagine < Warm (n) | T | (Hot,Imagine) > (Hot,Warm) (n) | (Hot,Imagine) < (Hot,Warm) (n) | T | (Hot,Imagine) > (Imagine,Warm) (n) | (Hot,Imagine) < (Imagine,Warm) (n) | T | Target > Other (n) | Target < Other (n) |
| 55b | 4.08** | 9 | 0 | -1.43 | 4 | 5 | 3.75* | 9 | 0 | 2.90 | 6 | 3 | 3.61* | 8 | 1 | 0.47 | 6 | 3 |
| aCAU | 2.68* | 7 | 2 | -0.58 | 5 | 4 | 2.85* | 9 | 0 | 3.08 | 9 | 0 | 1.51 | 8 | 1 | 0.14 | 6 | 3 |
| AMY BL CeM | 3.58* | 9 | 0 | 1.87 | 7 | 2 | 2.83* | 8 | 1 | 0.70 | 4 | 5 | 0.57 | 4 | 5 | 0.80 | 5 | 4 |
| AMY La | 1.72 | 6 | 3 | -0.83 | 5 | 4 | 2.59* | 7 | 2 | 0.61 | 5 | 4 | 0.07 | 4 | 5 | -1.58 | 3 | 6 |
| anterior AAC | 3.03* | 9 | 0 | 0.24 | 6 | 3 | 3.31* | 9 | 0 | 2.95 | 8 | 1 | 3.38* | 9 | 0 | -1.03 | 3 | 6 |
| anterior agranular insula | 4.19** | 9 | 0 | 2.59 | 8 | 1 | 3.75* | 9 | 0 | 1.86 | 7 | 2 | 2.65* | 7 | 2 | 3.68* | 8 | 1 |
| anterior BA7 | 4.51** | 9 | 0 | 2.69 | 7 | 2 | 4.51* | 9 | 0 | 2.03 | 6 | 3 | 4.94* | 8 | 1 | 3.80* | 9 | 0 |
| anterior IPL | 6.51** | 9 | 0 | 4.80* | 9 | 0 | 3.46* | 8 | 1 | 0.79 | 6 | 3 | 4.50* | 8 | 1 | 0.81 | 6 | 3 |
| anterior operculum | 5.45** | 9 | 0 | 3.39 | 9 | 0 | 2.98* | 8 | 1 | 1.21 | 6 | 3 | 2.73* | 7 | 2 | 0.79 | 6 | 3 |
| anterior precuneus | 3.55* | 9 | 0 | 0.82 | 5 | 4 | 4.16* | 8 | 1 | 1.27 | 4 | 5 | 2.29 | 6 | 3 | -0.30 | 5 | 4 |
| aPUT | 2.83* | 7 | 2 | 0.51 | 5 | 4 | 2.59* | 8 | 1 | 1.67 | 6 | 3 | 1.41 | 6 | 3 | 0.68 | 4 | 5 |
| auditory transition area | 5.37** | 9 | 0 | 3.49 | 8 | 1 | 3.31* | 7 | 2 | 2.01 | 6 | 3 | 3.46* | 9 | 0 | 3.96* | 9 | 0 |
| auditory+ | 3.70* | 9 | 0 | 2.06 | 8 | 1 | 2.50* | 8 | 1 | 1.18 | 5 | 4 | 1.62 | 6 | 3 | 7.13** | 9 | 0 |
| BA10 | 2.31* | 8 | 1 | 0.75 | 6 | 3 | 2.98* | 8 | 1 | 3.17 | 8 | 1 | 2.07 | 6 | 3 | 1.41 | 6 | 3 |
| BA11 | 3.71* | 9 | 0 | 1.39 | 6 | 3 | 4.22* | 9 | 0 | 3.33 | 9 | 0 | 2.90* | 8 | 1 | 0.23 | 5 | 4 |
| BA13 | 3.26* | 8 | 1 | 0.68 | 6 | 3 | 2.28* | 6 | 3 | 1.24 | 6 | 3 | 0.56 | 5 | 4 | 0.64 | 6 | 3 |
| BA44 | 3.67* | 9 | 0 | 1.65 | 6 | 3 | 3.17* | 9 | 0 | 2.05 | 8 | 1 | 2.45 | 7 | 2 | -0.46 | 4 | 5 |
| BA45 | 3.54* | 9 | 0 | 1.05 | 6 | 3 | 3.37* | 9 | 0 | 2.53 | 7 | 2 | 3.24* | 7 | 2 | 0.97 | 5 | 4 |
| BA46 | 3.96** | 9 | 0 | 0.48 | 5 | 4 | 3.86* | 9 | 0 | 3.15 | 8 | 1 | 2.95* | 7 | 2 | 1.35 | 6 | 3 |
| BA47 | 3.26* | 9 | 0 | 0.67 | 5 | 4 | 2.84* | 9 | 0 | 1.97 | 5 | 4 | 2.81* | 6 | 3 | 1.55 | 7 | 2 |
| BA5 | 5.41** | 9 | 0 | 1.82 | 7 | 2 | 3.74* | 9 | 0 | 0.71 | 4 | 5 | 1.52 | 7 | 2 | 3.09 | 8 | 1 |
| BA9 | 2.60* | 9 | 0 | 0.34 | 4 | 5 | 3.56* | 9 | 0 | 2.77 | 8 | 1 | 2.73* | 8 | 1 | -0.62 | 4 | 5 |
| Cblm CrusI CrusII VIIIb | 3.49* | 9 | 0 | -0.84 | 5 | 4 | 3.17* | 8 | 1 | 3.01 | 6 | 3 | 3.10* | 8 | 1 | 1.36 | 6 | 3 |
| Cblm I IV V VI | 3.12* | 9 | 0 | 0.92 | 6 | 3 | 3.67* | 9 | 0 | 2.51 | 8 | 1 | 2.63* | 9 | 0 | 2.60 | 7 | 2 |
| Cblm vermis | 2.26* | 9 | 0 | 0.34 | 4 | 5 | 3.22* | 8 | 1 | 2.30 | 7 | 2 | 2.12 | 7 | 2 | 0.07 | 4 | 5 |
| Cblm VIIIa VIIb IX X | 4.30** | 9 | 0 | 0.89 | 6 | 3 | 3.42* | 8 | 1 | 2.72 | 7 | 2 | 3.32* | 8 | 1 | 1.93 | 6 | 3 |
| Cholinergic nuclei | 1.52 | 6 | 3 | 0.33 | 4 | 5 | 1.83 | 7 | 2 | 2.06 | 7 | 2 | 1.23 | 6 | 3 | -0.19 | 3 | 6 |
| cingulate motor area | 4.24** | 9 | 0 | 2.47 | 7 | 2 | 2.99* | 8 | 1 | 1.26 | 4 | 5 | 1.80 | 7 | 2 | 2.91 | 7 | 2 |
| Cranial nuclei | 2.15* | 7 | 2 | 0.13 | 5 | 4 | 2.11* | 7 | 2 | 0.94 | 5 | 4 | 1.31 | 5 | 4 | 1.04 | 6 | 3 |
| dmPFC | 2.24* | 9 | 0 | 0.21 | 5 | 4 | 3.96* | 9 | 0 | 2.97 | 7 | 2 | 2.35 | 6 | 3 | -1.07 | 4 | 5 |
| dMT+ | 3.02* | 8 | 1 | 0.18 | 5 | 4 | 3.71* | 8 | 1 | 3.48 | 9 | 0 | 2.27 | 8 | 1 | -0.25 | 5 | 4 |
| dorsal intraparietal | 3.53* | 9 | 0 | 0.34 | 4 | 5 | 4.13* | 9 | 0 | 2.59 | 7 | 2 | 4.43* | 9 | 0 | 0.65 | 4 | 5 |
| early auditory ctx a1 | 4.01** | 9 | 0 | 2.26 | 7 | 2 | 2.43* | 6 | 3 | 0.07 | 4 | 5 | 1.11 | 6 | 3 | 4.82* | 9 | 0 |
| extrastriate | 2.60* | 9 | 0 | -3.33 | 0 | 9 | 3.89* | 9 | 0 | 2.12 | 8 | 1 | 1.70 | 6 | 3 | 1.40 | 5 | 4 |
| FFC | 2.15* | 7 | 2 | -1.34 | 1 | 8 | 5.49* | 9 | 0 | 1.84 | 6 | 3 | 1.99 | 6 | 3 | 1.74 | 6 | 3 |

| | Hot > Warm | | | Hot > Imagine | | | Imagine > Warm | | | (Hot, Imagine) > (Hot, Warm) | | | (Hot, Imagine) > (Imagine, Warm) | | | Target Body Site > Other Body Site | | |
|---|---|---|---|---|---|---|---|---|---|---|---|---|---|---|---|---|---|---|
| ROI | T | Hot > Warm (n) | Hot < Warm (n) | T | Hot > Imagine (n) | Hot < Imagine (n) | T | Imagine > Warm (n) | Imagine < Warm (n) | T | (Hot,Imagine) > (Hot,Warm) (n) | (Hot,Imagine) < (Hot,Warm) (n) | T | (Hot,Imagine) > (Imagine,Warm) (n) | (Hot,Imagine) < (Imagine,Warm) (n) | T | Target > Other (n) | Target < Other (n) |
| FOP1-BA43 | 8.85*** | 9 | 0 | 7.17** | 9 | 0 | 3.19* | 8 | 1 | 0.55 | 5 | 4 | 4.10* | 8 | 1 | 3.56* | 8 | 1 |
| GPe | 1.99* | 6 | 3 | -0.26 | 5 | 4 | 2.16* | 8 | 1 | 2.13 | 6 | 3 | 2.02 | 7 | 2 | 2.37 | 7 | 2 |
| GPi | 1.89 | 5 | 4 | 2.11 | 7 | 2 | 1.55 | 5 | 4 | 1.32 | 5 | 4 | 1.38 | 6 | 3 | 1.72 | 7 | 2 |
| Hipp | 2.10* | 9 | 0 | 1.47 | 7 | 2 | 3.41* | 7 | 2 | 0.03 | 3 | 6 | -0.12 | 5 | 4 | 0.27 | 5 | 4 |
| hippocampal formation | 2.74* | 8 | 1 | 0.07 | 4 | 5 | 4.23* | 8 | 1 | 2.59 | 7 | 2 | 1.61 | 6 | 3 | 0.74 | 4 | 5 |
| hypothalamus | 2.48* | 8 | 1 | 1.57 | 6 | 3 | 2.58* | 7 | 2 | 1.89 | 7 | 2 | 1.07 | 6 | 3 | 1.85 | 7 | 2 |
| IFJ | 2.86* | 8 | 1 | -1.03 | 5 | 4 | 3.31* | 8 | 1 | 1.83 | 7 | 2 | 2.18 | 7 | 2 | -0.80 | 4 | 5 |
| IFS | 4.64** | 9 | 0 | -0.04 | 5 | 4 | 3.16* | 9 | 0 | 2.34 | 8 | 1 | 3.05* | 8 | 1 | -0.53 | 4 | 5 |
| inferior dysgranular insula | 5.88** | 9 | 0 | 5.06* | 8 | 1 | 3.37* | 8 | 1 | -0.46 | 3 | 6 | 1.36 | 7 | 2 | 3.39 | 8 | 1 |
| inferior IPL | 3.09* | 9 | 0 | 0.66 | 6 | 3 | 3.43* | 9 | 0 | 3.22 | 7 | 2 | 3.39* | 8 | 1 | -0.30 | 3 | 6 |
| Inferior lateral BA8 | 3.13* | 8 | 1 | -0.88 | 3 | 6 | 3.32* | 9 | 0 | 3.24 | 7 | 2 | 2.34 | 6 | 3 | -0.37 | 4 | 5 |
| inferior premotor | 4.45** | 9 | 0 | 2.13 | 7 | 2 | 3.70* | 8 | 1 | 2.15 | 7 | 2 | 3.03* | 8 | 1 | 0.42 | 6 | 3 |
| inferior temporal sulcus | 2.29* | 9 | 0 | 0.36 | 5 | 4 | 4.50* | 8 | 1 | 3.09 | 6 | 3 | 1.62 | 5 | 4 | 1.30 | 5 | 4 |
| infralimbic | 2.16* | 8 | 1 | 0.83 | 6 | 3 | 2.22* | 6 | 3 | 2.62 | 7 | 2 | 1.44 | 6 | 3 | 0.76 | 6 | 3 |
| ISP1 | 3.66* | 9 | 0 | -0.31 | 4 | 5 | 3.98* | 9 | 0 | 1.67 | 7 | 2 | 2.21 | 7 | 2 | 1.82 | 6 | 3 |
| L PnO PnC | 1.01 | 6 | 3 | 0.57 | 6 | 3 | 0.52 | 4 | 5 | 0.50 | 4 | 5 | 0.92 | 4 | 5 | 1.49 | 6 | 3 |
| LC+ | 0.31 | 7 | 2 | -0.70 | 3 | 6 | 0.86 | 6 | 3 | 0.85 | 4 | 5 | 0.56 | 4 | 5 | -0.50 | 3 | 6 |
| LO | 3.08* | 8 | 1 | -2.44 | 2 | 7 | 5.39* | 9 | 0 | 3.36 | 8 | 1 | 3.16* | 9 | 0 | 2.83 | 8 | 1 |
| M1 | 4.17** | 9 | 0 | 0.87 | 7 | 2 | 3.02* | 9 | 0 | 0.94 | 5 | 4 | 1.41 | 7 | 2 | 2.86 | 8 | 1 |
| Medullary Raphe (Serotonergic) | 0.19 | 1 | 8 | -0.36 | 3 | 6 | 0.86 | 4 | 5 | -0.84 | 4 | 5 | -0.14 | 5 | 4 | 0.33 | 5 | 4 |
| Medullary reticular formation | 2.65* | 9 | 0 | 1.62 | 7 | 2 | 1.34 | 4 | 5 | 1.88 | 6 | 3 | 2.58 | 6 | 3 | 0.53 | 3 | 6 |
| midcingulate | 4.25** | 9 | 0 | 0.93 | 7 | 2 | 2.72* | 9 | 0 | 1.95 | 6 | 3 | 2.50 | 8 | 1 | 1.93 | 8 | 1 |
| middle IPL | 3.44* | 8 | 1 | 0.24 | 5 | 4 | 3.22* | 9 | 0 | 1.92 | 6 | 3 | 2.84* | 6 | 3 | 0.54 | 4 | 5 |
| middle temporal gyrus | 3.52* | 9 | 0 | 0.18 | 4 | 5 | 3.26* | 9 | 0 | 1.48 | 6 | 3 | 2.74* | 7 | 2 | 0.70 | 5 | 4 |
| MIP | 3.43* | 9 | 0 | 0.48 | 6 | 3 | 3.41* | 9 | 0 | 1.55 | 6 | 3 | 2.80* | 9 | 0 | -0.56 | 4 | 5 |
| OFC | 2.30* | 7 | 2 | 0.66 | 7 | 2 | 3.12* | 8 | 1 | 1.59 | 7 | 2 | 0.87 | 5 | 4 | 0.30 | 4 | 5 |
| Olivary complex | 1.74 | 7 | 2 | 1.17 | 6 | 3 | 2.57* | 8 | 1 | 1.33 | 6 | 3 | 1.24 | 5 | 4 | -0.64 | 5 | 4 |
| OP4 | 10.35*** | 9 | 0 | 6.99** | 9 | 0 | 1.15 | 6 | 3 | -1.59 | 3 | 6 | 3.31* | 8 | 1 | 4.46* | 8 | 1 |
| PAG | 1.47 | 7 | 2 | 0.29 | 6 | 3 | 1.69 | 7 | 2 | -0.34 | 3 | 6 | -0.07 | 4 | 5 | -1.59 | 1 | 8 |
| Parabrachial nuclei | 2.42* | 8 | 1 | 1.47 | 6 | 3 | 2.58* | 7 | 2 | 1.29 | 6 | 3 | 1.12 | 6 | 3 | -0.96 | 4 | 5 |
| parietal occipital sulcus | 3.43* | 9 | 0 | 0.90 | 6 | 3 | 3.64* | 9 | 0 | 2.13 | 8 | 1 | 2.31 | 7 | 2 | 0.75 | 5 | 4 |
| pCAU | 2.30* | 8 | 1 | -0.19 | 4 | 5 | 3.58* | 8 | 1 | 2.11 | 8 | 1 | 1.89 | 8 | 1 | -0.17 | 5 | 4 |
| PCV | 4.57** | 9 | 0 | -0.99 | 4 | 5 | 3.25* | 9 | 0 | 2.62 | 7 | 2 | 3.71* | 9 | 0 | -0.51 | 4 | 5 |

| ROI | Hot > Warm | | | Hot > Imagine | | | Imagine > Warm | | | (Hot, Imagine) > (Hot, Warm) | | | (Hot, Imagine) > (Imagine, Warm) | | | Target Body Site > Other Body Site | | |
|---|---|---|---|---|---|---|---|---|---|---|---|---|---|---|---|---|---|---|
| | T | Hot > Warm (n) | Hot < Warm (n) | T | Hot > Imagine (n) | Hot < Imagine (n) | T | Imagine > Warm (n) | Imagine < Warm (n) | T | (Hot,Imagine) > (Hot,Warm) (n) | (Hot,Imagine) < (Hot,Warm) (n) | T | (Hot,Imagine) > (Imagine,Warm) (n) | (Hot,Imagine) < (Imagine,Warm) (n) | T | Target > Other (n) | Target < Other (n) |
| PeEc | 3.59* | 9 | 0 | 0.99 | 7 | 2 | 2.70* | 9 | 0 | 1.20 | 5 | 4 | 0.49 | 5 | 4 | 2.87 | 7 | 2 |
| PH RPH | 2.52* | 9 | 0 | 0.46 | 6 | 3 | 4.65* | 9 | 0 | 2.12 | 7 | 2 | 2.63* | 9 | 0 | -0.25 | 2 | 7 |
| PHA | 2.55* | 6 | 3 | 0.05 | 4 | 5 | 3.86* | 9 | 0 | 2.46 | 6 | 3 | 2.47 | 8 | 1 | 0.57 | 5 | 4 |
| PIR | 5.70** | 9 | 0 | 2.92 | 8 | 1 | 2.98* | 8 | 1 | -0.36 | 3 | 6 | 1.50 | 5 | 4 | 0.47 | 6 | 3 |
| postcentral BA7 | 5.63** | 9 | 0 | 1.79 | 7 | 2 | 4.99* | 9 | 0 | 2.31 | 6 | 3 | 3.95* | 8 | 1 | 2.13 | 7 | 2 |
| posterior AAC | 2.84* | 9 | 0 | 0.45 | 5 | 4 | 3.13* | 8 | 1 | 2.35 | 7 | 2 | 3.20* | 9 | 0 | -0.63 | 4 | 5 |
| posterior BA7 | 3.05* | 8 | 1 | -0.44 | 4 | 5 | 3.17* | 8 | 1 | 1.73 | 5 | 4 | 2.33 | 7 | 2 | 0.31 | 5 | 4 |
| posterior granular insula | 5.95** | 9 | 0 | 5.14* | 9 | 0 | 2.31* | 7 | 2 | -1.58 | 3 | 6 | 2.28 | 7 | 2 | 1.18 | 5 | 4 |
| posterior IPL | 3.41* | 9 | 0 | -1.15 | 3 | 6 | 4.06* | 8 | 1 | 2.75 | 8 | 1 | 3.38* | 7 | 2 | 0.33 | 5 | 4 |
| posterior operculum | 13.24*** | 9 | 0 | 10.58*** | 9 | 0 | 2.86* | 6 | 3 | -0.05 | 4 | 5 | 3.30* | 7 | 2 | 4.49* | 8 | 1 |
| pPUT | 1.97* | 8 | 1 | -1.15 | 4 | 5 | 2.42* | 8 | 1 | 1.31 | 6 | 3 | 1.30 | 6 | 3 | 1.94 | 7 | 2 |
| precuneus proper | 3.09* | 9 | 0 | 0.37 | 5 | 4 | 3.00* | 8 | 1 | 1.96 | 5 | 4 | 2.06 | 7 | 2 | -0.47 | 2 | 7 |
| PSL | 4.58** | 9 | 0 | 3.42 | 7 | 2 | 3.63* | 8 | 1 | 1.72 | 6 | 3 | 3.57* | 8 | 1 | -0.32 | 4 | 5 |
| R PnO PnC | -0.11 | 4 | 5 | 0.55 | 4 | 5 | -0.76 | 3 | 6 | -0.17 | 3 | 6 | 0.37 | 3 | 6 | -0.39 | 4 | 5 |
| RN | 1.83 | 6 | 3 | -1.23 | 3 | 6 | 3.60* | 8 | 1 | 1.41 | 6 | 3 | 1.90 | 7 | 2 | -0.09 | 4 | 5 |
| Rostral Raphe (Serotonergic) | 0.98 | 6 | 3 | -0.06 | 5 | 4 | 1.47 | 7 | 2 | 3.57 | 8 | 1 | 2.11 | 6 | 3 | -1.14 | 2 | 7 |
| Rostral reticular formation | 2.34* | 7 | 2 | 0.86 | 7 | 2 | 1.25 | 6 | 3 | 0.31 | 3 | 6 | 0.84 | 5 | 4 | 0.08 | 6 | 3 |
| RSC | 3.90** | 9 | 0 | 0.69 | 6 | 3 | 2.77* | 9 | 0 | 1.76 | 6 | 3 | 2.53 | 8 | 1 | 0.06 | 5 | 4 |
| S1 | 4.33** | 9 | 0 | 2.19 | 8 | 1 | 3.30* | 9 | 0 | 1.27 | 6 | 3 | 2.04 | 7 | 2 | 3.71* | 8 | 1 |
| Shen Med | 2.45* | 9 | 0 | 1.59 | 6 | 3 | 1.91 | 6 | 3 | 1.74 | 6 | 3 | 2.55 | 6 | 3 | 1.10 | 6 | 3 |
| Shen Midb Lc | 2.44* | 7 | 2 | 1.14 | 5 | 4 | 2.91* | 9 | 0 | 1.98 | 8 | 1 | 0.84 | 5 | 4 | -0.25 | 5 | 4 |
| Shen Midb Lrd | 2.94* | 7 | 2 | -0.12 | 5 | 4 | 4.87* | 9 | 0 | 0.52 | 4 | 5 | -0.62 | 2 | 7 | 0.54 | 5 | 4 |
| Shen Midb Rcd | 2.61* | 9 | 0 | 0.83 | 6 | 3 | 4.10* | 9 | 0 | 1.55 | 6 | 3 | 0.72 | 4 | 5 | -1.93 | 3 | 6 |
| Shen Midb Rrd | 2.78* | 8 | 1 | 0.51 | 6 | 3 | 3.13* | 8 | 1 | -0.19 | 3 | 6 | -0.30 | 4 | 5 | 0.52 | 5 | 4 |
| Shen Pons Lcd | 2.83* | 8 | 1 | 1.12 | 7 | 2 | 4.06* | 9 | 0 | 0.90 | 6 | 3 | 0.80 | 4 | 5 | 0.13 | 3 | 6 |
| Shen Pons Lrd | 2.91* | 9 | 0 | 1.92 | 7 | 2 | 2.61* | 7 | 2 | 2.60 | 7 | 2 | 2.31 | 7 | 2 | -1.32 | 4 | 5 |
| Shen Pons Lv | 1.78 | 7 | 2 | 0.76 | 6 | 3 | 1.05 | 5 | 4 | -0.01 | 3 | 6 | -1.01 | 5 | 4 | -0.42 | 4 | 5 |
| Shen Pons Rcd | 2.46* | 8 | 1 | 1.40 | 6 | 3 | 2.55* | 8 | 1 | 1.36 | 6 | 3 | 0.89 | 6 | 3 | 0.70 | 5 | 4 |
| Shen Pons Rcv | 2.39* | 6 | 3 | 1.50 | 7 | 2 | 1.17 | 5 | 4 | -1.70 | 2 | 7 | 0.21 | 4 | 5 | -0.14 | 4 | 5 |
| Shen Pons Rrv | 1.11 | 5 | 4 | 0.01 | 5 | 4 | 1.82 | 6 | 3 | -0.09 | 3 | 6 | -0.13 | 4 | 5 | -1.07 | 3 | 6 |
| SII+ | 7.25** | 9 | 0 | 5.59** | 9 | 0 | 1.92 | 6 | 3 | -0.84 | 3 | 6 | 2.81* | 7 | 2 | 4.69* | 8 | 1 |
| SMA | 3.26* | 9 | 0 | -0.97 | 4 | 5 | 4.09* | 9 | 0 | 2.52 | 8 | 1 | 2.71* | 9 | 0 | 2.56 | 7 | 2 |
| STH | 1.71 | 6 | 3 | 0.61 | 6 | 3 | 1.43 | 6 | 3 | 1.58 | 6 | 3 | 0.82 | 7 | 2 | 0.06 | 5 | 4 |
| strate ctx | 2.85* | 9 | 0 | -2.26 | 4 | 5 | 4.39* | 9 | 0 | 2.70 | 7 | 2 | 1.51 | 6 | 3 | 0.82 | 5 | 4 |
| STV | 5.52** | 9 | 0 | 0.49 | 4 | 5 | 3.97* | 8 | 1 | 2.27 | 7 | 2 | 3.48* | 9 | 0 | -0.58 | 6 | 3 |

| | Hot > Warm | | | Hot > Imagine | | | Imagine > Warm | | | (Hot, Imagine) > (Hot, Warm) | | | (Hot, Imagine) > (Imagine, Warm) | | | Target Body Site > Other Body Site | | |
|---|---|---|---|---|---|---|---|---|---|---|---|---|---|---|---|---|---|---|
| ROI | T | Hot > Warm (n) | Hot < Warm (n) | T | Hot > Imagine (n) | Hot < Imagine (n) | T | Imagine > Warm (n) | Imagine < Warm (n) | T | (Hot,Imagine) > (Hot,Warm) (n) | (Hot,Imagine) < (Hot,Warm) (n) | T | (Hot,Imagine) > (Imagine,Warm) (n) | (Hot,Imagine) < (Imagine,Warm) (n) | T | Target > Other (n) | Target < Other (n) |
| Subiculum | 2.33* | 9 | 0 | 1.16 | 7 | 2 | 3.70* | 9 | 0 | 0.62 | 3 | 6 | -0.48 | 3 | 6 | -1.32 | 3 | 6 |
| superior IPL | 3.34* | 9 | 0 | 0.23 | 6 | 3 | 2.31* | 9 | 0 | 1.53 | 6 | 3 | 2.26 | 7 | 2 | 2.60 | 8 | 1 |
| Superior lateral BA8 | 2.81* | 9 | 0 | -0.34 | 3 | 6 | 3.69* | 9 | 0 | 3.15 | 8 | 1 | 2.87* | 8 | 1 | -0.72 | 3 | 6 |
| superior premotor | 4.37** | 9 | 0 | -0.27 | 3 | 6 | 3.51* | 9 | 0 | 1.67 | 6 | 3 | 1.94 | 8 | 1 | 2.58 | 7 | 2 |
| supplementary V3 | 2.80* | 9 | 0 | -2.99 | 2 | 7 | 3.83* | 9 | 0 | 2.80 | 9 | 0 | 2.38 | 7 | 2 | 2.76 | 8 | 1 |
| Tectum | 3.02* | 8 | 1 | 0.96 | 7 | 2 | 2.92* | 8 | 1 | 1.20 | 5 | 4 | 1.26 | 5 | 4 | 1.38 | 5 | 4 |
| temporal pole | 3.15* | 9 | 0 | 0.69 | 6 | 3 | 3.88* | 9 | 0 | 2.81 | 6 | 3 | 2.79* | 8 | 1 | 2.17 | 7 | 2 |
| Thal anterior | 1.92 | 7 | 2 | 1.14 | 7 | 2 | 2.62* | 7 | 2 | 1.81 | 8 | 1 | 1.46 | 6 | 3 | 0.11 | 4 | 5 |
| Thal dorsal | 2.29* | 8 | 1 | 1.09 | 7 | 2 | 3.91* | 7 | 2 | 1.51 | 7 | 2 | 0.87 | 6 | 3 | -1.70 | 2 | 7 |
| Thal Intralaminar-anterodorsal | 2.54* | 7 | 2 | 1.84 | 7 | 2 | 2.81* | 8 | 1 | 2.07 | 8 | 1 | 2.10 | 7 | 2 | -0.65 | 4 | 5 |
| Thal Intralaminar-antoventral | 0.93 | 6 | 3 | 1.07 | 6 | 3 | -0.06 | 3 | 6 | -0.10 | 4 | 5 | -0.40 | 4 | 5 | 1.92 | 7 | 2 |
| Thal Intralaminar-posterior | 2.68* | 7 | 2 | 1.93 | 6 | 3 | 2.25* | 7 | 2 | -0.32 | 4 | 5 | -0.15 | 4 | 5 | 2.84 | 7 | 2 |
| Thal Lateral-caudal | 2.52* | 9 | 0 | 2.21 | 7 | 2 | 3.02* | 8 | 1 | 1.70 | 6 | 3 | 1.85 | 8 | 1 | 0.34 | 6 | 3 |
| Thal Lateral-rostral | 2.20* | 9 | 0 | 1.77 | 7 | 2 | 3.16* | 7 | 2 | 1.43 | 8 | 1 | 1.63 | 5 | 4 | 0.01 | 4 | 5 |
| Thal LGN | 2.54* | 7 | 2 | 1.38 | 7 | 2 | 2.87* | 8 | 1 | -1.86 | 2 | 7 | -1.68 | 3 | 6 | -1.62 | 3 | 6 |
| Thal Medial | 3.13* | 8 | 1 | 2.00 | 7 | 2 | 3.73* | 8 | 1 | 1.42 | 7 | 2 | 1.55 | 7 | 2 | -2.50 | 2 | 7 |
| Thal MGN | 0.32 | 4 | 5 | -1.45 | 4 | 5 | 1.97* | 6 | 3 | 0.75 | 3 | 6 | -0.25 | 2 | 7 | 0.22 | 5 | 4 |
| Thal Pulvinar | 2.54* | 8 | 1 | 1.18 | 6 | 3 | 3.77* | 9 | 0 | 1.22 | 6 | 3 | 1.01 | 4 | 5 | -0.57 | 6 | 3 |
| TPOJ | 3.36* | 9 | 0 | 0.79 | 5 | 4 | 4.44* | 9 | 0 | 2.72 | 8 | 1 | 3.02* | 9 | 0 | 0.59 | 5 | 4 |
| V6 | 3.30* | 9 | 0 | -0.64 | 4 | 5 | 3.67* | 9 | 0 | 2.27 | 7 | 2 | 1.27 | 6 | 3 | 0.95 | 7 | 2 |
| V7 | 3.11* | 8 | 1 | -1.17 | 3 | 6 | 3.45* | 9 | 0 | 2.95 | 7 | 2 | 2.87* | 8 | 1 | 0.20 | 4 | 5 |
| ventral intraparietal | 4.35** | 9 | 0 | 1.46 | 6 | 3 | 3.32* | 7 | 2 | 1.25 | 7 | 2 | 3.38* | 8 | 1 | 2.53 | 8 | 1 |
| vmPFC | 2.40* | 9 | 0 | 0.62 | 4 | 5 | 3.07* | 8 | 1 | 2.65 | 7 | 2 | 2.10 | 7 | 2 | -0.66 | 4 | 5 |
| vMT+ | 3.53* | 9 | 0 | -0.31 | 5 | 4 | 4.68* | 9 | 0 | 3.78 | 9 | 0 | 3.47* | 8 | 1 | 1.94 | 7 | 2 |
| VMV | 1.79 | 7 | 2 | -2.35 | 3 | 6 | 2.86* | 9 | 0 | 2.11 | 8 | 1 | 2.20 | 7 | 2 | -0.35 | 5 | 4 |
| VStriatum | 2.95* | 7 | 2 | 1.61 | 7 | 2 | 3.39* | 9 | 0 | 2.14 | 6 | 3 | 1.15 | 5 | 4 | -0.67 | 4 | 5 |
| VTA PBP SN | 3.98** | 8 | 1 | 0.17 | 4 | 5 | 2.24* | 7 | 2 | 0.91 | 5 | 4 | 0.36 | 5 | 4 | 0.20 | 5 | 4 |
| VVC | 2.86* | 7 | 2 | -2.56 | 1 | 8 | 4.67* | 9 | 0 | 2.12 | 6 | 3 | 2.44 | 8 | 1 | 1.10 | 5 | 4 |

**Notes.** For each contrast, T is the signed one-sample t statistic (N = 9 participants) of the per-participant similarity difference (its sign indicates the direction of the effect). Asterisks denote significance in the named direction after false-discovery-rate (FDR) correction across the 131 parcels within that contrast (*q < .05, **q < .01, ***q < .001; one-tailed). The two count columns give the number of participants (of 9) whose similarity difference favored each direction. Rows are ordered alphabetically by region.
